# Advancing the understanding of the *Bombyx mori* genome through satellite DNA analysis: low strain differentiation and relationship with transposable elements

**DOI:** 10.1101/2025.04.21.649856

**Authors:** José M Rico-Porras, Pablo Mora, Teresa Palomeque, Pedro Lorite, Diogo C Cabral-de-Mello

**Author notes:** Correspondence (D.C.C.-d-M.). These authors had same contribution (José M Rico-Porras, Pablo Mora).

## Abstract

Despite the economic and scientific importance of *Bombyx mori*, its satellite DNA (satDNA) fraction remains poorly characterized. Here, we present the first comprehensive analysis of the *B. mori* satellitome, revealing its low abundance (∼0.76%), A+T enrichment, high monomer size variability, and predominantly dispersed chromosomal organization. Comparative analysis with *B. mandarina*, the wild ancestor of *B. mori*, demonstrated high satDNA conservation in abundance and sequence composition, despite speciation and domestication events. Divergence landscapes of satDNA sequences indicate predominantly ancient amplifications with few recent homogenization events. SatDNA families exhibited low strain differentiation across five *B. mori* strains, exemplified by the consistent presence and abundance of BmorSat01-575 and BmorSat29-84. The W chromosome was identified as a hotspot for satDNA accumulation, especially BmorSat24-753, which showed female-specific amplification and higher sequence homogeneity, suggesting functional or structural roles in W chromosome evolution. In contrast, the Z chromosome displayed similar satDNA content to autosomes, with two families being exclusive to this chromosome and conserved in both *Bombyx* species. Remarkably, some of *B. mori* satDNAs were found to be derived from transposable elements (TEs), supporting the hypothesis of TE-driven satDNA origin via multiple mechanisms, including tandemization of TE fragments. Our findings underscore the dynamic interplay between satDNAs and TEs in shaping genome architecture, the stability of the satellitome through domestication, and the sex chromosome-specific accumulation patterns. These insights lay the groundwork for future studies investigating the functional roles of satDNAs in genome regulation and chromatin dynamics in Lepidoptera.

## 1. Introduction

Bombycoidea comprises about 6092 species, making it one of the largest superfamilies of Lepidoptera. This superfamily includes the family Bombycidae, with 477 described species found in all zoogeographic regions, with a higher concentration in the Ethiopian and Oriental regions (Kitching et al., 2018). *Bombyx mori*, the domestic silkworm, is the most well-known Bombycidae species and is widely used as a model insect in various biological fields, including genetics and genomics. Moreover, this species is economically important due to silk production and is also used in the production of biomaterials with medical and industrial applications (Goldsmith et al., 2005; Ashraf and Qamar, 2023; Bitar et al., 2024; Rodríguez-Ortiz et al., 2024). The species was domesticated from its wild ancestor, *B. mandarina*, about 5000 years ago, and today more than 3000 silkworm strains are preserved worldwide (Underhill, 1997; Ryu et al., 2003; Xiang et al., 2018).

The silkworm genome is approximately 450 Mb in size and was first sequenced in 2004, making it the first Lepidoptera species to have its whole genome sequenced (Mita et al., 2004; Xia et al., 2004). Subsequent updates have been published (International Silkworm Genome Consortium, 2008; Kawamoto et al., 2019), with the most recent in 2022 presenting the pan-genome of the species (Tong et al., 2022). This latest update was based on the deeply re-sequencing of 1078 silkworm genomes and long-read assembly for 545 representatives, providing insights into its evolution and offering valuable information about alleles for breeding and functional genomics studies. Concerning repetitive DNA, the *de novo* assembly of the 545 genomes revealed that this DNA fraction averaged the 47% of the content, varying between 46-49% depending on the genome. Among them, the non-LTR transposons accounted for an average of 26%, making them the most abundant repeats. These numbers are similar to the previous genome assembly of the silkworm, which also estimated the abundance of other transposable elements (TEs). This included LTR elements, representing 2.33% of the genome, DNA elements at 2.67%, with the remaining 11.73% of interspersed repeats being unclassified. Simple repeats, together with low complexity sequences, corresponded to 0.96% in abundance (Kawamoto et al., 2019). However, no specific details about the tandem sequences, including the satellite DNAs (satDNAs), were studied in the genome assemblies.

Highly repetitive satDNA sequences constitute a substantial portion of eukaryotic genomes, primarily within heterochromatic regions (Garrido-Ramos, 2017). Their abundance varies among insects, ranging from 0.06% of the genome in *Diatraea plostineella*, Lepidoptera (Cabral-de-Mello et al., 2021) to more than 50%, as reported in *Triatoma delpontei*, Hemiptera (Mora et al., 2023). The repetitiveness nature of satDNAs and their tandem organization facilitate rapid evolution through mechanisms such as unequal crossing over and replication slippage. This process leads to variations in repeat number between individuals and species from a common ancestral set of sequences, as predicted by the library hypothesis (Fry and Salser, 1977; Walsh, 1987; Stephan, 1989). These variations can influence the evolutionary history of species. In addition, satDNAs can impact gene expression by altering chromatin structure or through the transcription of regulatory elements like non-coding RNAs (ncRNA) (Shatskikh et al., 2020; Liao et al., 2023). Thus, satDNAs are not merely structural and passive elements of genomes; they actively contribute to genome biology in terms of organization, dynamics, and evolution.

During genome assembly process, the high repetitiveness and similarity of satDNAs make them difficult to be assembled, often resulting in collapsed and fragmented scaffolds. This has caused the underrepresentation of satDNAs in genome assemblies for decades (Miga, 2015; Garrido-Ramos, 2017; Lower et al., 2018; Peona et al., 2021; Peona et al., 2018; 2022; Montiel et al., 2022; Mora et al., 2024). Therefore, complementary analyses of assembled genomes, short read sequencing, and cytogenetic mapping are important for deeper characterization of this genomic component. Analysis of short reads is useful for identifying and characterizing the evolutionary patterns of satDNAs. Nowadays, RepeatExplorer2/TAREAN (Novák et al., 2017) is frequently used in such studies, allowing the identification of the diversity of near to the complete satDNA library from a genome, known as the satellitome (Ruiz-Ruano et al., 2016). Cytogenetic mapping using Fluorescence *in situ* Hybridization (FISH) allows the physical mapping of satDNAs of interest on the chromosomes (Cabral-de-Mello and Marec, 2021), providing important information about their location. This can also be validated by *in silico* mapping on chromosome assemblies, using tools such as the CHRISMAPP pipeline (Rico-Porras et al., 2024), in cases where they are not completely collapsed.

Although the satDNA fraction has been characterized in some animals, including insects (Šatović-Vukšić and Plohl, 2023), it has not been properly identified and studied in some model organisms, including *B. mori* (Kawamoto et al., 2019; Tong et al., 2022). Among other Lepidoptera, despite progress in the genomic field (Triant et al., 2018), the analysis of the satDNA fraction is comparatively scarce. Only a few satDNAs have been studied, revealing generally low representation in their genomes and a common scattered chromosomal distribution (Věchtová et al., 2016; Dalíková et al., 2017a, 2023; Cabral-de-Mello et al., 2021; Haq et al., 2022). Given the economic and scientific significance of *B. mori*, as well as the crucial role of satDNAs in genome organization and evolution, there is a substantial gap in our knowledge regarding this genomic component in the species. Through comprehensive genomic and chromosomal analyses, we have identified and characterized the satellitome of *B. mori*, advancing our understanding of its genome organization and evolution. Moreover, we also characterized the satDNAs present in *B. mori* genome in the genome of its wild ancestor, *B. mandarina*, providing insights into the evolutionary aspects of satDNAs in closely related species.

## 2. Material and Methods

### 2.1. Animals, chromosomes and polyploid nucleus obtaining

Eggs from *B. mori* were provided by the company Gusilandia (https://gusilandia.com/). After eggs hatching the larvae were reared in the lab at 28°C and feed using mulberry leaves. The specimens belong to Europe-Local strain (Tong et al., 2022). Larvae between 4^th^ and 5^th^ instars were dissected for chromosome obtaining. Spread chromosome preparations were made as described previously (Mediouni et al., 2004; Šíchová et al., 2013). Testes and wing disks were dissected from males, hypotonized for 10 minutes (75mM KCl) and fixed in Carnoy’s solution (ethanol, chloroform, acetic acid, 6:3:1) for at least 15 min. For females wing disks and ovaries were dissected. Female wing disks also passed through hypotonization step, but the ovaries were directly fixed in Carnoy’s (ethanol, chloroform, acetic acid, 6:3:1). The fixed tissues were disaggregated in 50 µL of 45% glacial acetic acid by pipetting inside a microtube. The solution was dropped in a glass slide under heating of 45°C. The slides were observed under a phase contrast microscope and slides with sufficient quality were dehydrated in ethanol series (70, 80, and 100%, 30 s each) and stored at –20°C until use. The dissected larvae with the sex identified were stored in 100% ethanol and stored at –20 °C for genomic DNA (gDNA) extraction.

The Malpighian tubes from male and female larvae were used to obtain polyploid interphase nuclei (Marec and Traut, 1994). The tubules were dissected in physiological solution and fixed in Carnoy’s solution on a slide for about 1 min directly above a glass slide. The cells were stained with 1.25% lactic acetic orcein for 3–5 min and the slides mounted in the staining solution. The presence or absence of W chromatin (reviewed in Traut and Marec, 1996) was analyzed under a light microscope.

### 2.2. Short reads genomic data

The male and female gDNAs from larvae maintained in the lab from Europe-Local were extracted from their heads using the kit Wizard Genomic DNA Purification Kit (Promega, Madison, WI), following the manufacture’s protocol. About 2 µg of gDNA was used for genome sequencing based in Polymerase Chain Reaction (PCR) free library and using the Illumina HiSeq™ 2000 platform by Novogene (HK) Co., Ltd. (Hong Kong, China). It was obtained a total of 16,966,278 paired reads from males and 16,742,496 paired reads from females, both with 150 bp in length. The reads were deposited in the Sequence Read Achieve (SRA) under accession numbers **<u>XXXX</u>** (male) and **<u>XXXX</u>** (female).

Additionally, in order to have a broader information about the satDNAs in *B. mori* we also analyzed comparatively the genomes from individuals belonging to other geographic regions, including other four silkworm strains: CHN-1 strain improved in China (individuals BomO129 and BomO130), China-Local strain (individuals BomL2 and Bom L3), Japan-Local strain (individuals BomP23 and BomP24), and Tropic-Local strain (individuals BomL122 and BomL201). Details about these strains could be seen in Tong et al. (2022) and the raw reads are available in the SilkMeta, Silworm Pangenome Database (http://silkmeta.org.cn/home). Finally, to have a better understanding of satDNA library evolution, additionally, we included in the analysis the genomes from one male (DRX086504) and from one female (SRX17298094) of the wild silkworm *B. mandarina*, as it is the wild ancestor of *B. mori*.

### 2.3. Characterization of *Bombyx mori* satellitome

For identification of satDNAs in the genome of *B. mori* we run the RepeatExplorer2/TAREAN (Novák et al. 2017) with default options using as input a total of 12,000,000 reads from the female specimen belonging to the Europe-Local strain reared in the lab. The female was selected because it is the heterogametic sex in Lepidoptera, harboring ZW system in *B. mori*. For the final custom satDNA library we considered the satDNAs directly identified by TAREAN and identified by our manual curation of the RepeatExplorer2 output. We considered as putative satDNAs the clusters with sphere or ring-like shapes within the top clusters. Through Geneious Pro v.4.8.5. (Kearse et al. 2012) the identification of clusters that contained true satDNA sequences were confirmed. Moreover, using this software we also determined the size of satDNA monomers (repeat unit), as well as the consensus sequence for each satDNA family, based in the assembling of all reads of each cluster. The possible similarities between the satDNAs families in the custom library was checked through a BLAST all-to-all analysis with blastn and –e 0.001 options.

The quantification of abundance and divergences of each satDNA family was estimated through RepeatMasker v4.1.4 (Smit et al. 2013-2015) with the “-a” option and the RMBlast search engine. For this, we randomly selected one million of raw reads for each strain of *B. mori* and for *B. mandarina*, aligning them against the sequences of satDNAs identified from *B. mori* genome. It was used dimers for satDNAs families with monomer size ≥100 bp in length or monomer concatenation ranching up 200 bp in length for satDNA families with monomer size shorter than 100 bp in length. Divergence was estimated based in Kimura 2-parameter (K2P) model with the Perl script calcDivergenceFromAlign.pl from the RepeatMasker suite. The satDNAs were named according to the mean abundance between male and female genomes from the individuals belonging to Europe-Local strain.

To have a deeper information about patterns of accumulation and diversification of satDNAs between sex chromosomes and autosomes we run the RepeatMasker using the reads from assembled chromosomes, from a female genome of *B. mori* (ASM3026992v2), and from a male of *B. mandarina* (ASM3026744v2). The landscapes (abundance *versus* divergence) were plotted using a custom script in RStudio (R studio Team, 2024) and the ggplot2 package (Wickham, 2016).

We used the RepeatProfiler tool (Negm et al. 2021) to have better understanding of genomic organization of satDNAs. For this we used separately the reads from male and female genomes from Europe-Local strain (accession numbers **<u>XXX</u>** and **<u>XXX</u>**). The RepeatProfiler tool generate a coverage and base pair composition plot using Illumina sequencing reads through mapping them against each consensus of satDNA sequences. This allows the confirmation of occurrence of in tandem repeated organization by occurrence of hybrid rads between computationally assembled monomers. In this way, RepeatProfiler analysis was run using as reference trimers or multimers up to 200 bp in length, depending on the satDNA family size.

Similarity of the identified satDNA families from *B. mori* genome with previously describes sequences was checked using as query each satDNA against RepBase (Bao et al. 2015) (http://www.girinst.org/). For more accurate data about similarity between the satDNAs and TEs, avoiding only spurious similarities, we considered parameters similar do defined by Tunjić-Cvitanić et al. (2024). In this way, satDNAs with hits with more than 50% monomer coverage and identity > 70% identity were considered as related to TEs.

### 2.4. Mapping of satellite DNAs in the chromosomes of *Bombyx mori* and *Bombyx mandarina*

In other to physical locate the satDNA families on the chromosomes of *B. mori* to know the possible enrichment in specific chromosomes, we applied two strategies, i.e. Fluorescence *in situ* hybridization (FISH) and CHRomosome In Silico MAPPing (CHRISMAPP), following the recommendation of Cabral-de-Mello and Marec (2021) and Rico-Porras et al. (2024), respectively. For FISH we used slides obtained from the larvae of Europe-Local strain and the probes were obtained from oligonucleotides based on the most conserved regions from each satDNA family (**Supplementary Table 1**) through labelling with biotin-16-dUTP (Roche) or digoxigenin-11-dUTP (Roche) using terminal transferase (Roche), following the instructions provided by the supplier. Probe detection was performed using streptavidin conjugated with Alexa Fluor™ 488 (Thermo Fisher Scientific, Waltham, MA, USA) at a concentration of 10 µg/mL for biotin labeled probes and anti-digoxigenin-rhodamine (Roche) at a concentration of 1 µg/mL for digoxigenin labeled probes. Slides were mounted and chromosomes counterstained using VECTASHIELD with 4’-6-diamino-2-phenylindole (DAPI) (Vector Laboratories, Burlingame, CA, USA). Slides were analyzed using a BX51 Olympus® fluorescence microscope (Olympus, Hamburg, Germany) equipped with a CCD camera (Olympus® DP70). Image acquisition and processing were carried out using DP Manager software v1.1.1.71 and Adobe Photoshop CS4 software (Adobe Systems, San Jose, CA, USA).

As the FISH of satDNAs were performed in the chromosomes from larvae maintained in the lab belonging to Europe-Local strain to have a general characterization of its karyotype we also mapped few other probes, including the canonical insect telomeric repeat (TTAGG)*_n_*, 18S rDNA, and genomic DNAs from male and female, i.e. comparative genomic hybridization (CGH). A fragment of the 18S rDNA was amplified using the primers 18S-965 and 18S-1573R (Mullin et al., 2005). The telomeric probe was synthetized by non-template PCR according to Ijdo et al. (1991) using primers (TTAGG)*_5_* and (CCTAA)*_5_*. These two probes were labeled with digoxigenin-11-dUTP (Roche) and FISH was done according to Cabral-de-Mello and Marec (2021). For CGH, the gDNA from male was labeled with biotin-16 dUTP (Roche), while the female gDNA with digoxigenin-11-dUTP (Roche) and the protocol from Traut et al. (1999) with modifications proposed by Dalíková et al. (2017b) was followed.

Several CHRISMAPP were performed using the genomes assembled at chromosome level from a female of *B. mori* (ASM3026992v2) and from a male of *B. mandarina* (ASM3026744v2) using default options suggested by the authors (Rico-Porras et al., 2024). The full monomers of each satDNA were used in the analysis. Considering that it has been documented chromosomal number differences in between *B. mori* (n=28) and *B. mandarina* (n=27) (Banno et al., 2004) we run an analysis of synteny between the two assembled genomes used for CHRISMAPP analysis using the synteny-PlotteR pipeline (Quigley et al. 2023). For this, we retrieved the BUSCO tables from b A3Cat (Feron and Waterhouse, 2022) using the Lepidoptera database from BUSCO and generated linear plot graphics based in chromosomal position of each conserved gene. The chromosomal position of each single copy gene from each species were merged by gene name taking only the “complete” genes in each species. The Z chromosome was identified based on the presence of *Kettin* gene that is known to be in this chromosome (Koike et al., 2003).

## Results

### The chromosomes of *Bombyx mori* Europe-Local strain

As the FISH mapping of the satDNAs was done in the chromosomes obtained from larvae maintained in the lab from local Europe-Local strain (see methods) we performed the classical characterization of its chromosomes. The cytological analysis revealed the chromosome counting of 2n=56 (**Figure 1a,b**), occurrence of the W chromosome in females as evidenced by the sex chromatin, revealing the karyotypes 2n=56, ZZ in male and 2n=56, WZ in female (**Figure 1c**), being consistent with previous descriptions (Traut, 1976; Sahara et al., 2016). Telomeres occupied exclusively the terminal regions of chromosomes (**Figure 1d**) and the 18S rDNA was placed at interstitial position of one chromosome (**Figure 1e**), as described by Traut (1976). As noticed by Sahara et al (2003), through CGH we observed similar labeling of the W chromosome for male and female gDNAs, but it was not more intensely labeled by DAPI. The autosomes and Z chromosome also labeled equally for both gDNA probes, but the signals were a bit less intense than on the W chromosome (**Figure 1f,g**). This evidences that the W chromosome does not differ in DNA concentration or repeat composition in comparison to other chromosomes, but this accumulates more repetitive DNAs, giving more intense gDNA probes signals (Sahara et al., 2003).

**Figure 1.**
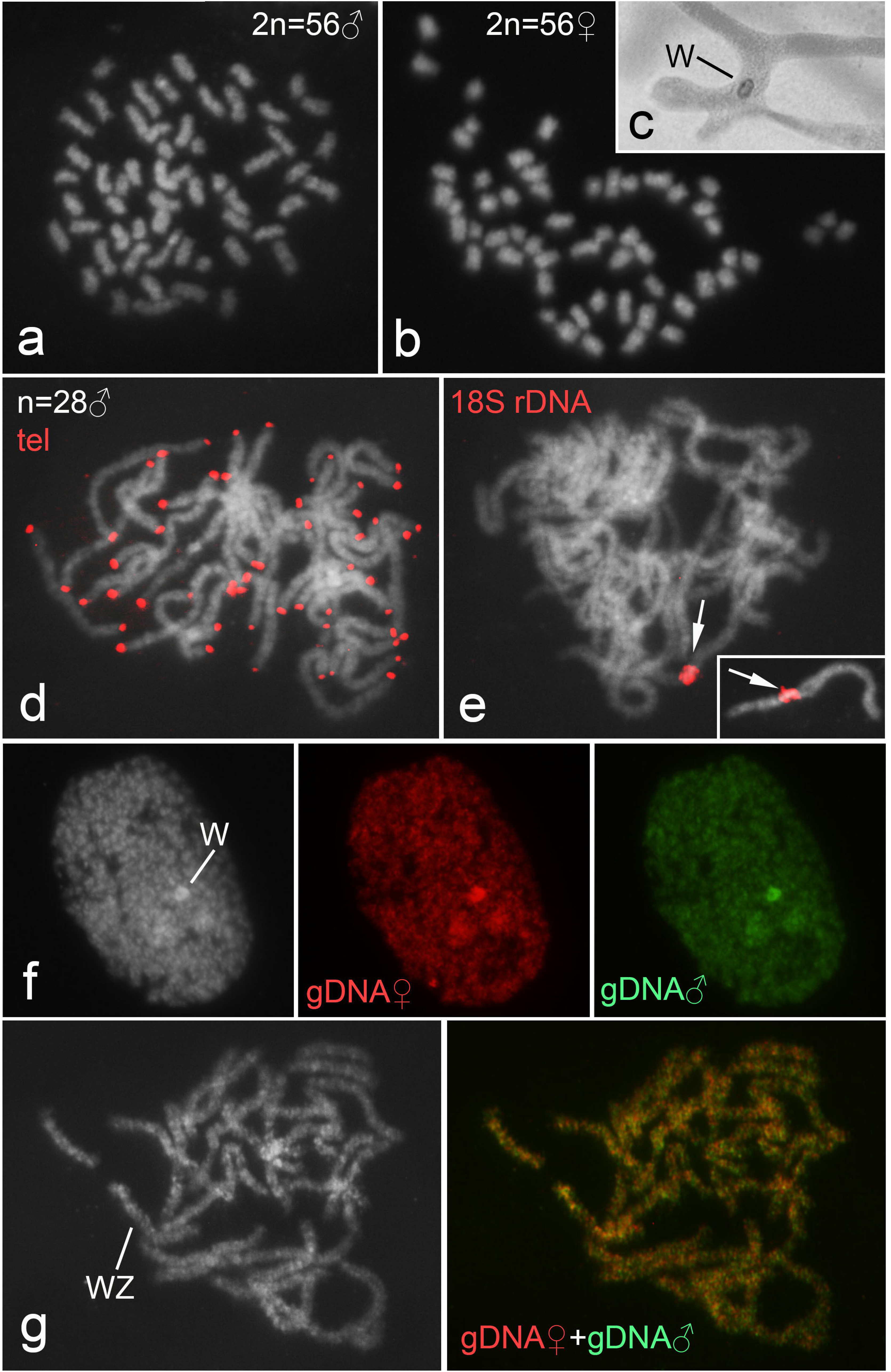
The chromosomal characteristics of *Bombyx mori* belonging to Europe-Local strain. (a,b) Male and female mitotic metaphases obtained from wing disks showing 2n=56 chromosomes in both sexes. The occurrence of sex chromatin in (c) observed in polyploid cells of Malpighian tubes from female insects confirms the occurrence of regular W sex chromosome in the strain. (d,e) Mapping through FISH telomeric motif (TTAGG)*_n_* and the 18S rDNA in male pachytene cells, showing occurrence of telomeres restrict to terminal regions of the chromosomes (d) and an interstitial cluster of rDNA (e). In (e) the inset shows the chromosome harboring the 18S rDNA cluster in detail. (f,g) CGH in polyploid interphase cell (f) and in a female pachytene (g), showing similar labeling for both gDNAs, female (red) and male (green). In both cases it is possible to see that the W chromosome is highlighted. Bar = 10 µm.

### Similarity of satDNA profiles between *Bombyx mori* strains and its wild ancestor *Bombyx mandarina*

Synteny chromosome analysis allowed the identification of the rearranged chromosomes between the two *Bombyx* species with one chromosome (the M chromosome) of *B. mandarina* (chromosome AP028284.1) corresponding to two small chromosomes of *B. mori* (chromosomes AP028184.1, chromosome 14 and AP028197.1 chromosome 27), in accordance with cytogenetic analysis (Banno et al., 2004) and the comparative genome assembly (Lee et al., 2025). The Z chromosome corresponded to the chromosome AP028171.1 in *B. mori* and chromosome AP028283.1 in *B. mandarina*, named as chromosome 1 in both genome assemblies.

Through the analysis of RepeatExplorer2 output it was possible to identify a total of 29 satDNA families in the genome of *B. mori*. The monomer sizes ranged from 13 bp (BmorSat13-13) to 1132 (BmorSat02-1132), with mean size of 300 bp. The A+T content reached a maximum of 78.23% (BmorSat05-124), a minimum of 30.00% (BmorSat17-20), and had a mean of 56.98%. The total abundance for complete satellitome was quite similar between the five strains, ranging from 0.738% to 0.77% with mean of 0.756%. The lowest and highest abundant satDNA families was the same in all strains considering the male and female mean, BmorSat01-575 and BmorSat29-84, respectively. Following the general satellitome abundance, the abundance of each satDNA family was also quite stable among the strains (**Table 1; Supplementary Table 2**). The K2P divergence was also highly similar for individual satDNAs between the strains, with the lowest and highest mean abundance between males and females for BmorSat03-37 (K2P around 1%) and BmorSat21-84 (K2P around 26%), respectively. Bearing in mind the similar abundances and divergence of satDNAs between the strains, as expected, the landscapes (abundance *versus* divergence) were also quite similar. Noticeably, only the BmorSat02-1132 and BmorSat03-379 presented more evident patterns of amplification/homogenization with the main peak of abundance in K2P=0%. Although the satDNA BmorSat01-575 is the most abundant, and putatively experienced waves of amplifications, it presented more than one main peak of values of K2P. Other satDNAs also experienced a more characteristic recent wave of amplification/homogenization, like BmorSat10-80 and BmorSat16-38. The other satDNAs, in general, experienced older amplification and accumulated nucleotide differences (**Figure 2, Supplementary Figure 1**).

**Figure 2.**
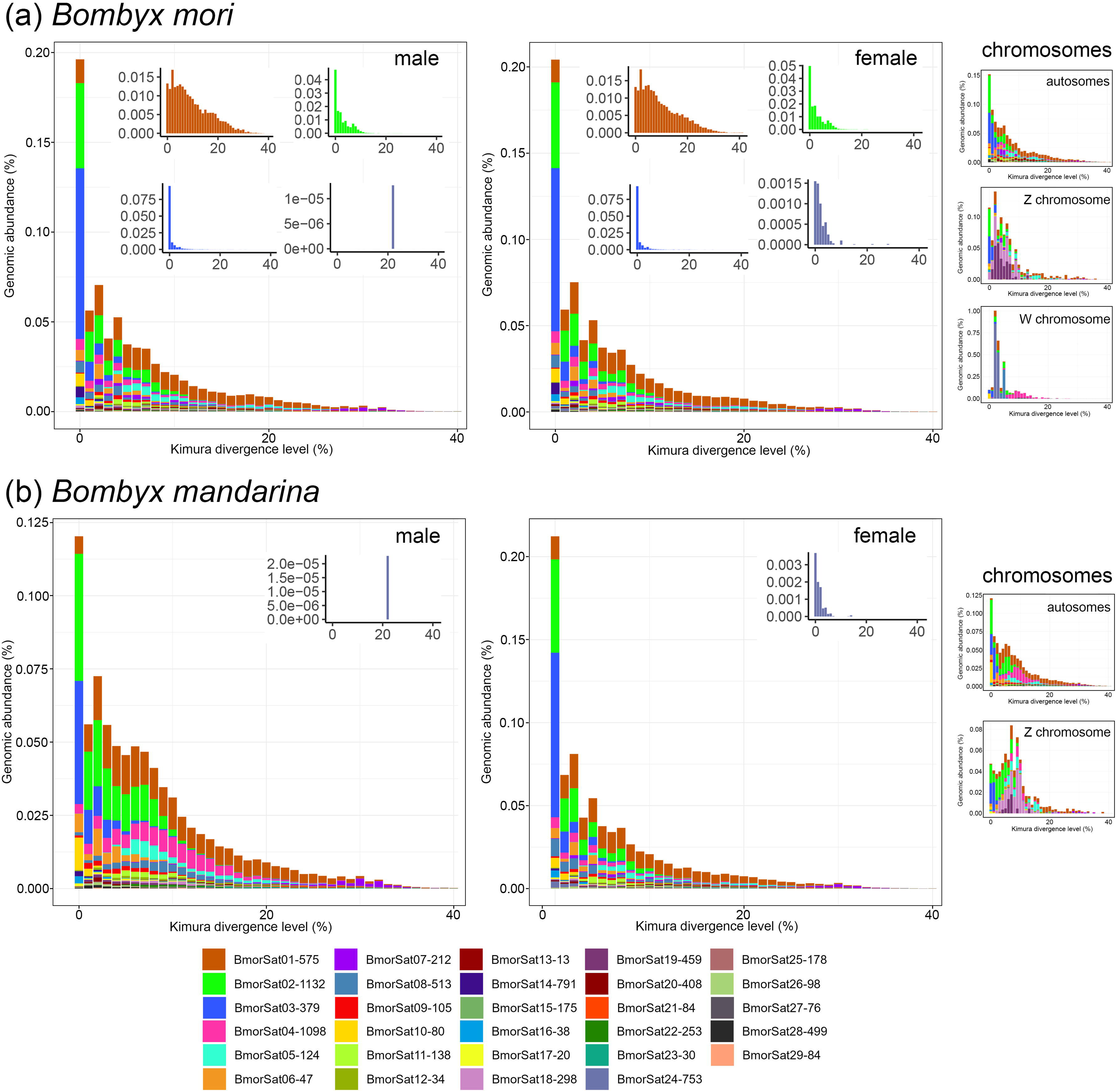
Landscapes (abundance *versus* K2P divergence) for satellite DNAs in the genomes of Europe-Local strain of *Bombyx mori* (a) and *Bombyx mandariana* (b). Inside the main box of each landscape is showed individual landscapes for specific satDNA families. In the right side it is showed the landscape for distinct chromosomes, including all autosomes, the W chromosome, and the Z chromosome. Each color represents on satDNA family. Note the difference in abundance and K2P divergence between male and female for the satDNA Bmori24-753 (light purple). Moreover it is evident differences in the landscapes for the specific chromosome.

**Table 1.**
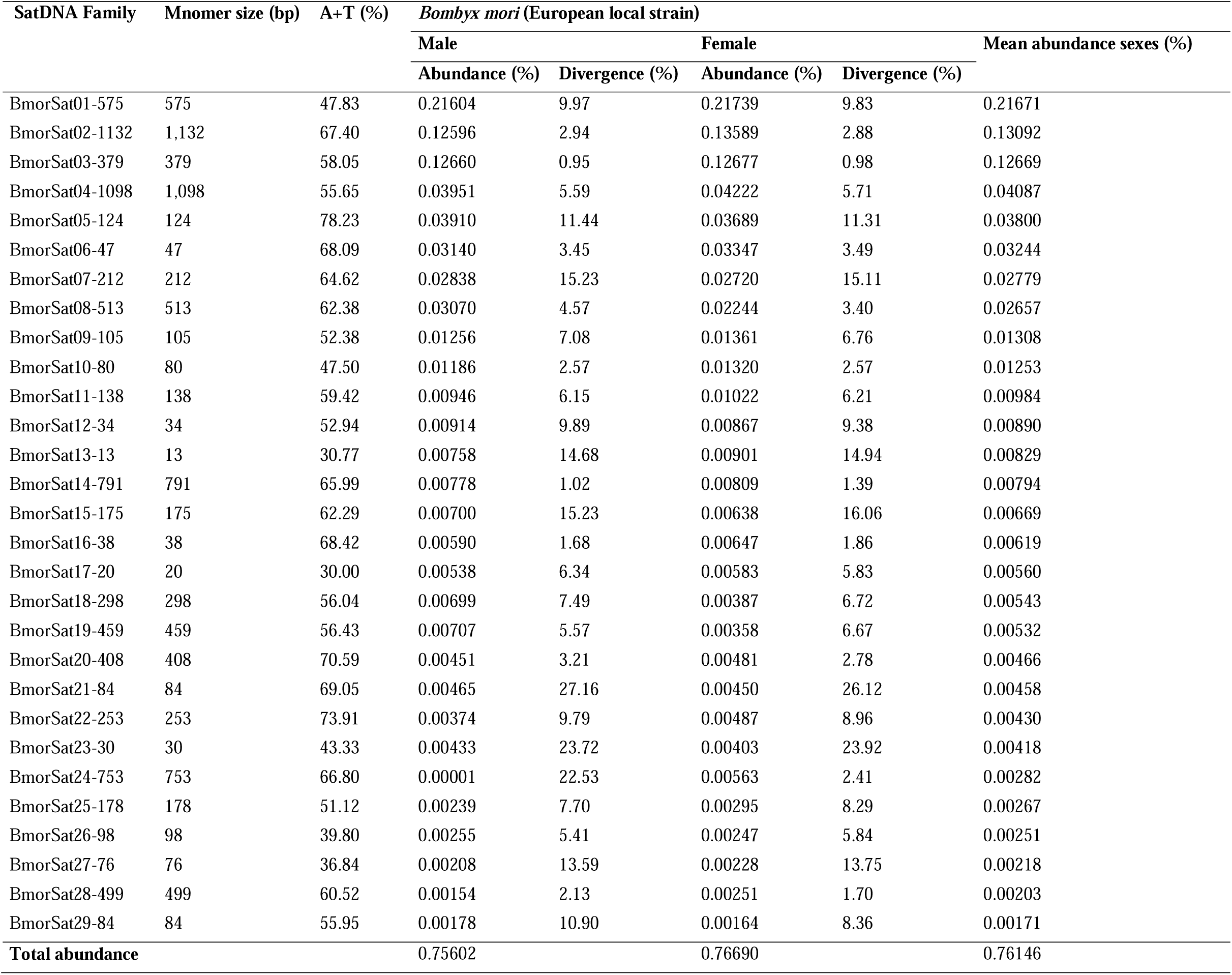
Satellite DNAs identified in the genome of *Bombyx mori* from Europe-Local strain and their characteristics.

In contrast with the high similarity of abundances and divergences of the satDNAs between the strains, we noticed a clear differentiation for the BmorSat24-753 between male and female genomes across all strains. Invariably, this satDNA occurred in consistently very low abundance in males compared to females. Among males it presented a mean abundance of 0.00001% in females the mean abundance was 0.0069%. In females the mean divergence across strains for this satDNA was much lower than in males, being 2.09% in females and 22.4% in the three males were this satDNA was present, i.e. quantified by RepeatMasker. The landscape of females clearly shows a pattern of recent amplification in comparison to males (**Figure 2**).

The analysis of the whole satellitome of *B. mori* in *B. mandarina* genome revealed similar picture to the observed across *B. mori* strains. The mean abundance (male and female) of the satDNA in this species was 0.8%, with variation from 0.00187% (BmorSat29-84) to 0.225% (BmorSat01-575). All satDNAs had similar abundances and divergence between sexes, except the BmorSat24-753 that similarly to *B. mori* was more abundant and less divergent in females than in males. In males the abundance of BmorSat24-753 was 0.000023% and the divergence 22.53%, while in females the abundance was 0.0088% and the divergence 1.76% (**Figure 2; Supplementary Table 3**).

In order to test if there is differential patterns of amplification and diversification between the autosomes and sex chromosomes, we analyzed the abundance and divergence of each satDNAs separately between chromosomes, i.e. W, Z, and autosomes. In *B. mori* the landscape for autosomes was quite similar to general landscape for the whole genome. On the contrary, the sex chromosomes presented differential patterns of accumulation and diversification of satDNAs compared to autosomes. On Z chromosome the satDNAs BmorSat18-298 and BmorSat19-459 were differentially amplified. While, for the W chromosome a clear pattern of accumulation was noticed for the satDNA BmorSat24-753. Similarly in *B. mandaria* the satDNAs BmorSat18-298 and BmorSat19-459 were also differentially amplified on the Z chromosome. For *B. mandarina* genome assembly the W is not included (**Figure 2; Supplementary Table 4**).

The RepeatProfiler analysis confirmed occurrence of tandem organization for all satDNA families with occurrence of hybrid rads in the monomer connections. The coverage was more similar along the entire sequences for most satDNAs, while in few cases, as in BmorSat01-575, BmorSat04-1098, and BmorSat16-30 variable coverage along the sequence was observed, suggesting occurrence of truncated copies of these satDNAs and also putative spread organization, i.e. not in tandem. This pattern was similar for male and female genomes (**Figure 3a,b; Supplementary Figure 2**), except for the satDNA BmorSat24-753 that is enriched in the female genome. In this case, it is clearly observed that this repetitive DNA is in tandem in the genome of female, with similar coverage along the sequence and in its monomer connections. On the contrary, in male no coverage was observed, putatively due to the low amount of this repeat and/or its non-tandem organization (**Figure 3c**).

**Figure 3.**
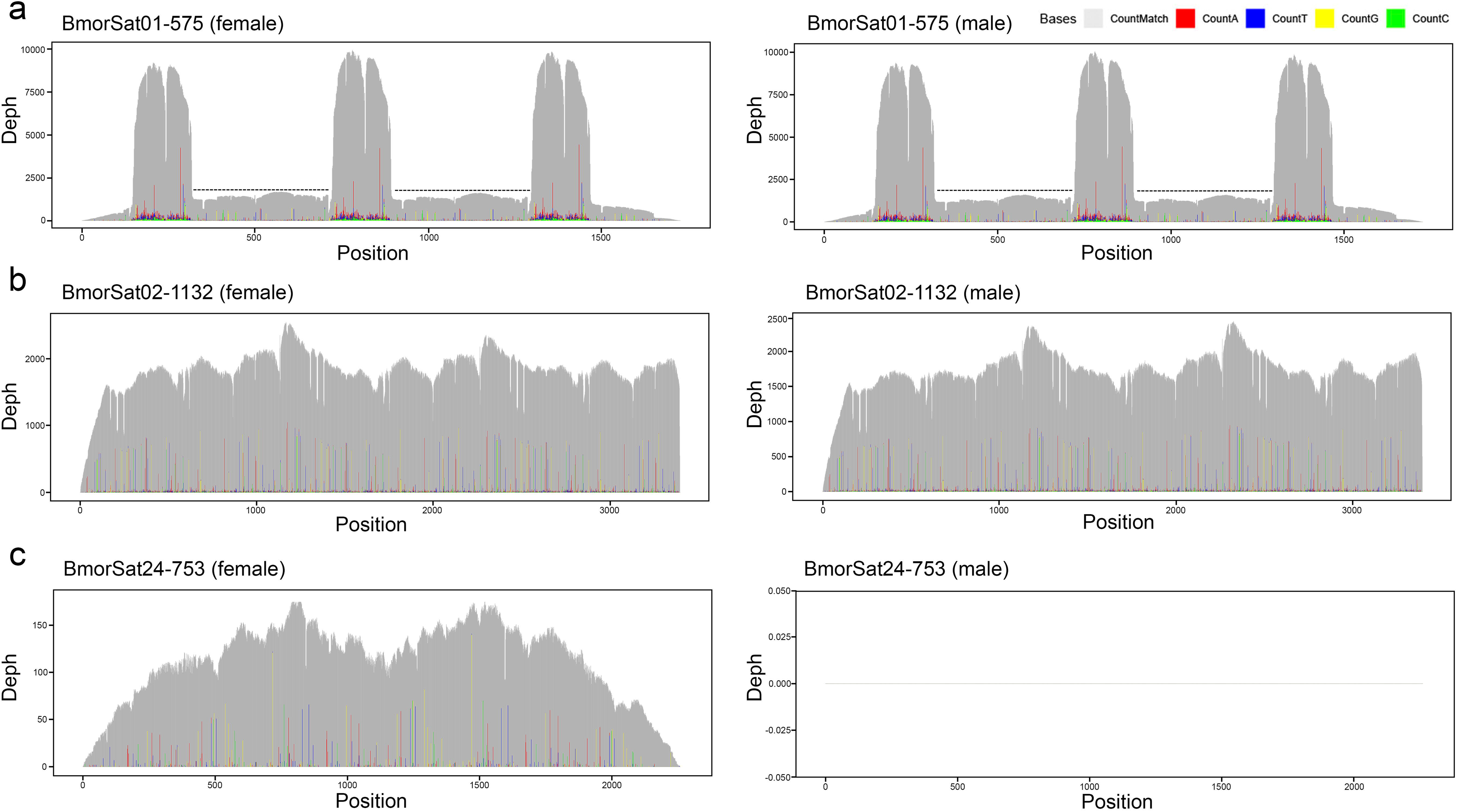
Examples of RepeatProfiler analysis of satDNAs in *Bombyx mori*, showing differences of coverage along the monomers and in their connections. In (b) it is evidenced different coverage along the monomers and in their connections in for BmorSat01-575, evidencing the occurrence of this sequence in tandem organization, but also as spread repetitions. Note the decrease of coverage indicated by dashed lines. (b) Similar read coverage along the monomer and in their connections, that corresponds to hybrid reads, revealing the tandem organization for BmorSat02-1132. In (c) it is showed the absence of coverage for BmorSat24-753 in males. The other RepeatProfiler could be seen in **Supplementary Figure 2**. The data was obtained from short genomic reads form Europe-Local strain.

### Relationship between satDNAs and TEs in *Bombyx mori* genome

The CENSOR analysis using RepBase database revealed the occurrence of six satDNAs related to distinct TEs, as follows: BmorSat01-575 has full similarity with LTR-2_BM-I element and LTR-2_BM-LTR; BmorSat02-1132 has full similarity with TE-1_BM element; BmorSat05-124 has extensive similarity with Helitron-1_RPr; and BmarSat29-84 has almost full similarity with DIRS-8B_DR element. The BmorSat03-379 and BmorSat10-80 have similarity with TE-2_BM, being full similarity for BmorSat10-80 and extensive similarity for BmorSat10-80. The elements LTR-2_BM-I, LTR-2_BM-LTR, TE-1_BM, and TE-2_BM were previously described in the genome of *B. mori*, while Helitron-1_RPr in the genome of the true bug *Rhodnius prolixus* (**Table 2, Supplementary Figure 3**). As the element DIRS-8B_DR was described in the fish *Danio rerio* we did not consider for further analysis.

**Table 2.**
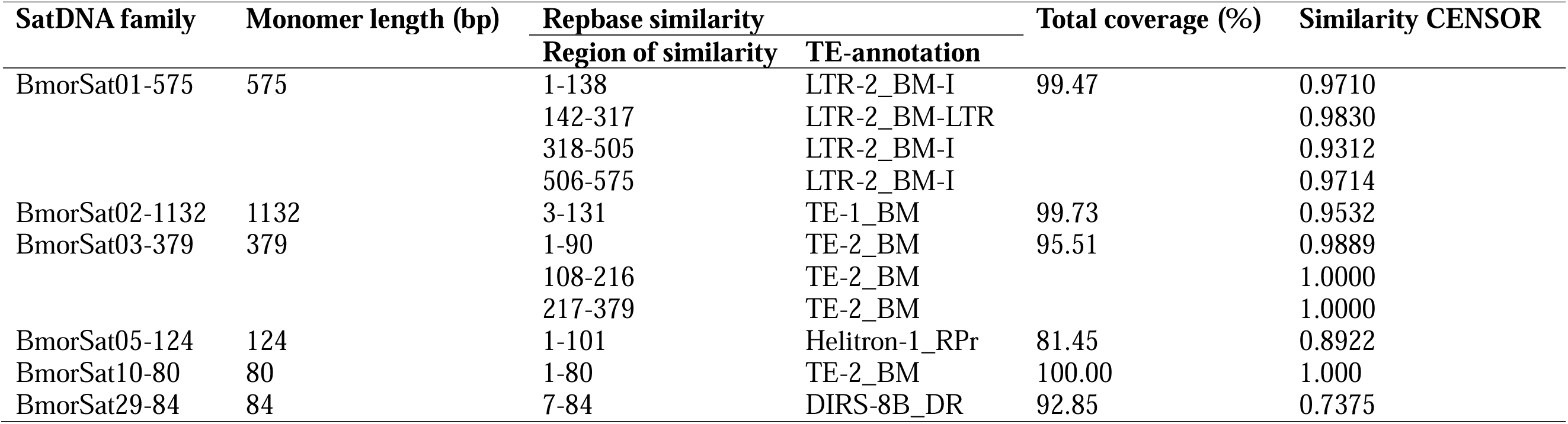
Summary data for satDNAs related to TEs found in the genome of *Bombyx mori*.

The comparison of full sequence of TEs with the satDNAs related to TEs revealed the origin of these satDNAs from distinct regions of the TEs and in variable ways (**Figure 4**). The BmorSat05-124 (**Figure 4a**) and BmorSat10-80 (**Figure 4b**) were originated from a small central part of the TE elements, Helitron-1_RPr and TE-2_BM, respectively. The BmorSat02-1132 (**Figure 4c**) was originated from almost full length of the TE-1_BM, only with a large final deletion. Similarly, the BmorSat03-379 was originated from a large portion from the TE-2_BM, being a shorter sequence because of some internal and final deletions (**Figure 4c**). The BmorSat01-575 has the most intriguing origin, involving two different TEs, elements LTR-2_BM-I and LTR-2_BM-LTR. A terminal part of BmorSat01-575 was originated from a terminal part of LTR-2_BM-I joined with the entire LTR-2_BM-LTR. The other terminal part of the LTR-2_BM-I was added in internal position and one internal position from LTR-2_BM-I formed the other end of the satDNA (**Figure 4d**).

**Figure 4.**
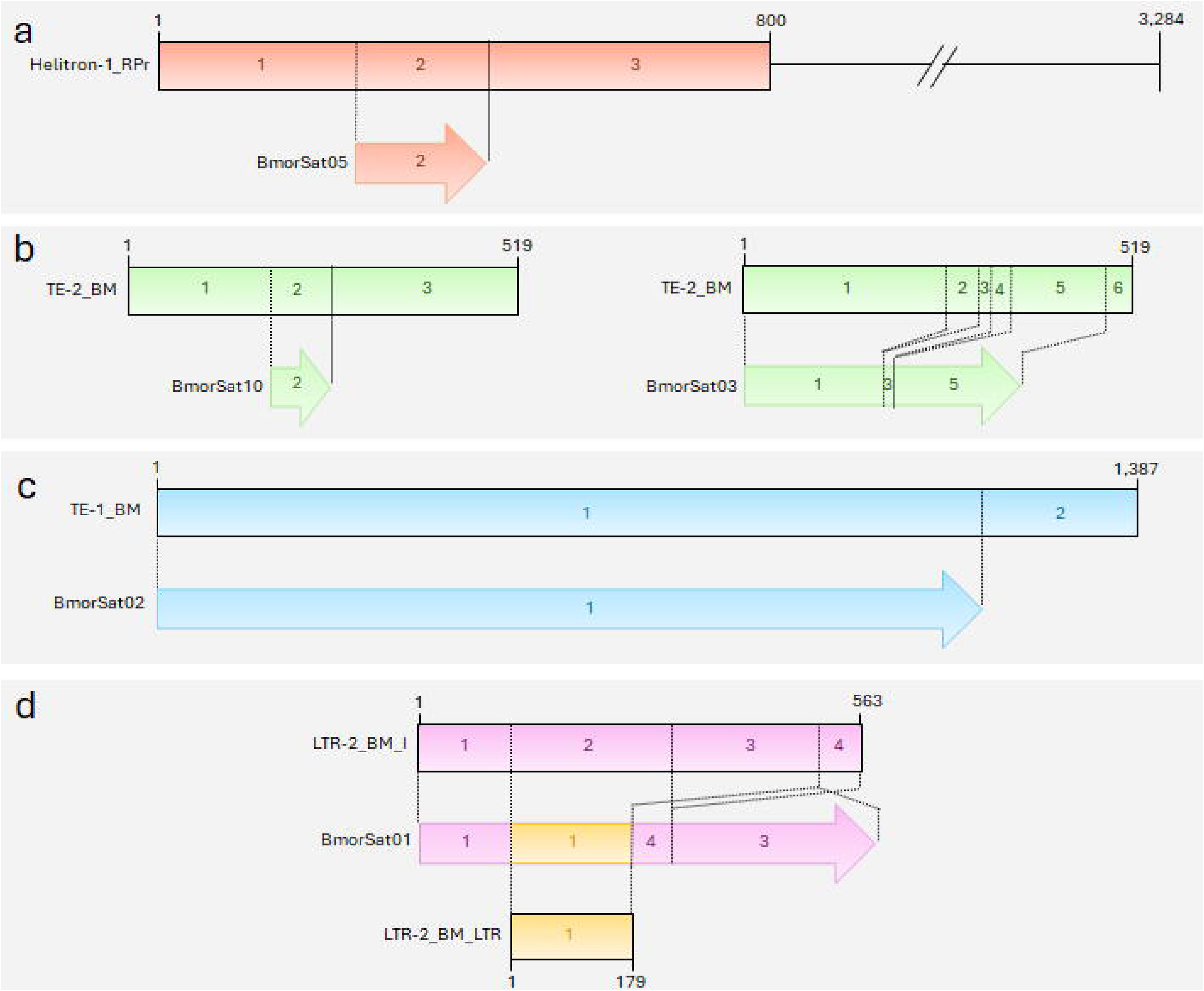
Patterns of origin of four satDNA families from transposable elements (TEs) in *Bombyx mori* genome, (a) BmorSat05-124, (b) BmorSat10-80 and BmorSat03-379, (c) BmorSat02-1132, and (d) BmorSat01-575. More details about TEs masked regions identified by CENSOR analysis that are involved with satDNAs origin are given in **Supplementary Figure 3**.

### The spread organization of satDNAs in *Bombyx mori* and *Bombyx mandarina* chromosomes

Due to the low abundance and spread organization of satDNAs we were able to see signals through FISH only for the two most abundant satDNA families, BmorSat01-575 and BmorSat02-1132 (**Figure 5a**), revealing spread signals. This data is consistent with observed through CHISMAPP for these two elements (**Supplementary Figure 4**). As for the other satDNAs we did not notice FISH signals, due to their low abundance and spread organization, that limits the technique resolution, their locations were defined through CHRISMAPP (**Figure 5b**). As for the two most abundant satDNAs, most other elements were mostly dispersed in all chromosomes. Controversially, some satDNAs presented enrichment in few or specific chromosomes, including or the BmorSat09-105, BmorSat11-138, BmorSat14-791, BmorSat18-298, BmorSat19-459, BmorSat20-408, BmorSat21-84, BmorSat22-253, BmorSa23-30, BmorSat24-753, BmorSat26-98, BmorSat28-499, BmorSat29-84. For BmorSat24-753 the enrichment was observed exclusively on the W chromosome, consistent with the abundance differences between male and female genomes (**Figure 5b, Supplementary Figure 3**).

**Figure 5.**
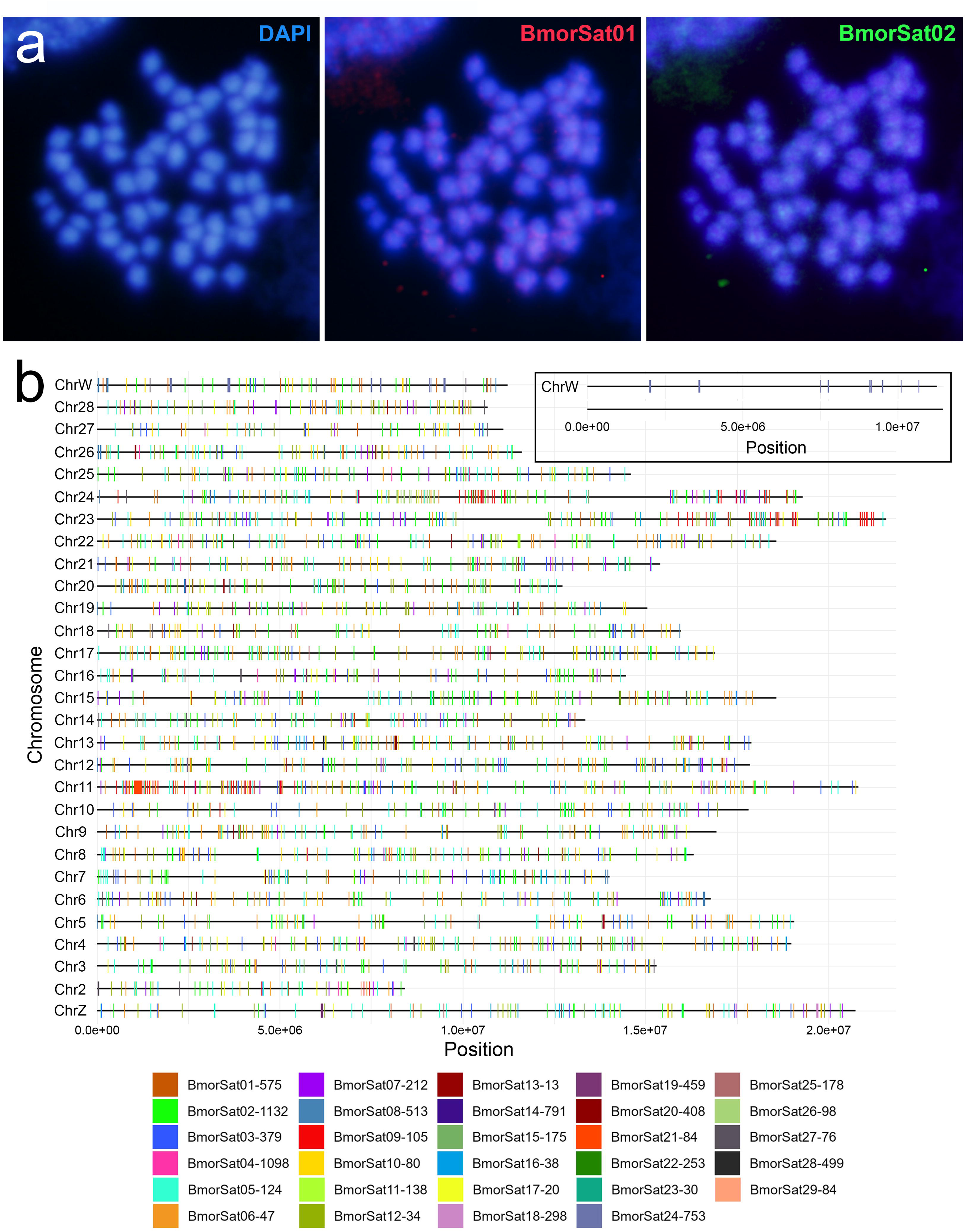
Chromosomal organization of the satDNAs in *Bombyx mori*. (a) FISH mapping of BmorSat01-575 (red) and BmorSat02-1132 (green) in mitotic metaphase obtained from female wing disk. (b) CHRISMAPP analysis of all 29 satDNA families. Note the remarkable spread distribution for most satDNAs that are represented by distinct colors. The box in (b) shows the exclusive location of BmorSat24-753 in the W chromosome. For individual CHRISMAPP images for each satDNA could be seen in **Supplementary Figure 4**.

The distribution of satDNAs in the chromosomes of *B. mandarina* analyzed through CHRISMAPP was quite similar to observed in *B. mori*, with slightly differences only in number of clusters mapped, due to differences in abundance for the satDNAs between the species. Interestingly the satDNAs BmorSat18-298 and BmorSat19-459 occupied specifically similar position on the Z chromosome in both species (**Supplementary Figure 4**).

## Discussion

Our detailed analysis of the *B. mori* satellitome contributes to a better understanding of genome organization and the evolutionary patterns of the satDNA fraction in this model species. Compared to TEs, the role of satDNAs in genome evolution have been poorly investigated, not only in *B. mori* but also in Lepidoptera as a whole (Ray et al., 2019; Baril and Hayward, 2022; Höök et al., 2023; Perrier et al., 2024; Wright et al., 2024). But the satDNAs sequences could also act as important genomic players in Lepidoptera, promoting chromosome diversification, as suggested for *Leptidea* species (Cabral-de-Mello et al., 2024) and in the repositioning of rDNA (Dalíková et al., 2023), being necessary better exploration of this genomic fraction. In *B. mori* the genome size is about 458 Mb (Tong et al., 2022) and low heterochromatin content is observed (Yoshido et al., 2005), consistently with the low amount of satDNAs. This suggests that, compared to other repeat types such as TEs that are highly abundant, corresponding to about 475 Mb in the Nichi01 genome (Waizumi et al., 2023), satDNAs have a lower impact on the genome organization of this species, as they are low abundant. Similarly, in Lepidoptera as a whole, despite the early stage of satellitome research, most studied species with low heterochromatin content exhibit only small amounts of satDNA in their genomes (Cabral-de-Mello et al., 2021; Gasparotto et al., 2022). This contrasts with a few species, such as those in the *Leptidea* genus, which possess large amounts of satDNA and extensive heterochromatin blocks (Cabral-de-Mello et al., 2024). Other characteristics of the *B. mori* satellitome align with those observed in other lepidopterans, including: (i) A+T enrichment, which is consistent with the general composition of its genome and for Lepidoptera (Wright et al., 2024; Lee et al., 2025); (ii) the absence of a principal, i.e. highly abundant and predominant satDNA family, a common pattern in lepidopterans with low heterochromatin content (Cabral-de-Mello et al., 2021; Gasparotto et al., 2022; Haq et al., 2022); and (iii) high variability in monomer size, including satDNAs with large sequence units, also observed in other insects (Anjos et al., 2023; Rico-Porras et al., 2024; Vidal et al., 2025).

Although the satDNA are fast evolving sequences and it is experienced extensive differentiation among species, even in related ones, it is expected that in early diverging lineages the changes are less intense, as observed here in the five *B. mori* strains, evidencing satDNA stability during the domestication process, dating about 5000 years ago. This stability could be exemplified by the most abundant satDNA, BmorSat01-575, and the least abundant, BmorSat29-84, that were consistent across all *B. mori* strains. Additionally, the divergence landscapes were highly similar among strains, with most satDNA families exhibiting more ancient amplification patterns (higher divergence), except for BmorSat02-1132 and BmorSat03-379, which showed evidence of more recent homogenization (less divergence). In the noctuid moth *Spodoptera frugiperda* the analysis of satDNAs in distinct populations revealed differences in copy number for seven satDNA families (Haq et al., 2022). As expected, in comparison to *B. mandarina*, higher quantitative differences are evidenced, but all satDNAs are shared between the species revealing conservation of satDNAs after speciation. Despite evolutionary divergence, both species maintained a satellitome abundance of ∼0.75–0.8% and the most abundant satDNA was the same (BmorSat01-575) between them, even in similar amounts, i.e. mean between sexes of 0.216% (*B. mori*) and 0.225% (*B. mandarina*), suggesting that this element could be important in the genome organization of the species, being conserved. Additionally, divergence landscapes were highly similar, with most satDNA families exhibiting divergence common for old divergent elements, except for BmorSat02-1132 and BmorSat03-379, which showed evidence of recent homogenization. Besides stability for satDNAs in both species general chromosome structure is also conserved, being the species differentiated by only one polymorphic chromosome fusion in *B. mandaria* (Banno et al., 2004; Lee et al., 2025). For *B. mori* strains the occurrence of the same diploid number, including the strain studied here have been reported (Yoshido et al., 2005; Sahara et al., 2016; Li et al., 2024; Lee et al., 2025).

The low pattern of homogenization observed for most satDNA families could be consequence of the arrangements of these repeats on the chromosomes of the species. Though FISH and mainly by CHRISMAPP analysis it is evident the dispersed arrangement of all satDNAs, forming multiple small clusters along the whole chromosomes, that are less prone to homogenize than large repetitive blocks, frequently formed by heterochromatin, that are absent in the chromosomes of *B. mori*. The hypothesis that most satDNAs exist in low-copy dispersed arrangements could be supported by the difficult to visualize clusters cytologically by FISH. Similar distribution for satDNAs have been observed through FISH in other lepidopterans, that in general have satDNAs dispersed along entire chromosomes (Věchtová et al., 2016; Cabral-de-Mello et al., 2021; Gasparotto et al., 2022). According to Cabral-de-Mello et al. (2021) this pattern of distribution in short arrays might be caused by the fact that special environment is necessary for the formation of long arrays, i.e. heterochromatin, that is in general lacking in Lepidoptera, although there are exceptions, as documented in *Leptidea* (Zrzavá et al., 2018; Cabral-de-Mello et al., 2024).

The only extensive occurrence of heterochromatin in Lepidoptera is observed mainly in the W chromosome, that in general is fully heterochromatic and more probable to accumulate repetitive DNAs (Sahara et al., 2012). This is consistent with the higher abundance and lower K2P divergence of some satDNAs in female genome of *B. mori*, indicating accumulation on the W element through differential amplification, suggesting functional or structural role in W chromosome evolution. Analysis based in assembled chromosomes also revealed higher general abundance of satDNAs in the W chromosome than in other chromosomes. Moreover, this element harbors a smaller number of satDNAs families in comparison to other chromosomes, suggesting elimination of repeats along with amplification of specific ones. The most evident example is the BmorSat24-753 that is tandemly organized exclusively in the female genome, presents higher sequence homogenization in females, supporting the hypothesis of W-specific satDNA turnover, potentially linked to heterochromatin organization or gene regulation. As for most other satDNAs, signals were not observed for BmorSat24-753 through FISH even in the W chromosome, being the evidence of accumulation of this element on W chromosome revealed by CHRISMAPP. This supports the absence of long arrays for the BmorSat24-753. The W chromosome as a repository of repeats in *B. mori* has been also supported by accumulation of LTR and non-LTR retrotransposons (Abe et al., 2005; Han et al., 2024; Lee et al., 2025). Accumulation of repetitive DNAs in W chromosomes of Lepidoptera were observed in *Plodia interpunctella* (the satDNA PiSAT1, Daliková et al., 2017), *Diatraea saccharalis* (some TEs, Gasparotto et al., 2022), and *Peribatodes rhomboidaria* (mainly LINE or LTR retrotransposons, Hejníčková et al., 2023). Moreover, general identification of repeats was performed in a limited number of species, including *Ephestia kuehniella* (Traut et al., 2013), *Cydia pomonella* (Fuková et al., 2007), some Crambidae species (Cabral-de-Mello et al., 2021), and *Leptidea* representatives (Cabral-de-Mello et al., 2024). Accumulation of repeats is a common feature of heterogametic sex chromosomes for many organisms, influenced by decrease of recombination, that act as important process in sex chromosomes differentiation. Although, most evidence gathered until now indicates that TEs are the main repeats with tendency to colonize the W chromosome of Lepidoptera (Abe et al., 2005; Traut et al., 2013; Hejníčková et al., 2023), the satDNAs could be also important players as revealed here for *B. mori*, deserving more analysis on other lepidopterans.

On the contrary to W chromosome, the content of satDNA of Z chromosome of *B. mori* is much more similar to the autosomes. The similarity of Z chromosome with autosomes in Lepidoptera was postulated by cytogenetic analysis and genome assembly, that revealed mainly euchromatic nature and occurrence of multiple genes in both chromosomes (Sahara et al., 2012; Wright et al., 2024). Among the 29 satDNAs, the analysis based in chromosome assemblies, 18 satDNAs are shared between the Z chromosome and autosomes, but only eight are shared between W chromosome and autosomes. Similar situation is noticed in *B. mandarina* with 15 satDNAs shared between Z chromosome and autosomes. Although similarities are noticed for the autosomes and Z chromosomes, this last element has some degree of differentiation with accumulation of two satDNA families (BmorSat18-298 and BmorSat19-459) that are exclusively present in the Z chromosome in both species, occupying similar position. This support that the conservation of Z chromosomes between the two species besides the gene content occurs also at certain levels for highly dynamic elements, as the satDNAs.

Finally, it is noticeable in the genome of *B. mori* the occurrence of some satDNAs families with TE similarities, suggesting that TEs are involved in satDNA origin. In other species, the refinement of analysis based in bioinformatic tools the body of evidence concerning relationship between satDNAs and TEs has been growing (Zattera and Bruschi, 2022; Tunjić-Cvitanić et al., 2024). For *B. mori* the origin of satDNAs occurred putatively by multiply processes as the simple tandemization of specific parts of TEs, like noticed for BmorSat02-1132, BmorSat05-124, and BmorSat10-80; Tandemization of distinct parts of the same TE, as observed for BmorSat03-379; or by fusion of distinct parts of different TEs, as revealed for BmorSat01-575. The genome of *B. mori* clearly exemplifies how complex and dynamic could be the origin of satDNAs from TEs. Recently extensive dominance of transposable element related satDNAs was documented in oysters (Tunjić Cvitanić et al., 2024) and in *Drosophila*, the role of TEs as building blocks of satDNAs has been documented (Dias et al., 2014; Silva et al., 2023). In *B. mori*, most part of satellitome abundance is from TE derived elements, representing about 69% of satellitome content (0.52485% of 0.76146), evidencing that the TEs could be a main contributor in the satellitome origin and expansion.

Our findings underscore the evolutionary stability of satDNA composition, abundance, and chromosomal organization in *B. mori* strains and its ancestor species *B. mandarina*, while highlighting sex chromosome-specific satDNA dynamics. The differential accumulation of BmorSat24-753 on the W chromosome, coupled with its conserved female-biased enrichment, suggests a potential role in W chromosome evolution. The widespread association of satDNAs with TEs further supports a transposon-derived origin for many repeat families, reinforcing the interplay between repetitive DNA classes in shaping genome architecture. Moreover, the occurrence of satDNAs on euchromatin can have functional implications, as they can act as regulators of gene transcription. Future studies integrating chromatin accessibility and transcriptional profiling will be essential to elucidate the functional implications of these repetitive elements in *Bombyx* genome evolution.

## Acknowledgements

This work was funded by Fundação de Amparo à Pesquisa do Estado de São Paulo (FAPESP, process numbers 2023/02581-02), the Coordenação de Aperfeiçoamento de Pessoal de Nível Superior—Brasil (CAPES, code 001), Conselho Nacional de Desenvolvimento Científiico e Tecnológico (CNPq, process number 308290/2020-8), and the Spanish Junta de Andalucía through the “Programa Operativo FEDER Andalucía 2014–2020”. Pablo Mora was supported by FAPESP 2024/01521-9.

## Supplementary material

**Supplementary Figure 1.** Satellite DNAs landscapes (abundance *versus* K2P divergence) from males and females of four *Bombyx mori* strains.

**Supplementary Figure 2.** RepeatProfiler graphs for all satDNAs identified in the genome of *Bombyx mori*. The analysis was done using short reads from Europe-Local strain.

**Supplementary Figure 3.** CENSOR analysis for the five satDNAs related to transposable elements (TEs) identified in *Bombyx mori* genome. In red it is showed the masked regions.

**Supplementary Figure 4.** Individual CHRISMAPP images for the 29 satDNAs identified in the *Bombyx mori* genome.

**Supplementary Figure 5.** General and individual CHRISMAPP images for the 29 satDNAs of *Bombyx mori* in the chromosomes of *Bombyx mandarina*.

**Supplementary Table 1.** Oligonucleotides of the 29 satDNAs used as probes for chromosomal mapping through FISH in the chromosomes of *Bombyx mori*.

**Supplementary Table 2.** Abundance and K2P divergence for the 29 satDNAs in the five strains of *Bombyx mori*, including male and female individuals.

**Supplementary Table 3.** Abundance and K2P divergence for the satDNAs prospected from the genome of *Bombyx mori* that are present in *Bombyx mandarina* genome, including male and female individuals.

**Supplementary Table 4.** Abundance and K2P divergence for the 29 satDNAs in the assembled chromosomes of *Bombyx mori* and *Bombyx mandarina*.

## Notes

### Competing Interest Statement

The authors have declared no competing interest.

