## Supplementary figures and images for "Advancing the understanding of the *Bombyx mori* genome through satellite DNA analysis: low strain differentiation and relationship with transposable elements"

### Supp fig 1

**Supplementary Figure 1.**

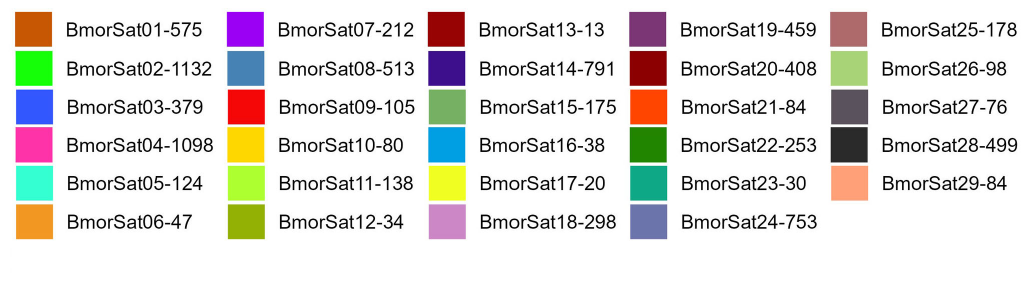

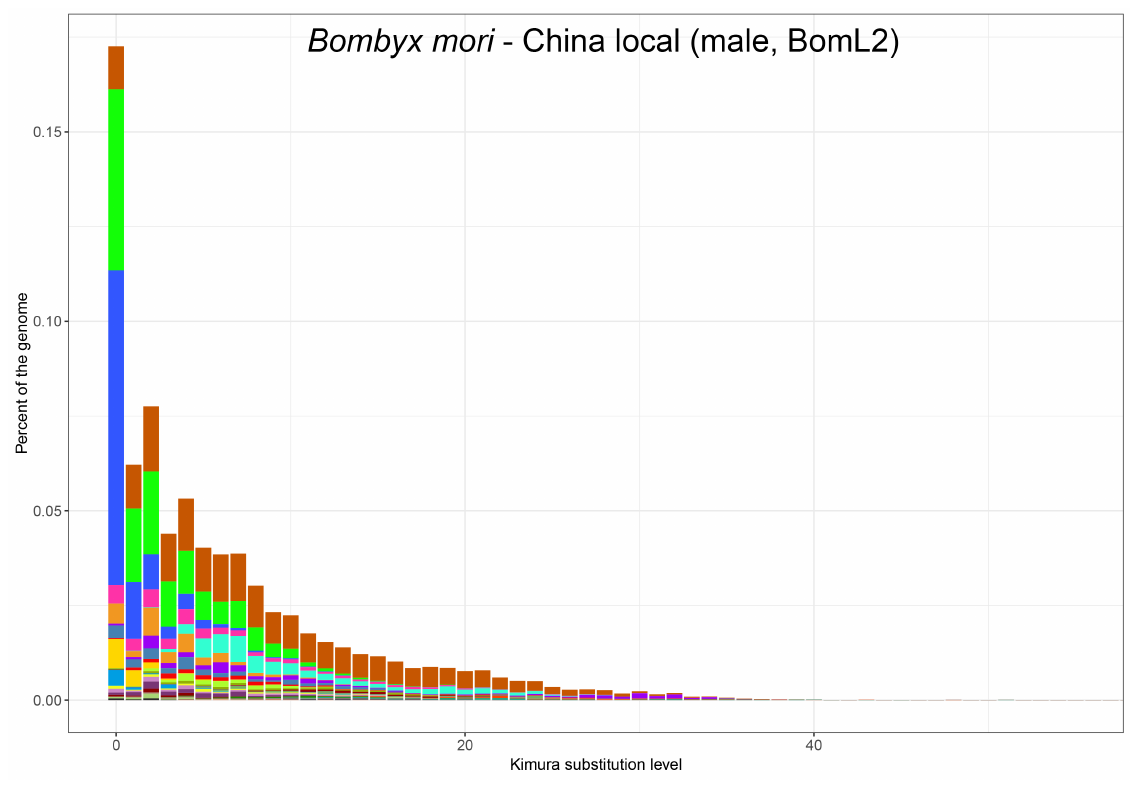

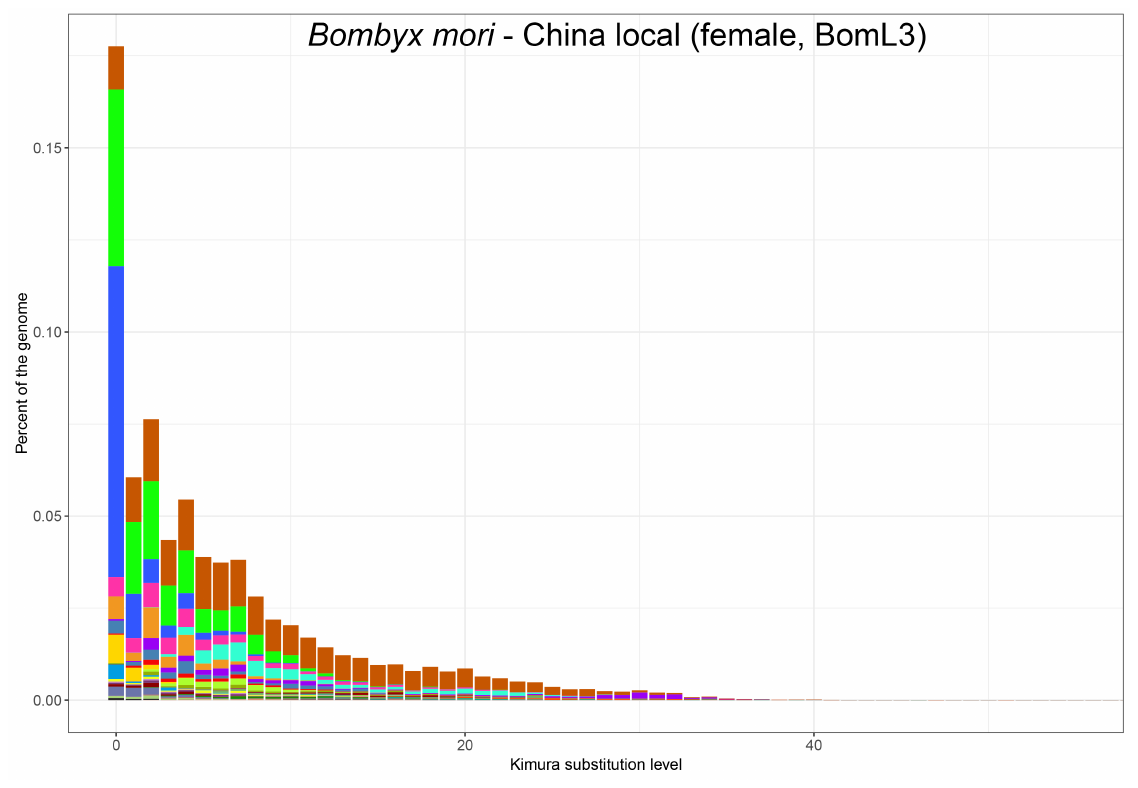

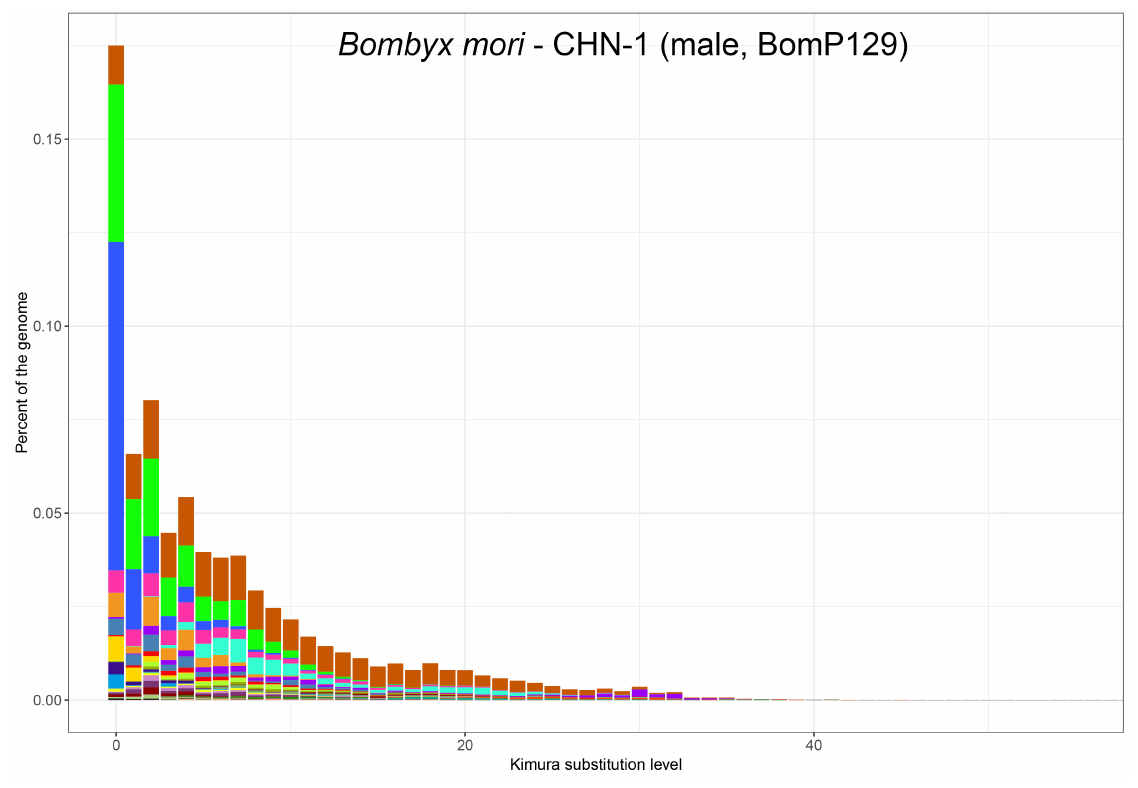

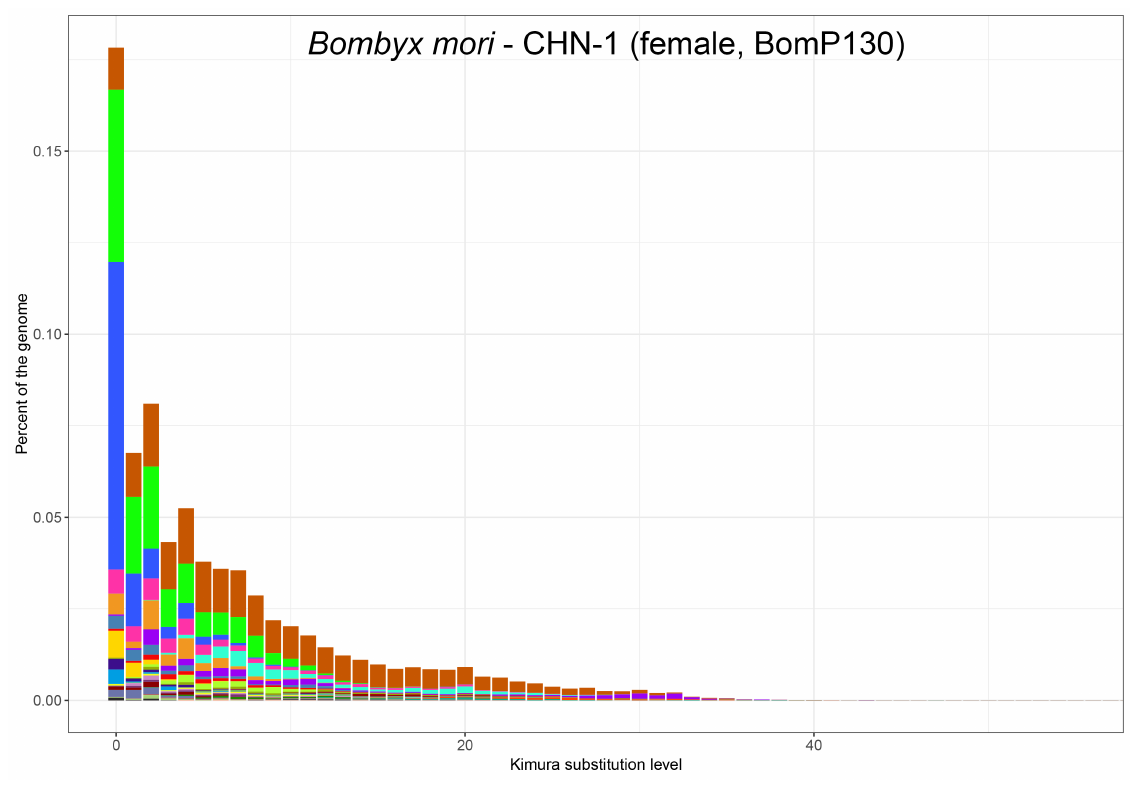

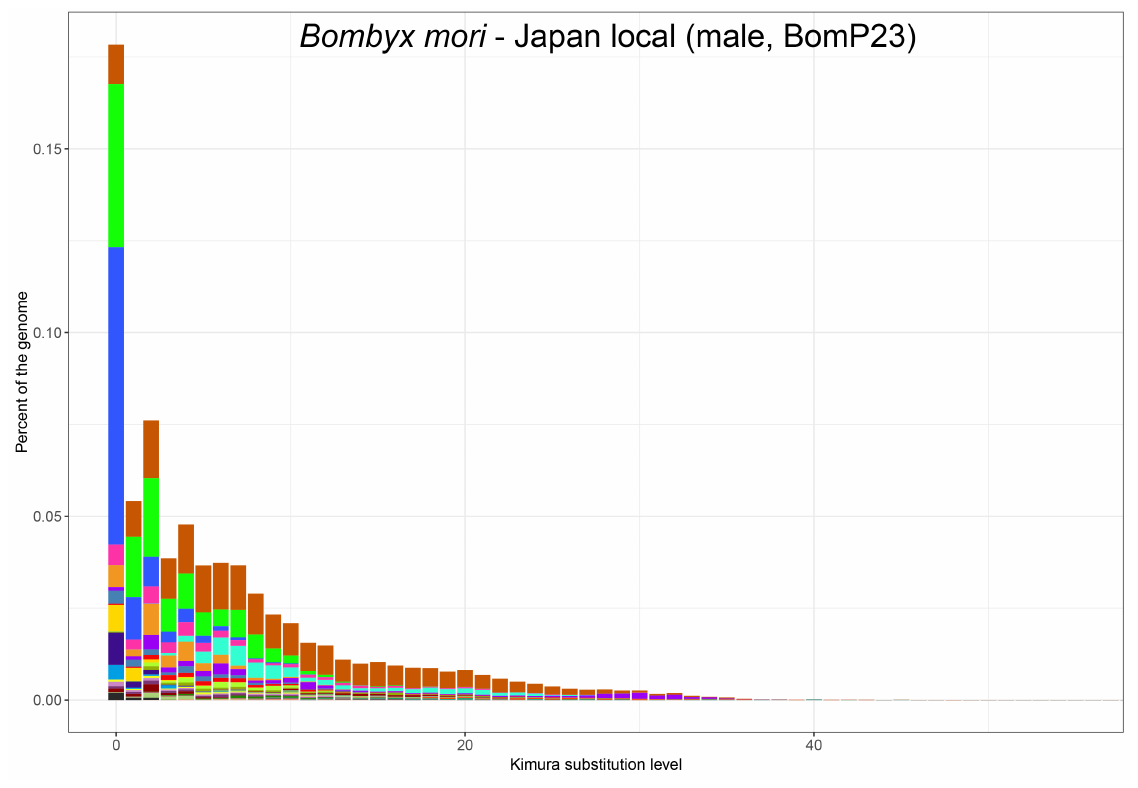

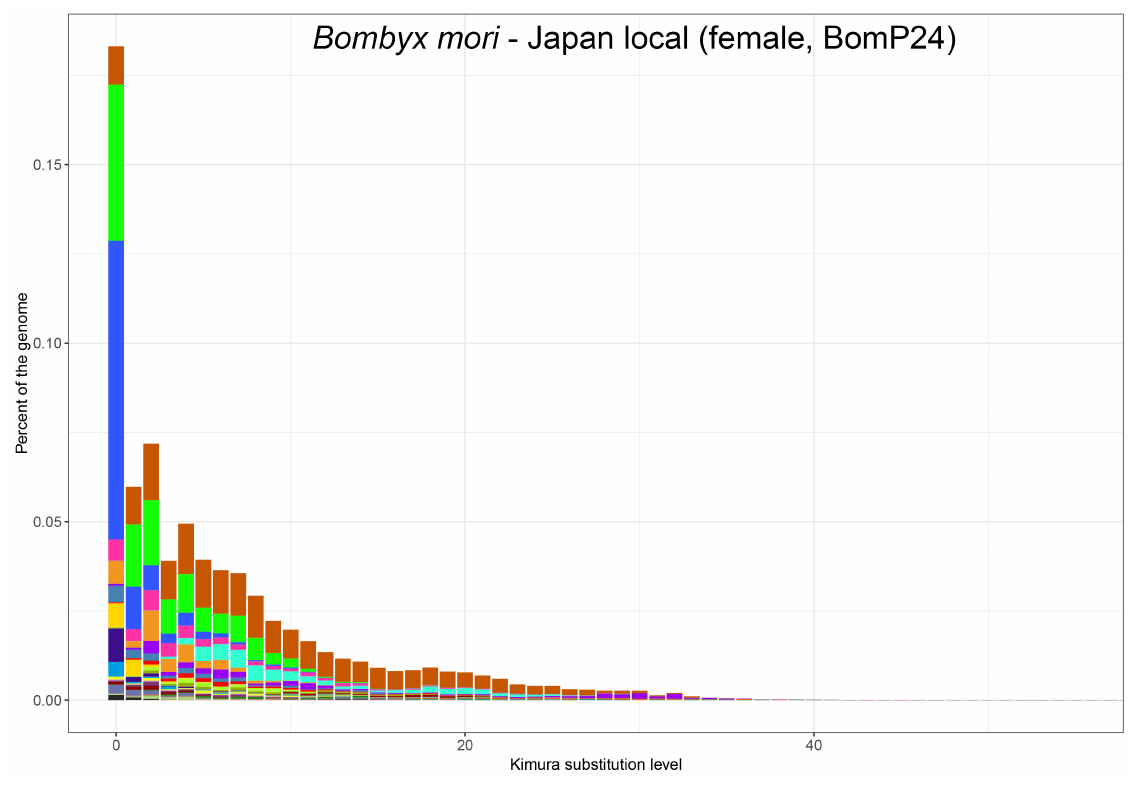

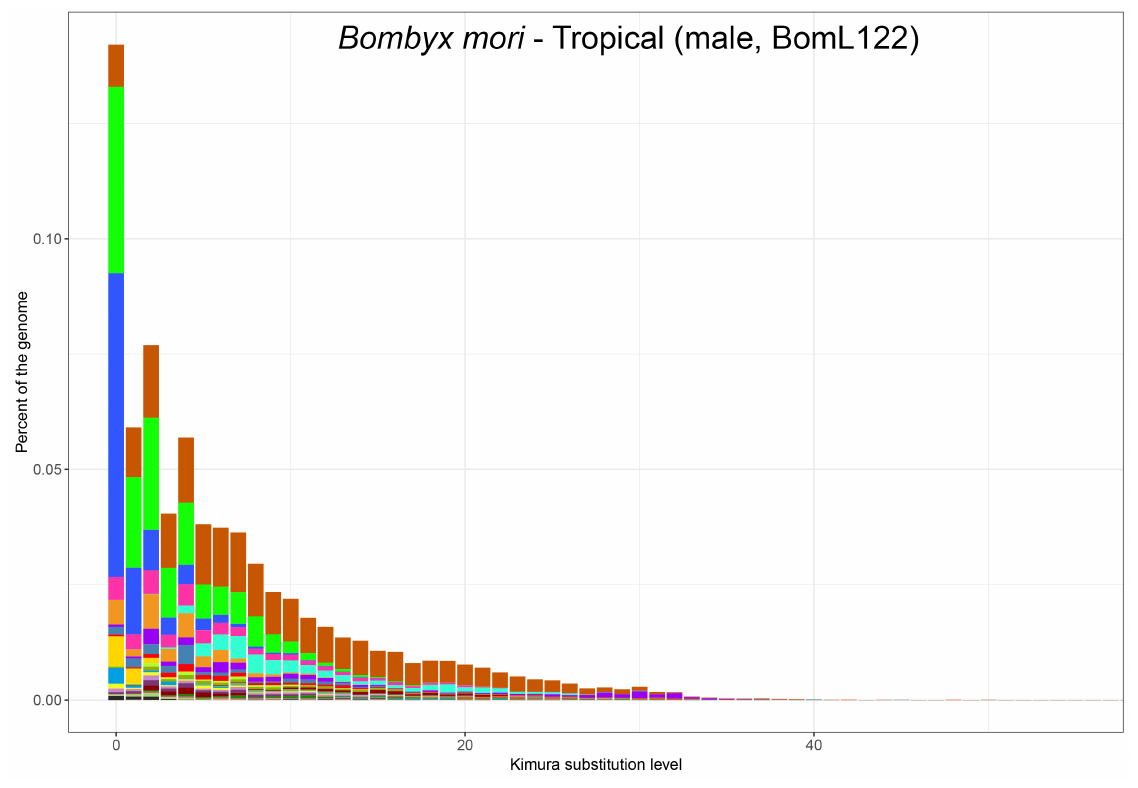

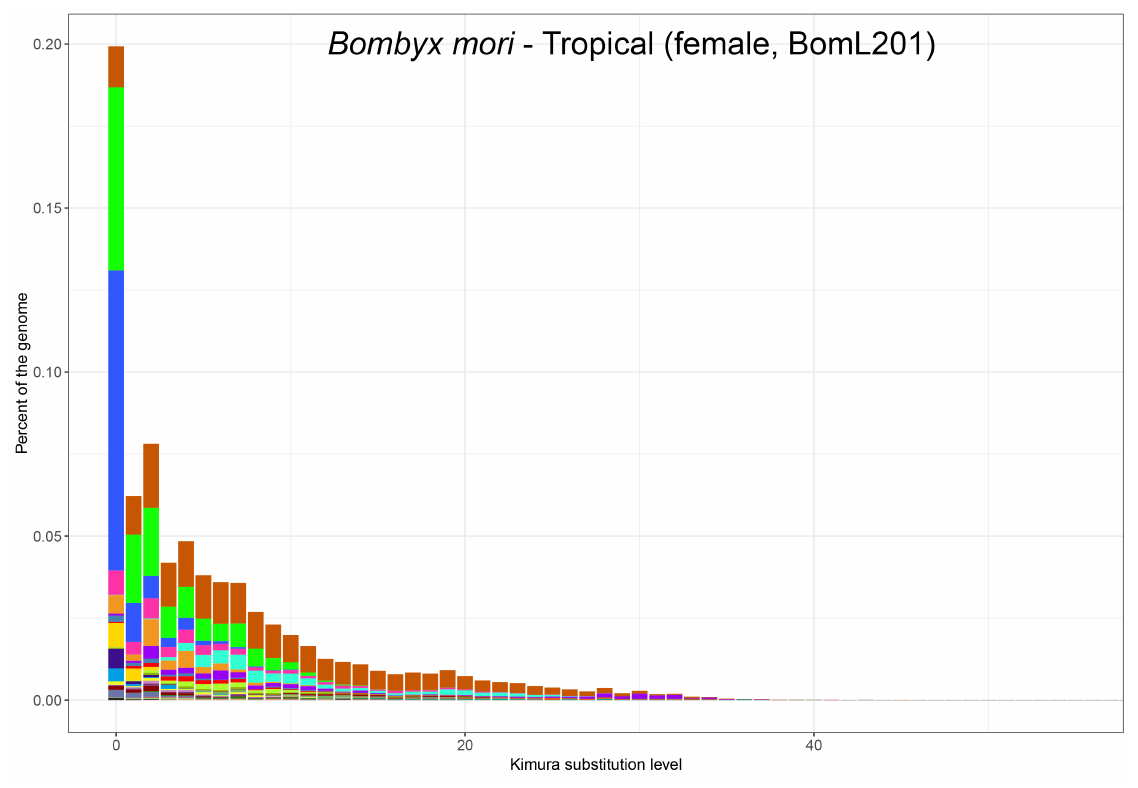

### Supp fig 2

**Supplementary Figure 2.**

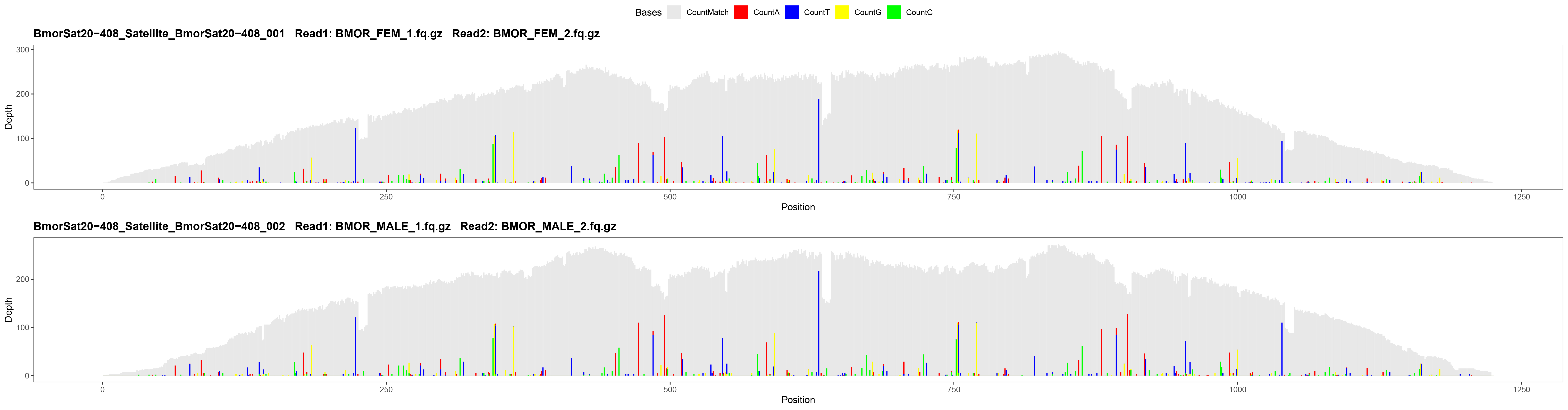

### Supp fig 3

**Supplementary Figure 3.**

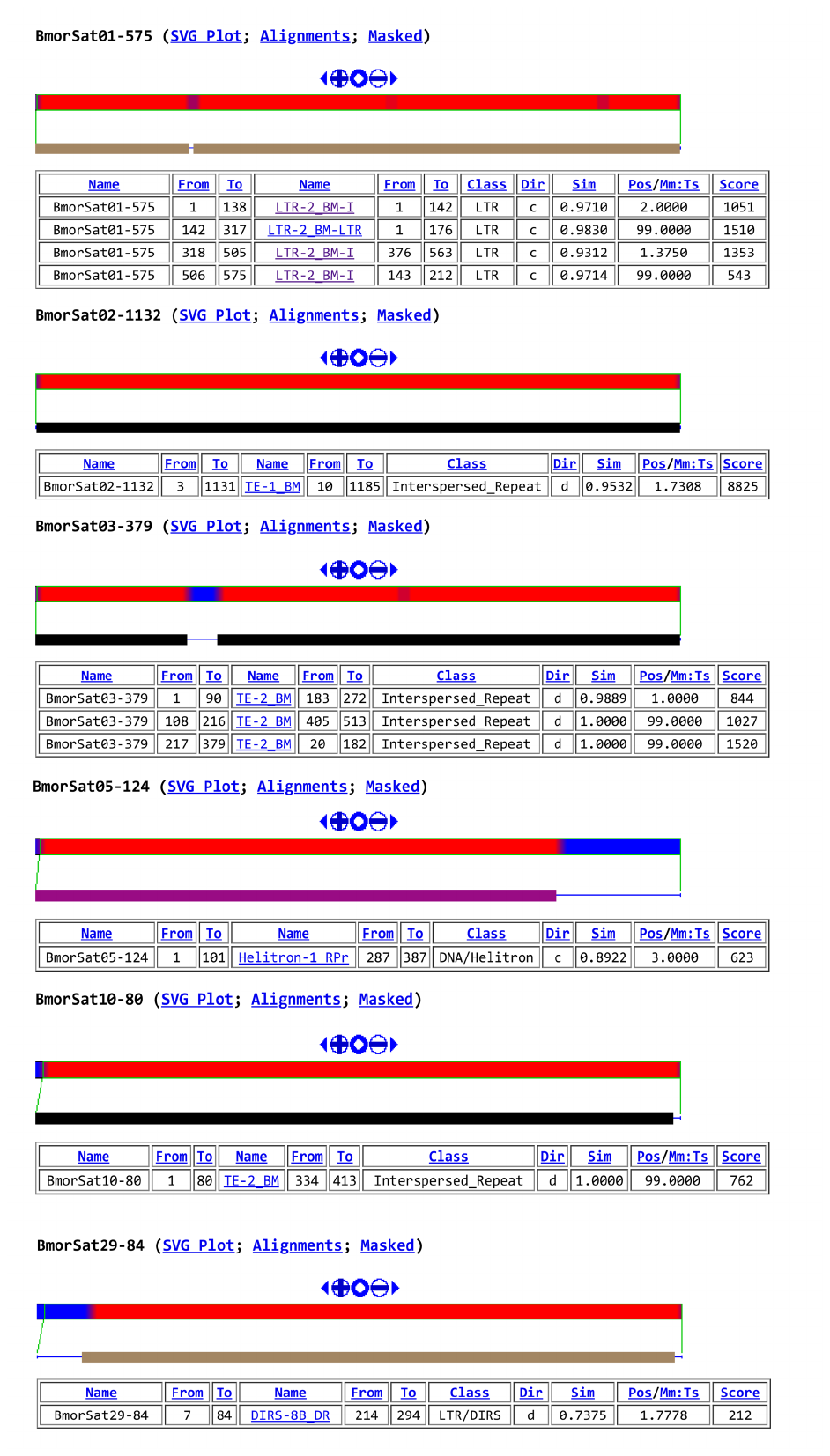
