## Supplementary material for "Advancing the understanding of the *Bombyx mori* genome through satellite DNA analysis: low strain differentiation and relationship with transposable elements": Supp fig 4

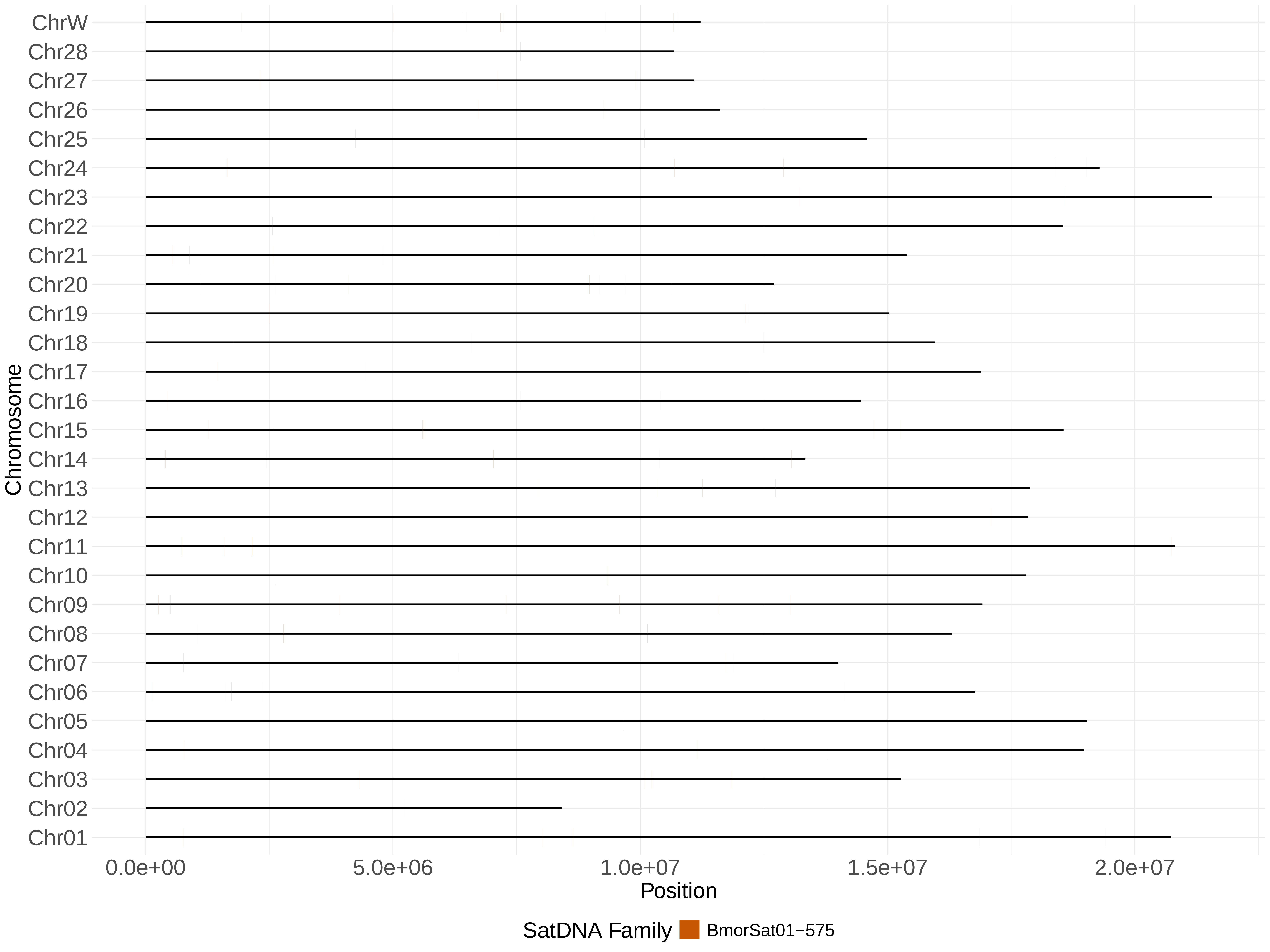

ChrW
Chr28
Chr27
Chr26
Chr25
Chr24
Chr23
Chr22
Chr21
Chr20
Chr19
Chr18
Chr17
Chr16
Chr15
Chr14
Chr13
Chr12
Chr11
Chr10
Chr09
Chr08
Chr07
Chr06
Chr05
Chr04
Chr03
Chr02
Chr01
Chromosome
0.0e+00
5.0e+06
1.0e+07
1.5e+07
2.0e+07
Position
SatDNA Family
BmorSat01−575

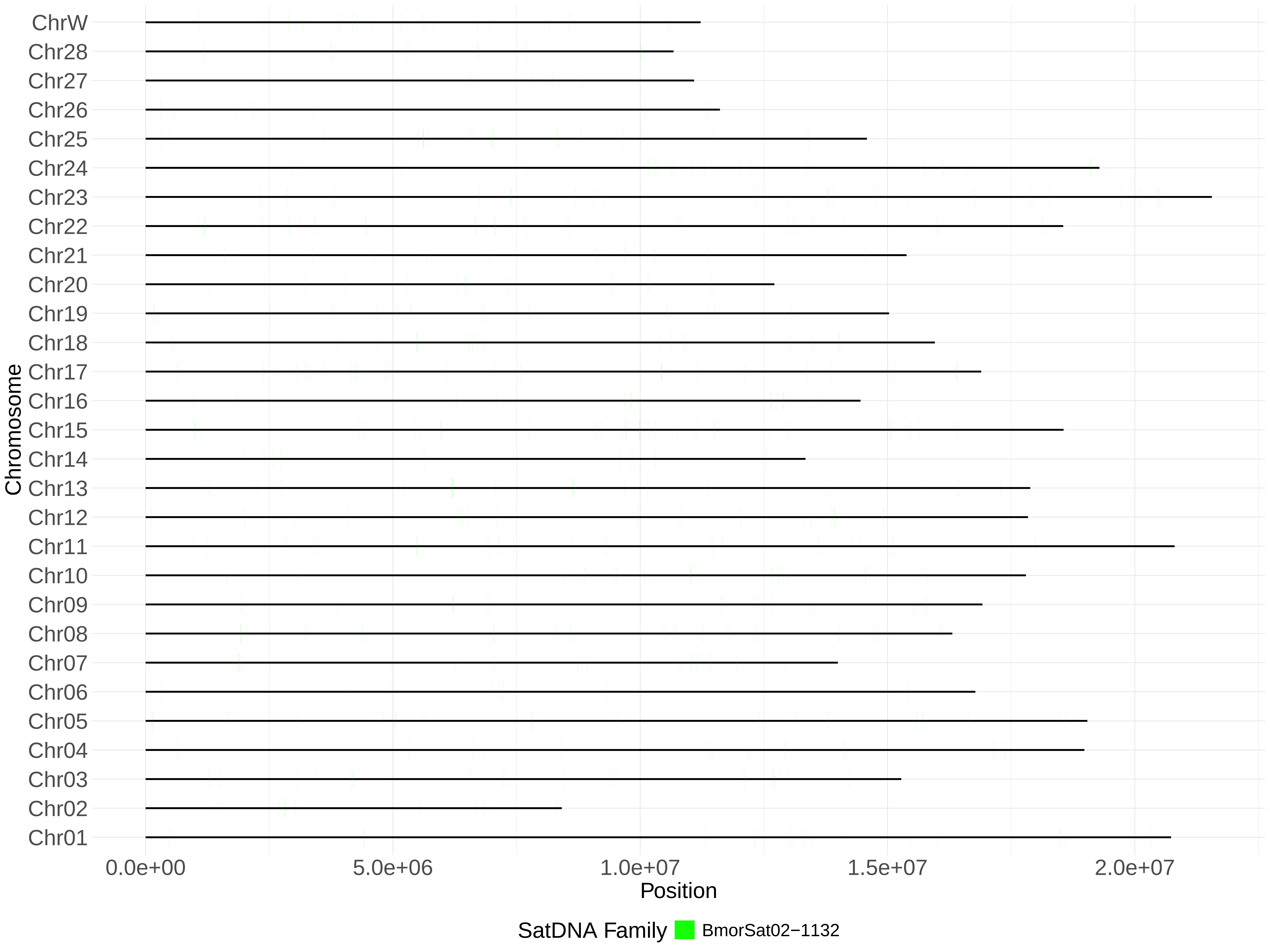

ChrW
Chr28
Chr27
Chr26
Chr25
Chr24
Chr23
Chr22
Chr21
Chr20
Chr19
Chr18
Chr17
Chr16
Chr15
Chr14
Chr13
Chr12
Chr11
Chr10
Chr09
Chr08
Chr07
Chr06
Chr05
Chr04
Chr03
Chr02
Chr01
Chromosome
0.0e+00
5.0e+06
1.0e+07
1.5e+07
2.0e+07
Position
SatDNA Family
BmorSat02−1132

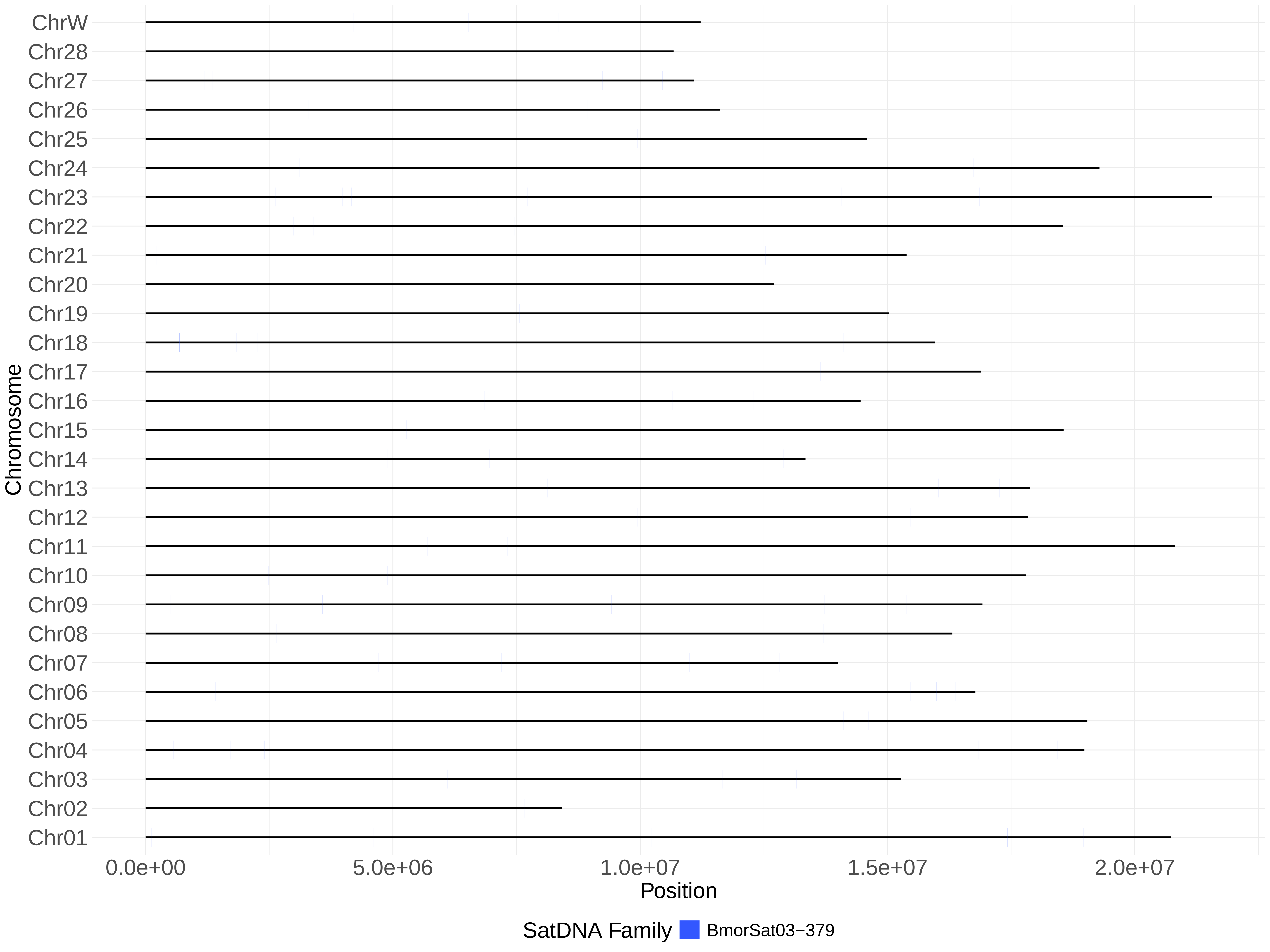
ChrW
Chr28
Chr27
Chr26
Chr25
Chr24
Chr23
Chr22
Chr21
Chr20
Chr19
Chr18
Chr17
Chr16
Chr15
Chr14
Chr13
Chr12
Chr11
Chr10
Chr09
Chr08
Chr07
Chr06
Chr05
Chr04
Chr03
Chr02
Chr01
Chromosome
0.0e+00
5.0e+06
1.0e+07
1.5e+07
2.0e+07
Position
SatDNA Family
BmorSat03−379

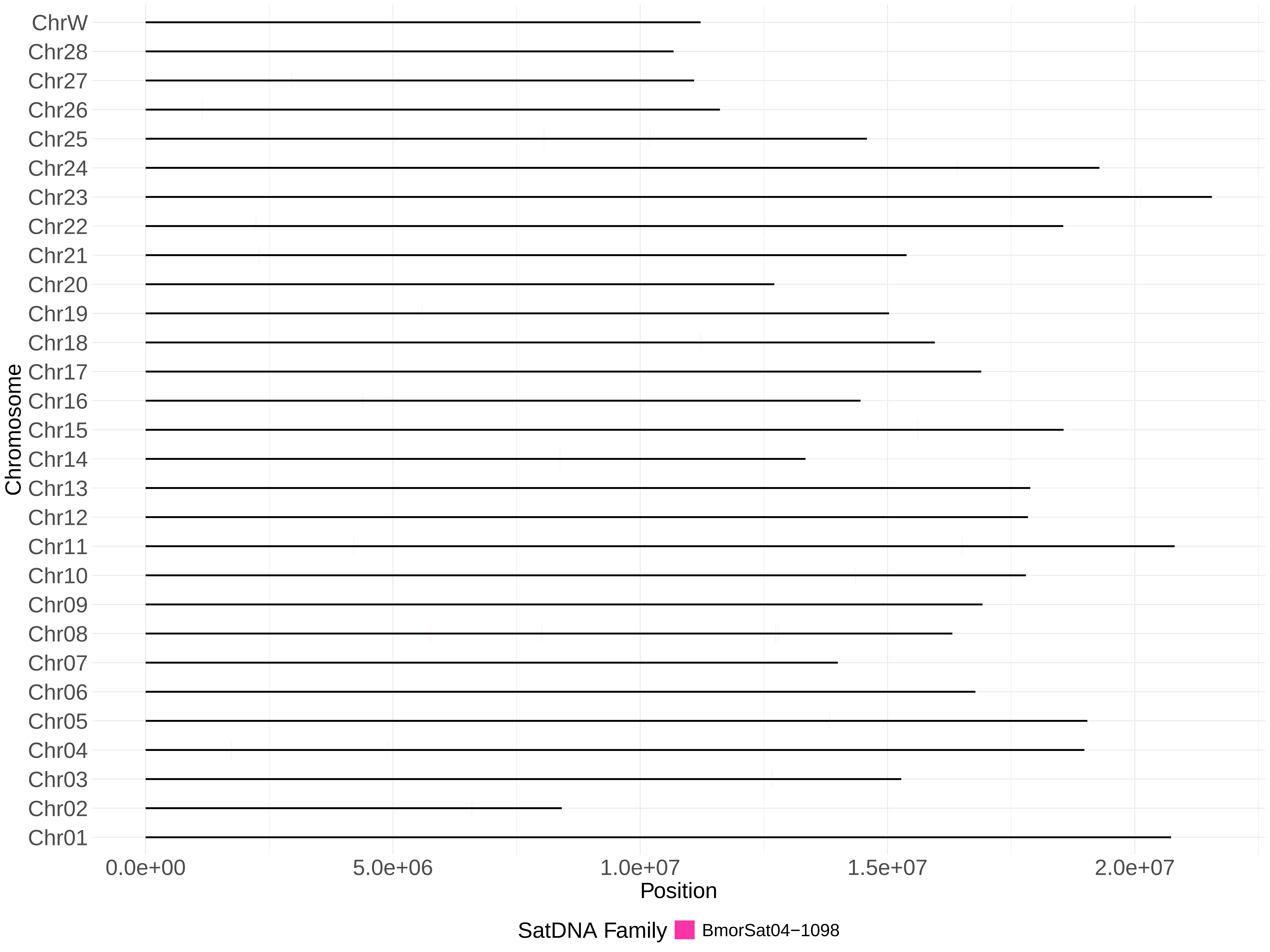

ChrW
Chr28
Chr27
Chr26
Chr25
Chr24
Chr23
Chr22
Chr21
Chr20
Chr19
Chr18
Chr17
Chr16
Chr15
Chr14
Chr13
Chr12
Chr11
Chr10
Chr09
Chr08
Chr07
Chr06
Chr05
Chr04
Chr03
Chr02
Chr01
Chromosome
0.0e+00
5.0e+06
1.0e+07
1.5e+07
2.0e+07
Position
SatDNA Family
BmorSat04−1098

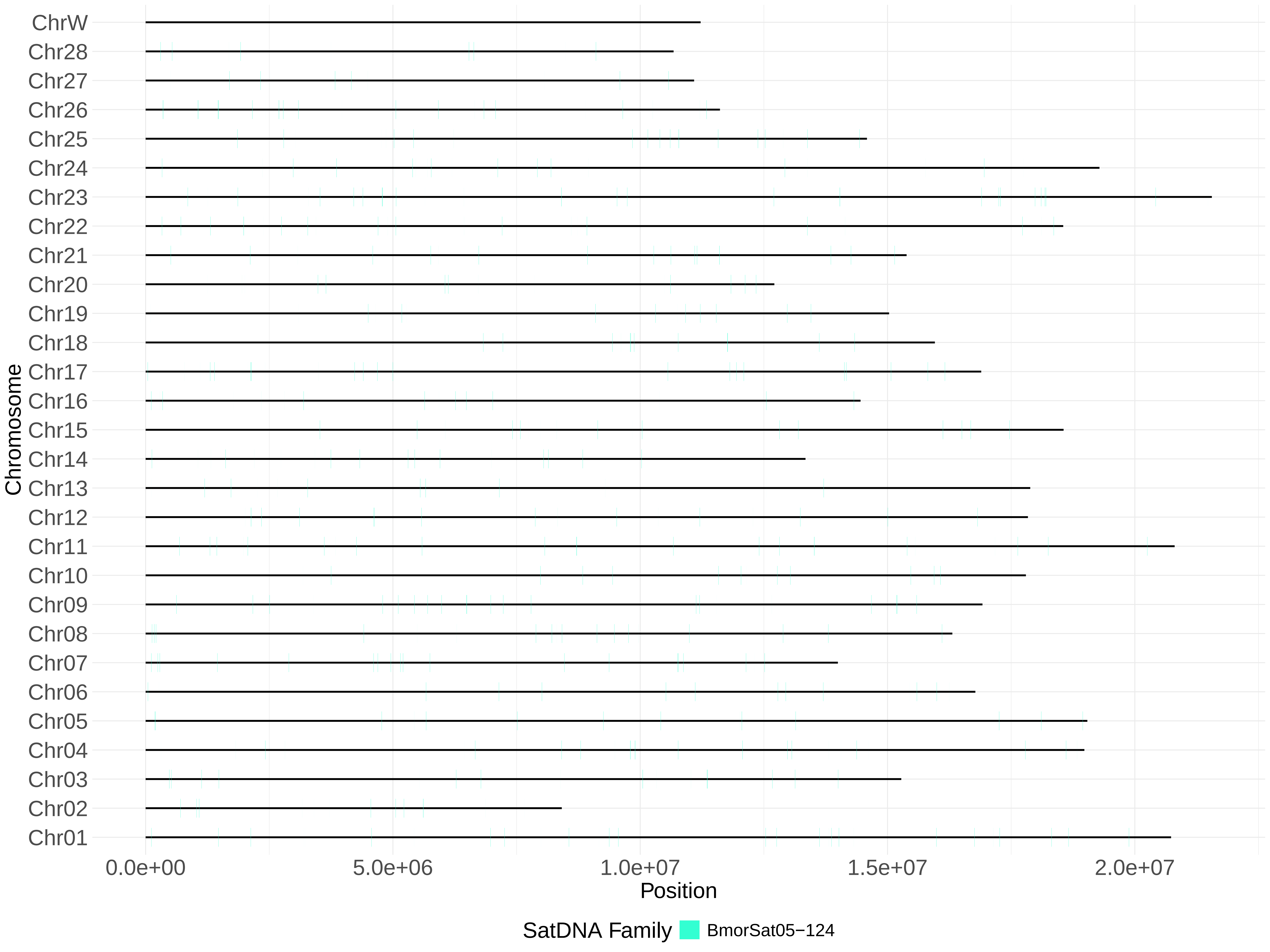
ChrW
Chr28
Chr27
Chr26
Chr25
Chr24
Chr23
Chr22
Chr21
Chr20
Chr19
Chr18
Chr17
Chr16
Chr15
Chr14
Chr13
Chr12
Chr11
Chr10
Chr09
Chr08
Chr07
Chr06
Chr05
Chr04
Chr03
Chr02
Chr01
Chromosome
0.0e+00
5.0e+06
1.0e+07
1.5e+07
2.0e+07
Position
SatDNA Family
BmorSat05−124

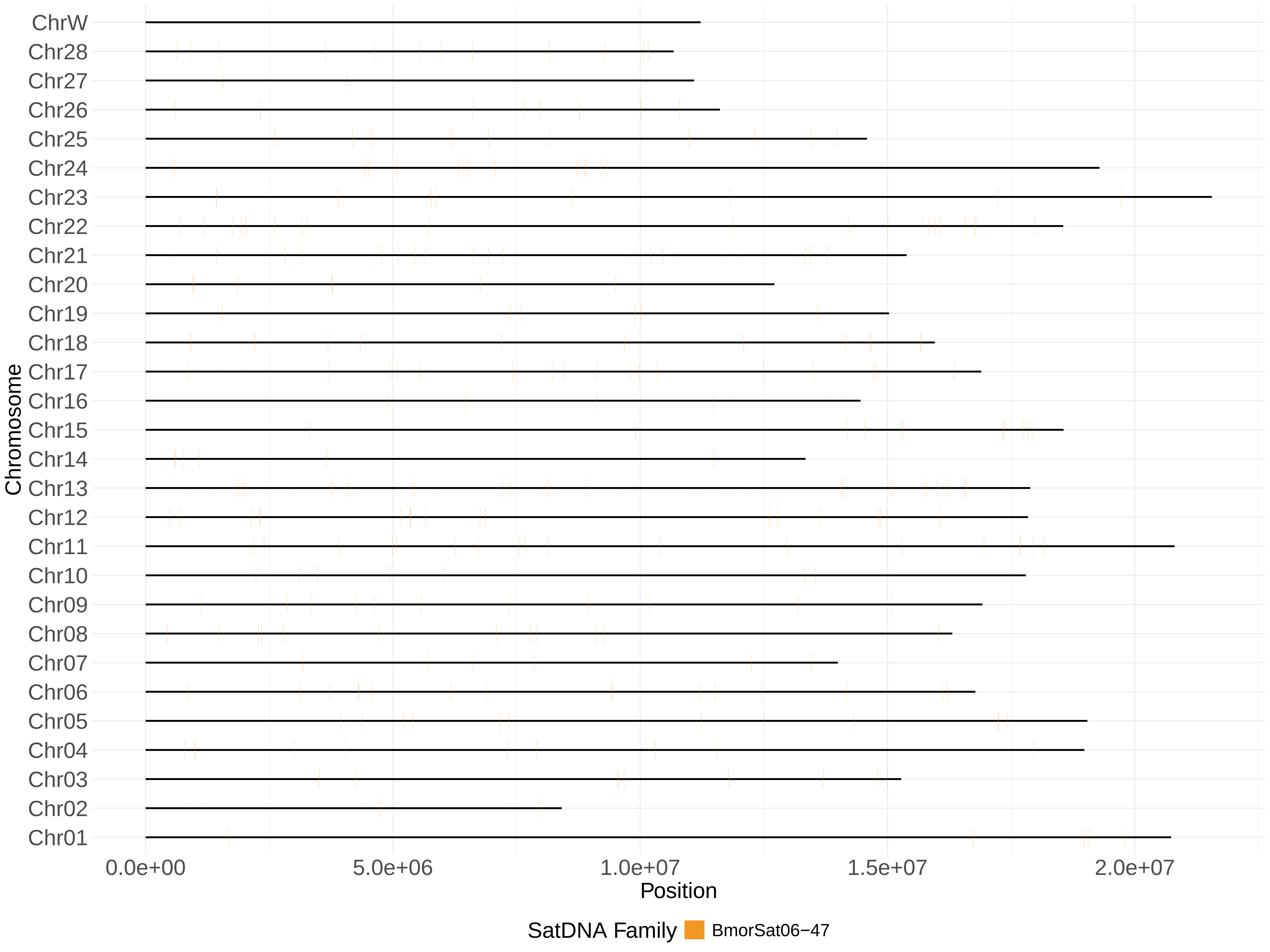

ChrW
Chr28
Chr27
Chr26
Chr25
Chr24
Chr23
Chr22
Chr21
Chr20
Chr19
Chr18
Chr17
Chr16
Chr15
Chr14
Chr13
Chr12
Chr11
Chr10
Chr09
Chr08
Chr07
Chr06
Chr05
Chr04
Chr03
Chr02
Chr01
Chromosome
0.0e+00
5.0e+06
1.0e+07
1.5e+07
2.0e+07
Position
SatDNA Family
BmorSat06−47

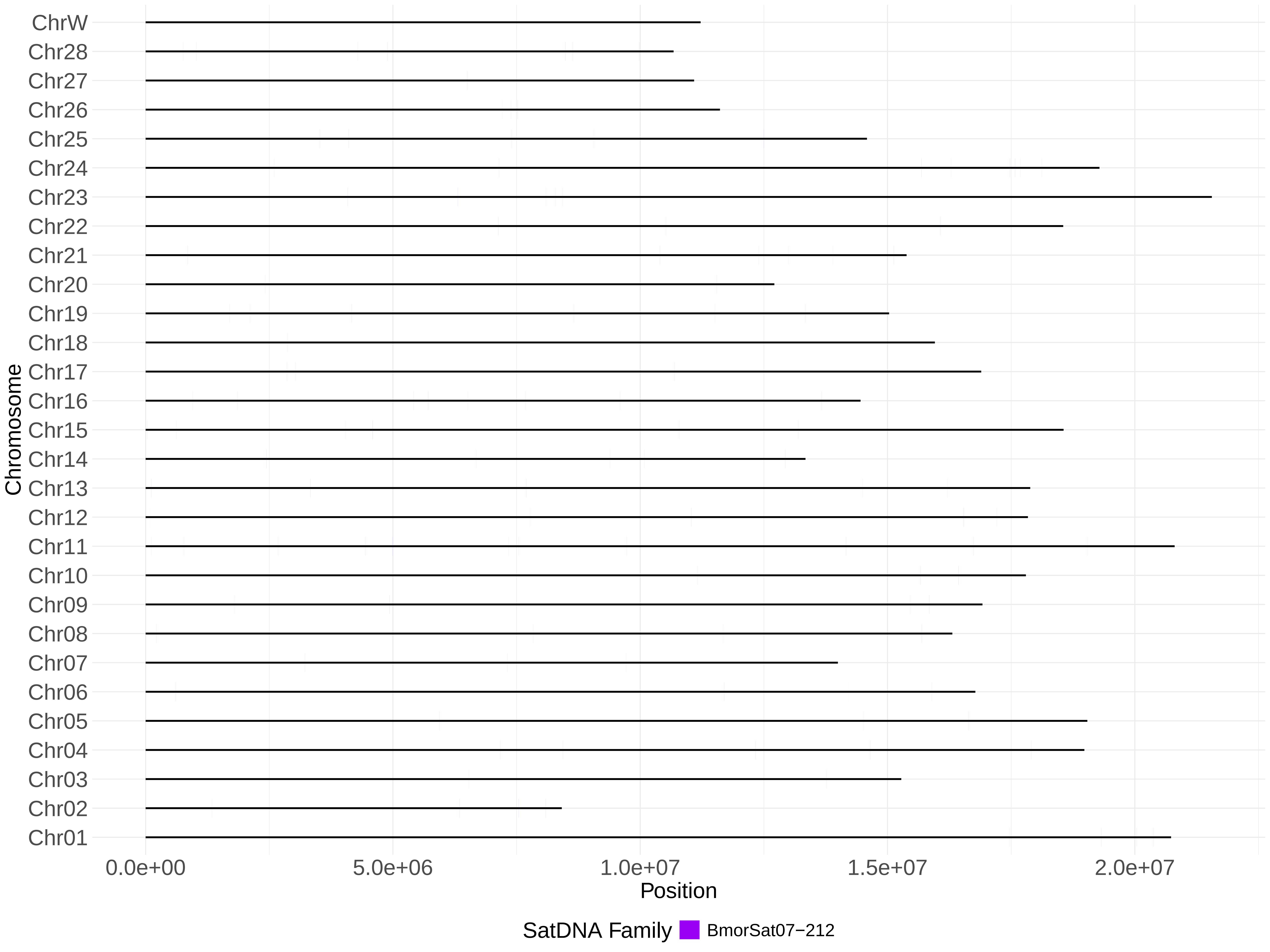
ChrW
Chr28
Chr27
Chr26
Chr25
Chr24
Chr23
Chr22
Chr21
Chr20
Chr19
Chr18
Chr17
Chr16
Chr15
Chr14
Chr13
Chr12
Chr11
Chr10
Chr09
Chr08
Chr07
Chr06
Chr05
Chr04
Chr03
Chr02
Chr01
Chromosome
0.0e+00
5.0e+06
1.0e+07
1.5e+07
2.0e+07
Position
SatDNA Family
BmorSat07−212

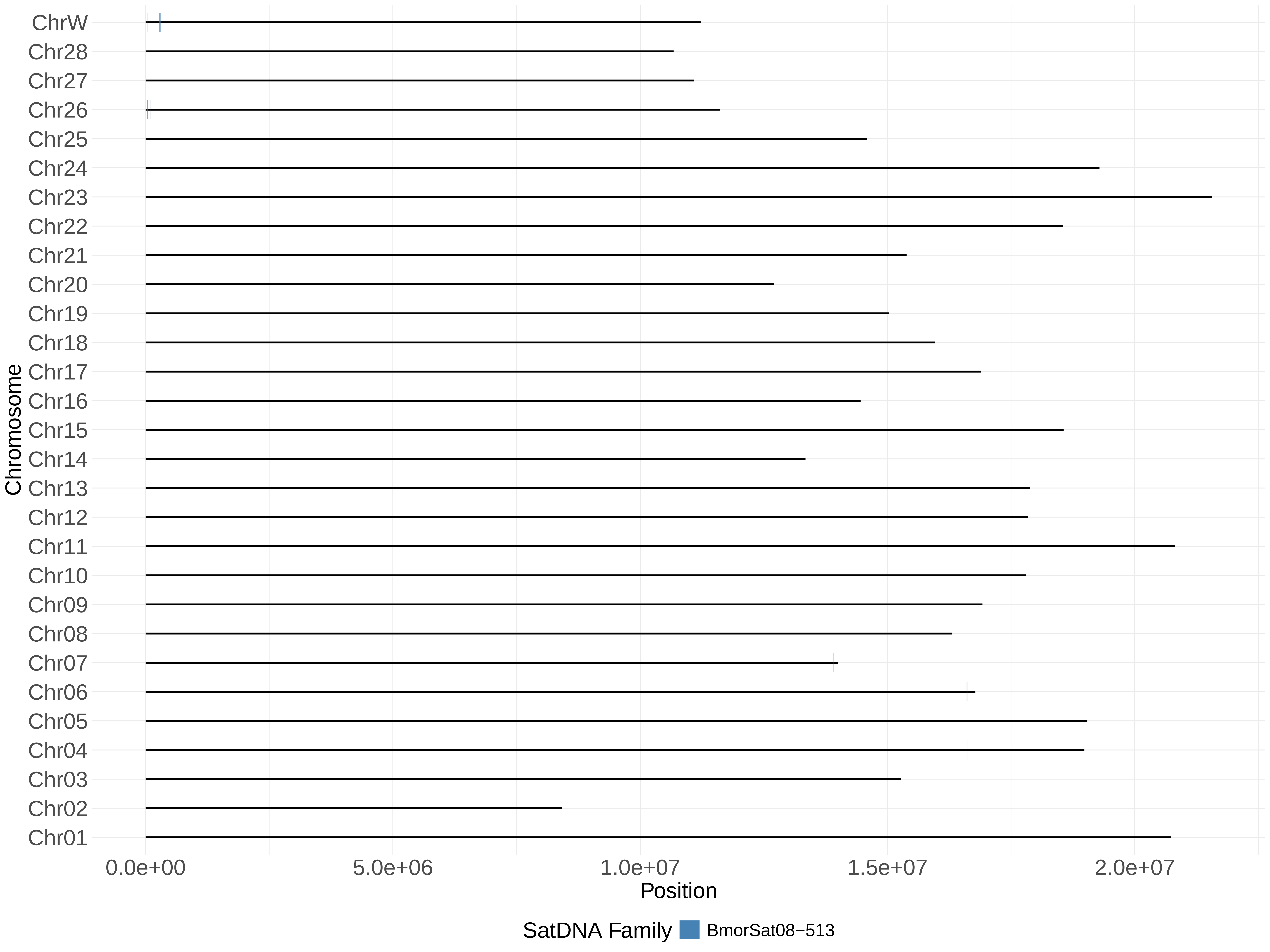

ChrW
Chr28
Chr27
Chr26
Chr25
Chr24
Chr23
Chr22
Chr21
Chr20
Chr19
Chr18
Chr17
Chr16
Chr15
Chr14
Chr13
Chr12
Chr11
Chr10
Chr09
Chr08
Chr07
Chr06
Chr05
Chr04
Chr03
Chr02
Chr01
Chromosome
0.0e+00
5.0e+06
1.0e+07
1.5e+07
2.0e+07
Position
SatDNA Family
BmorSat08−513

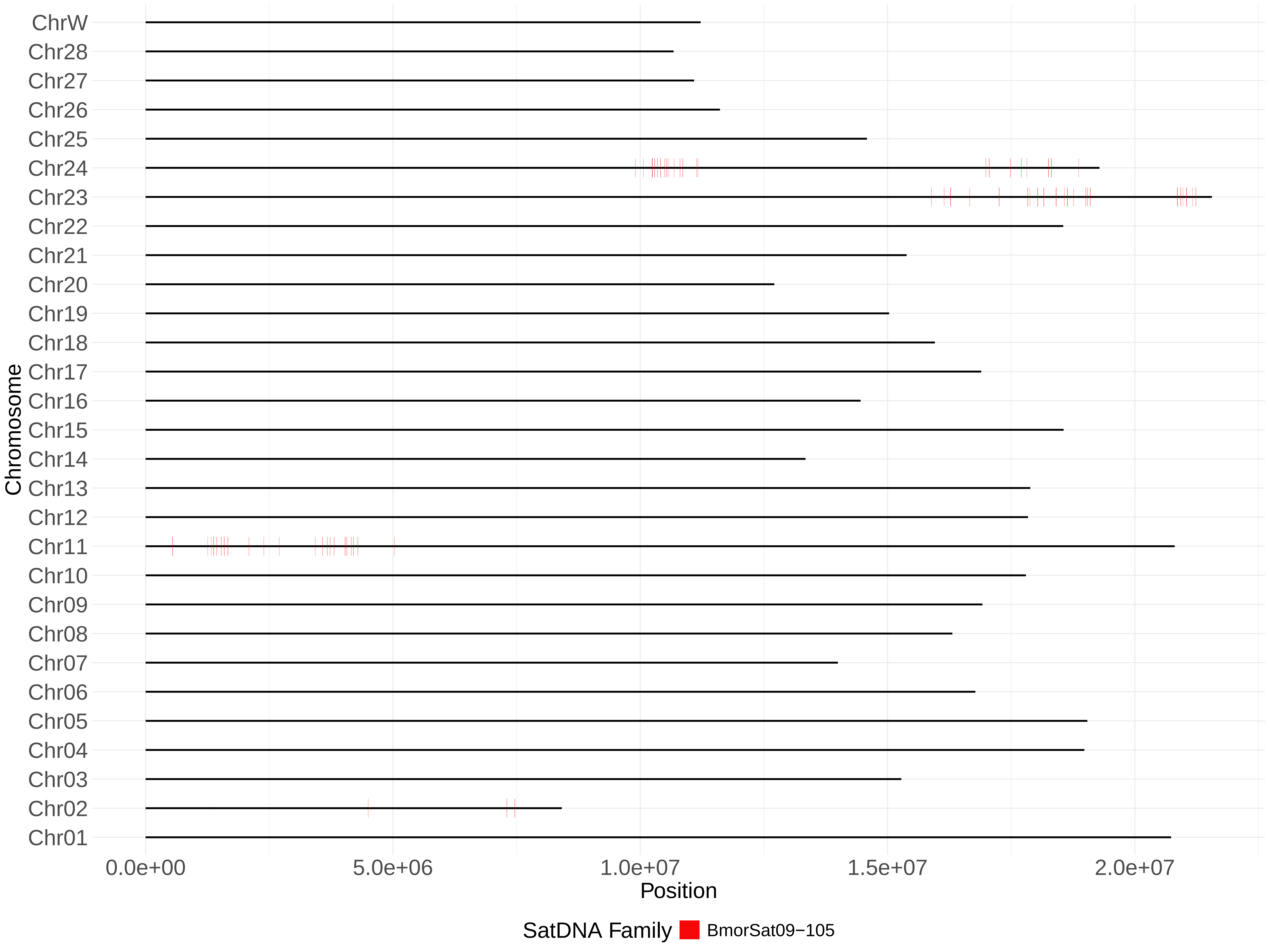

ChrW
Chr28
Chr27
Chr26
Chr25
Chr24
Chr23
Chr22
Chr21
Chr20
Chr19
Chr18
Chr17
Chr16
Chr15
Chr14
Chr13
Chr12
Chr11
Chr10
Chr09
Chr08
Chr07
Chr06
Chr05
Chr04
Chr03
Chr02
Chr01
Chromosome
0.0e+00
5.0e+06
1.0e+07
1.5e+07
2.0e+07
Position
SatDNA Family
BmorSat09−105

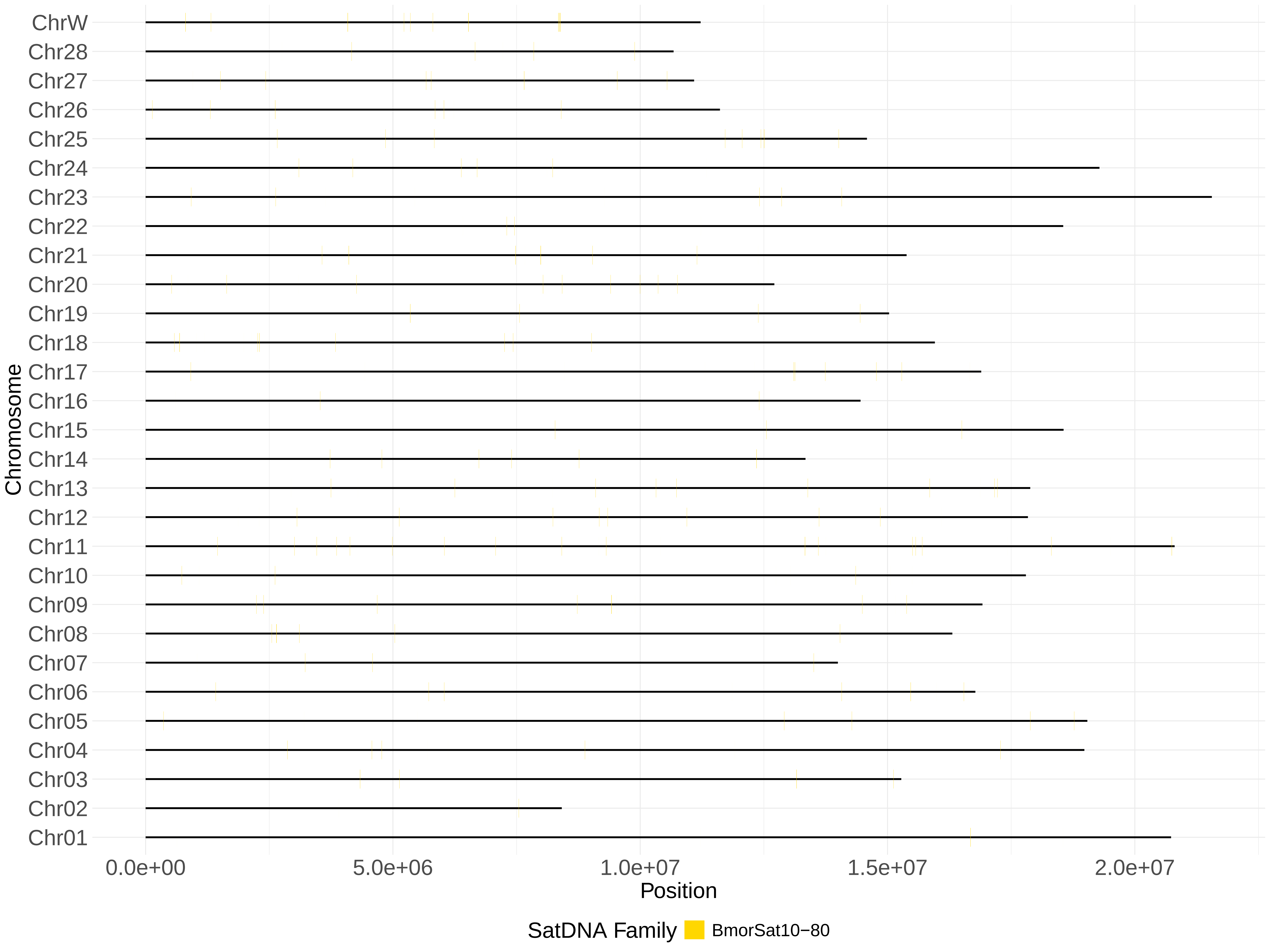

ChrW
Chr28
Chr27
Chr26
Chr25
Chr24
Chr23
Chr22
Chr21
Chr20
Chr19
Chr18
Chr17
Chr16
Chr15
Chr14
Chr13
Chr12
Chr11
Chr10
Chr09
Chr08
Chr07
Chr06
Chr05
Chr04
Chr03
Chr02
Chr01
Chromosome
0.0e+00
5.0e+06
1.0e+07
1.5e+07
2.0e+07
Position
SatDNA Family
BmorSat10−80

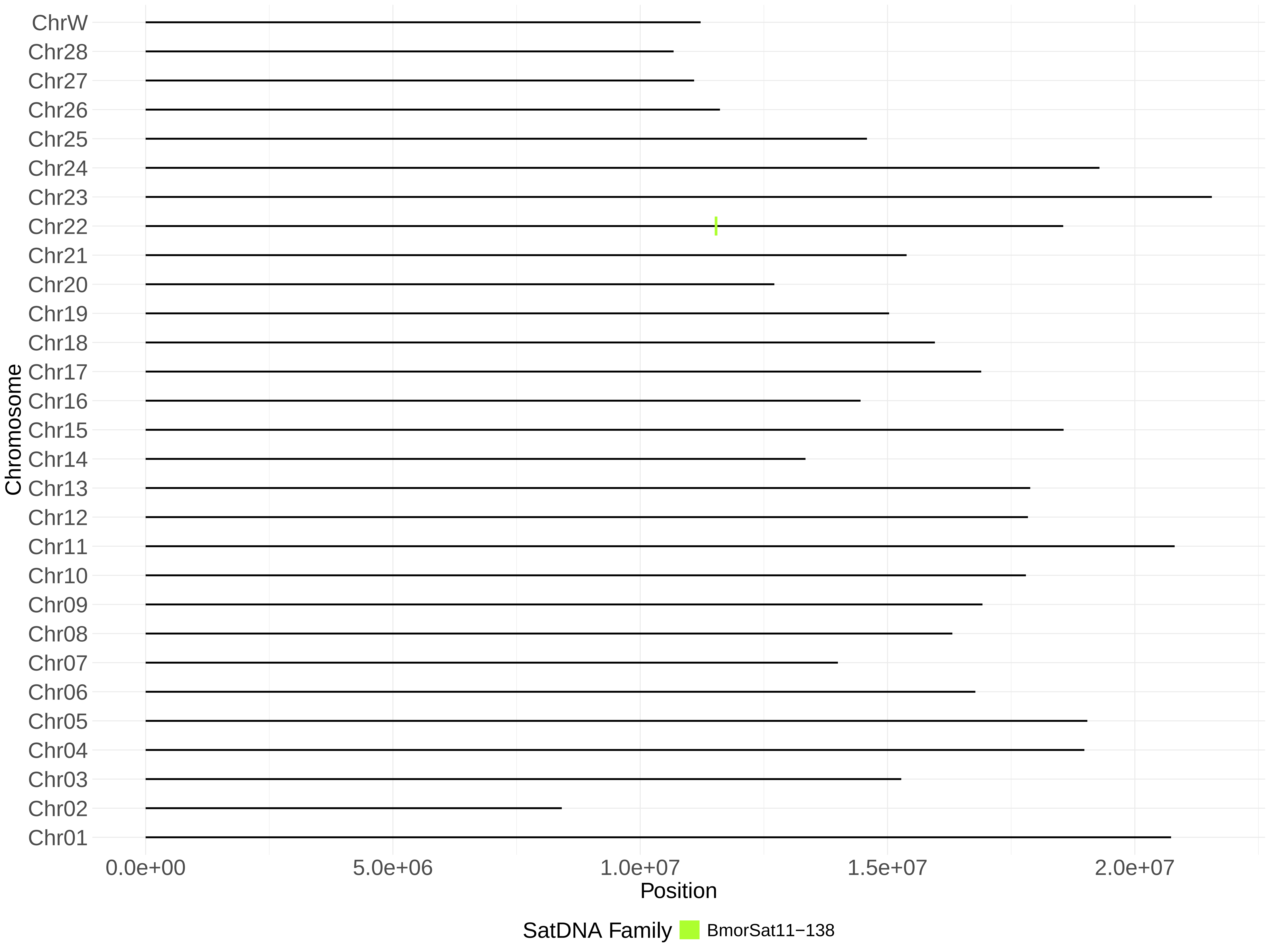

ChrW
Chr28
Chr27
Chr26
Chr25
Chr24
Chr23
Chr22
Chr21
Chr20
Chr19
Chr18
Chr17
Chr16
Chr15
Chr14
Chr13
Chr12
Chr11
Chr10
Chr09
Chr08
Chr07
Chr06
Chr05
Chr04
Chr03
Chr02
Chr01
Chromosome
0.0e+00
5.0e+06
1.0e+07
1.5e+07
2.0e+07
Position
SatDNA Family
BmorSat11−138

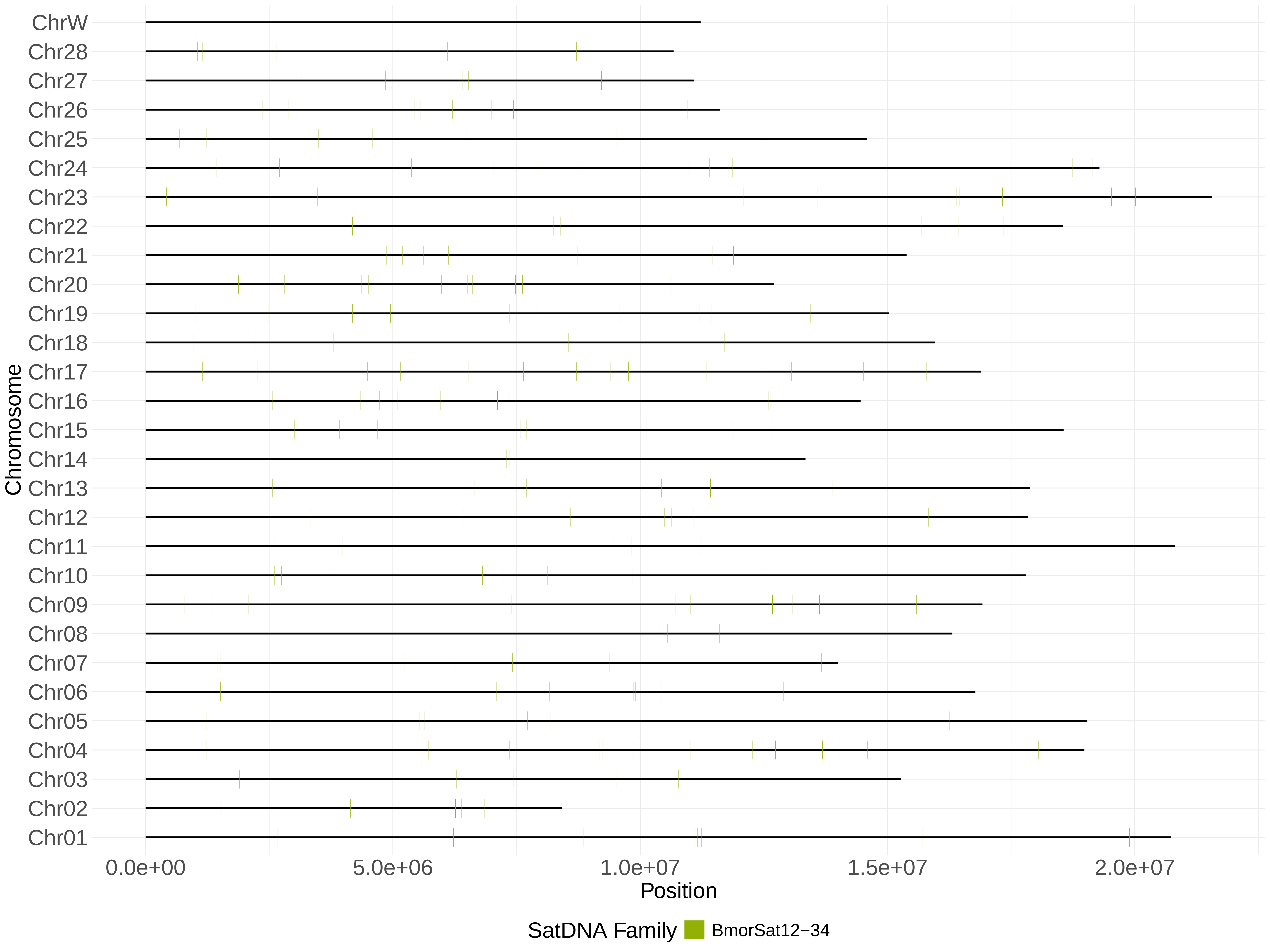
ChrW
Chr28
Chr27
Chr26
Chr25
Chr24
Chr23
Chr22
Chr21
Chr20
Chr19
Chr18
Chr17
Chr16
Chr15
Chr14
Chr13
Chr12
Chr11
Chr10
Chr09
Chr08
Chr07
Chr06
Chr05
Chr04
Chr03
Chr02
Chr01
Chromosome
0.0e+00
5.0e+06
1.0e+07
1.5e+07
2.0e+07
Position
SatDNA Family
BmorSat12−34

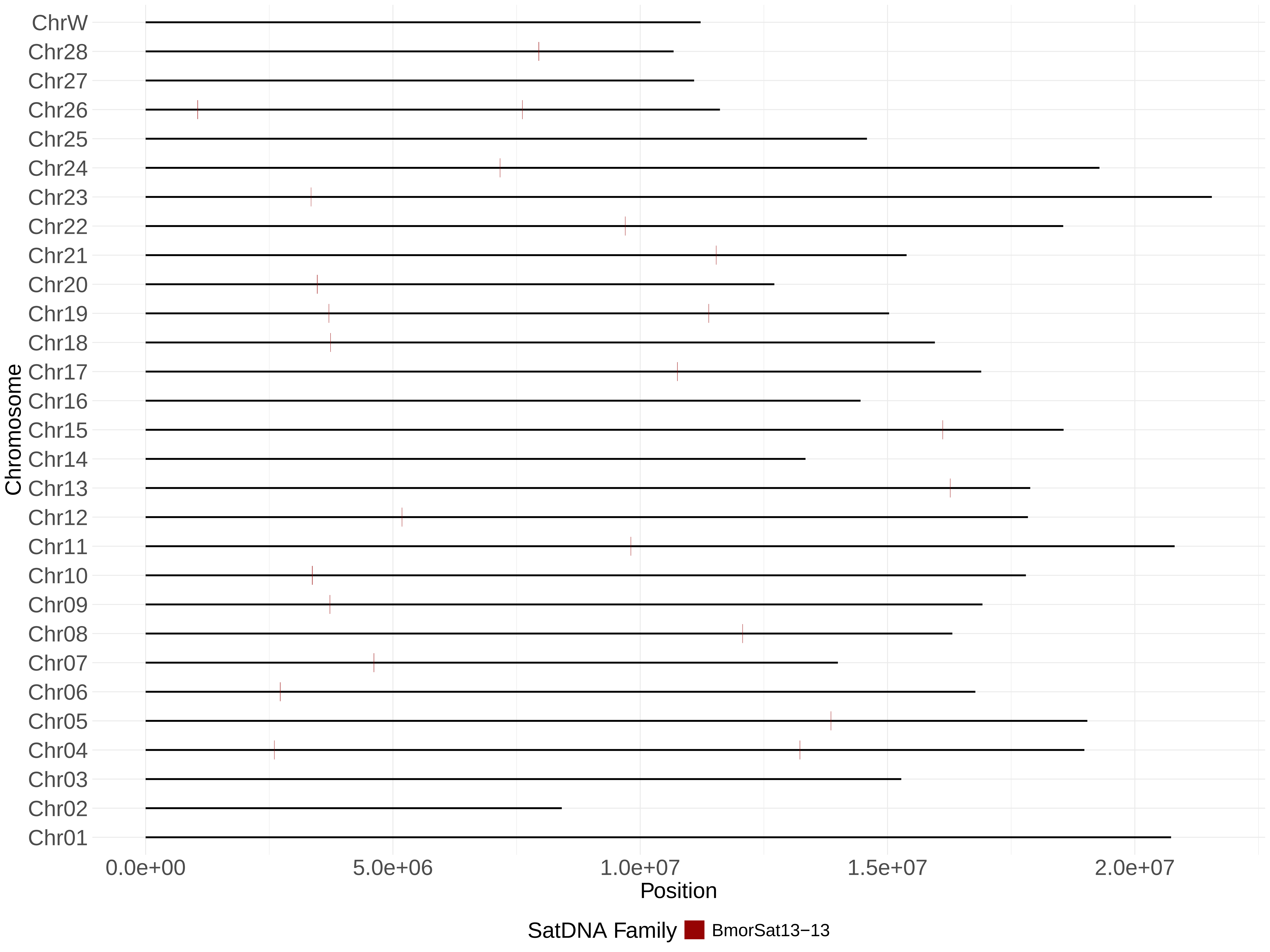

ChrW
Chr28
Chr27
Chr26
Chr25
Chr24
Chr23
Chr22
Chr21
Chr20
Chr19
Chr18
Chr17
Chr16
Chr15
Chr14
Chr13
Chr12
Chr11
Chr10
Chr09
Chr08
Chr07
Chr06
Chr05
Chr04
Chr03
Chr02
Chr01
Chromosome
0.0e+00
5.0e+06
1.0e+07
1.5e+07
2.0e+07
Position
SatDNA Family
BmorSat13−13

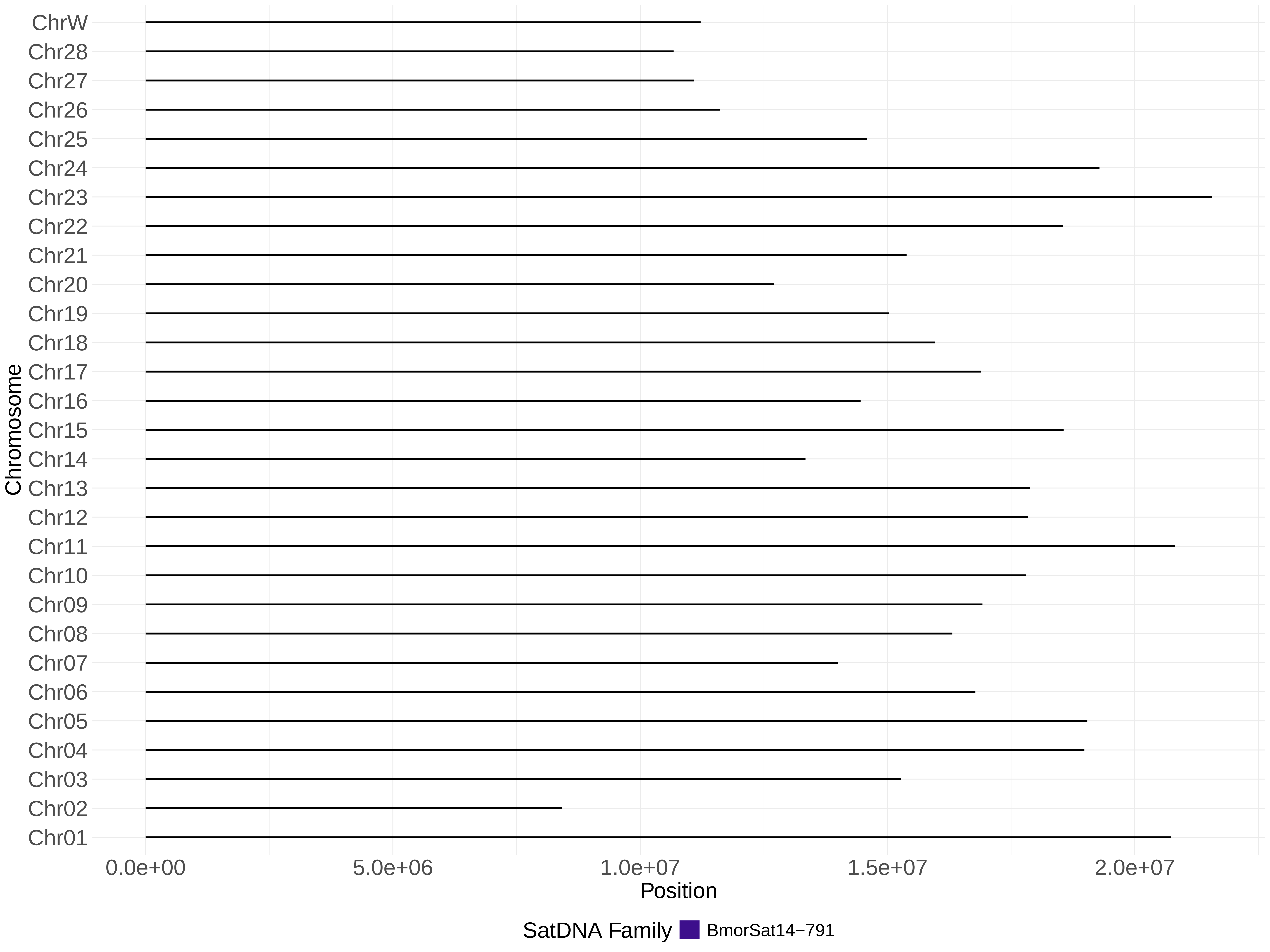

ChrW
Chr28
Chr27
Chr26
Chr25
Chr24
Chr23
Chr22
Chr21
Chr20
Chr19
Chr18
Chr17
Chr16
Chr15
Chr14
Chr13
Chr12
Chr11
Chr10
Chr09
Chr08
Chr07
Chr06
Chr05
Chr04
Chr03
Chr02
Chr01
Chromosome
0.0e+00
5.0e+06
1.0e+07
1.5e+07
2.0e+07
Position
SatDNA Family
BmorSat14−791

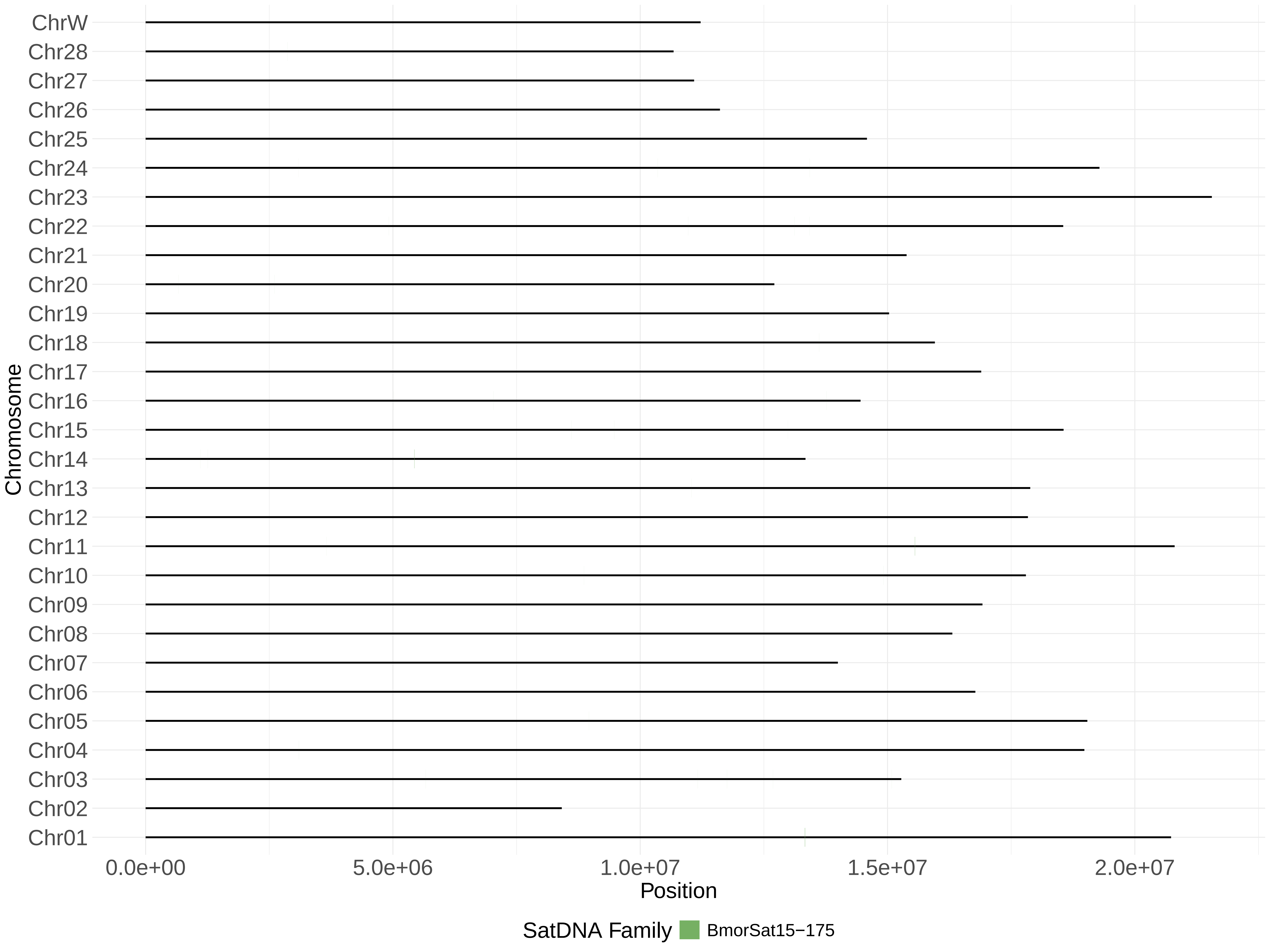
ChrW
Chr28
Chr27
Chr26
Chr25
Chr24
Chr23
Chr22
Chr21
Chr20
Chr19
Chr18
Chr17
Chr16
Chr15
Chr14
Chr13
Chr12
Chr11
Chr10
Chr09
Chr08
Chr07
Chr06
Chr05
Chr04
Chr03
Chr02
Chr01
Chromosome
0.0e+00
5.0e+06
1.0e+07
1.5e+07
2.0e+07
Position
SatDNA Family
BmorSat15−175

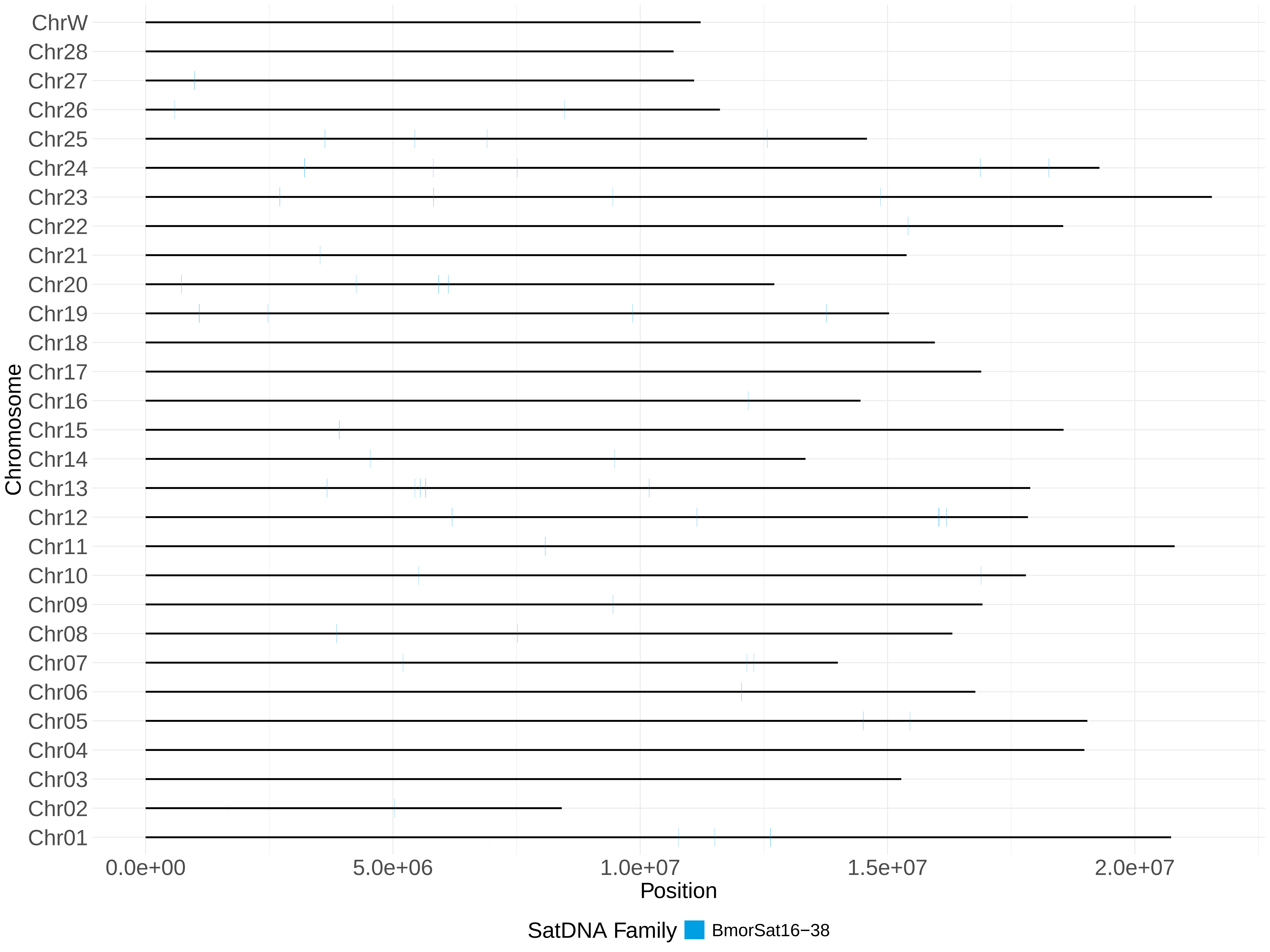

ChrW
Chr28
Chr27
Chr26
Chr25
Chr24
Chr23
Chr22
Chr21
Chr20
Chr19
Chr18
Chr17
Chr16
Chr15
Chr14
Chr13
Chr12
Chr11
Chr10
Chr09
Chr08
Chr07
Chr06
Chr05
Chr04
Chr03
Chr02
Chr01
Chromosome
0.0e+00
5.0e+06
1.0e+07
1.5e+07
2.0e+07
Position
SatDNA Family
BmorSat16−38

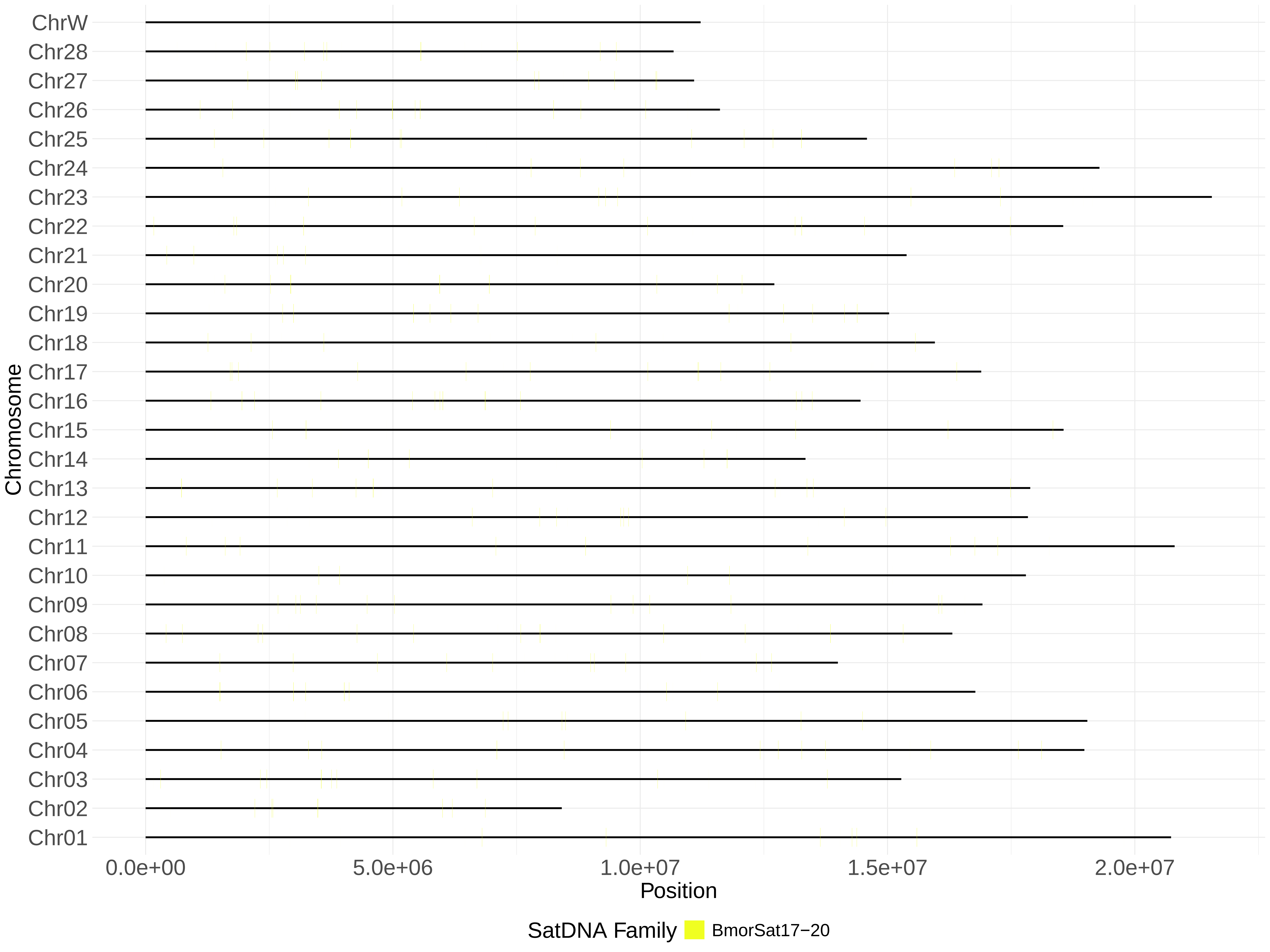

ChrW
Chr28
Chr27
Chr26
Chr25
Chr24
Chr23
Chr22
Chr21
Chr20
Chr19
Chr18
Chr17
Chr16
Chr15
Chr14
Chr13
Chr12
Chr11
Chr10
Chr09
Chr08
Chr07
Chr06
Chr05
Chr04
Chr03
Chr02
Chr01
Chromosome
0.0e+00
5.0e+06
1.0e+07
1.5e+07
2.0e+07
Position
SatDNA Family
BmorSat17−20

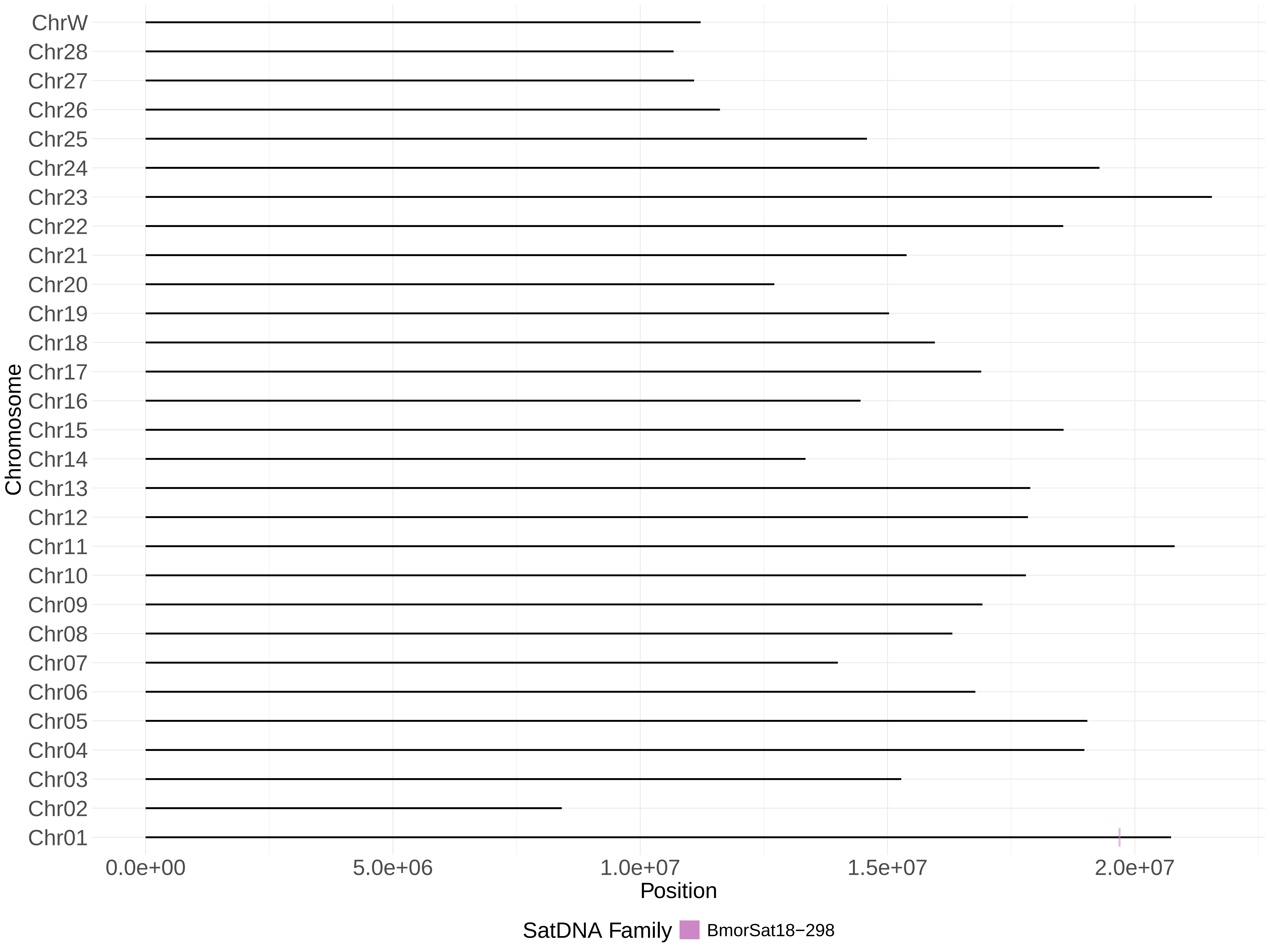
ChrW
Chr28
Chr27
Chr26
Chr25
Chr24
Chr23
Chr22
Chr21
Chr20
Chr19
Chr18
Chr17
Chr16
Chr15
Chr14
Chr13
Chr12
Chr11
Chr10
Chr09
Chr08
Chr07
Chr06
Chr05
Chr04
Chr03
Chr02
Chr01
Chromosome
0.0e+00
5.0e+06
1.0e+07
1.5e+07
2.0e+07
Position
SatDNA Family
BmorSat18−298

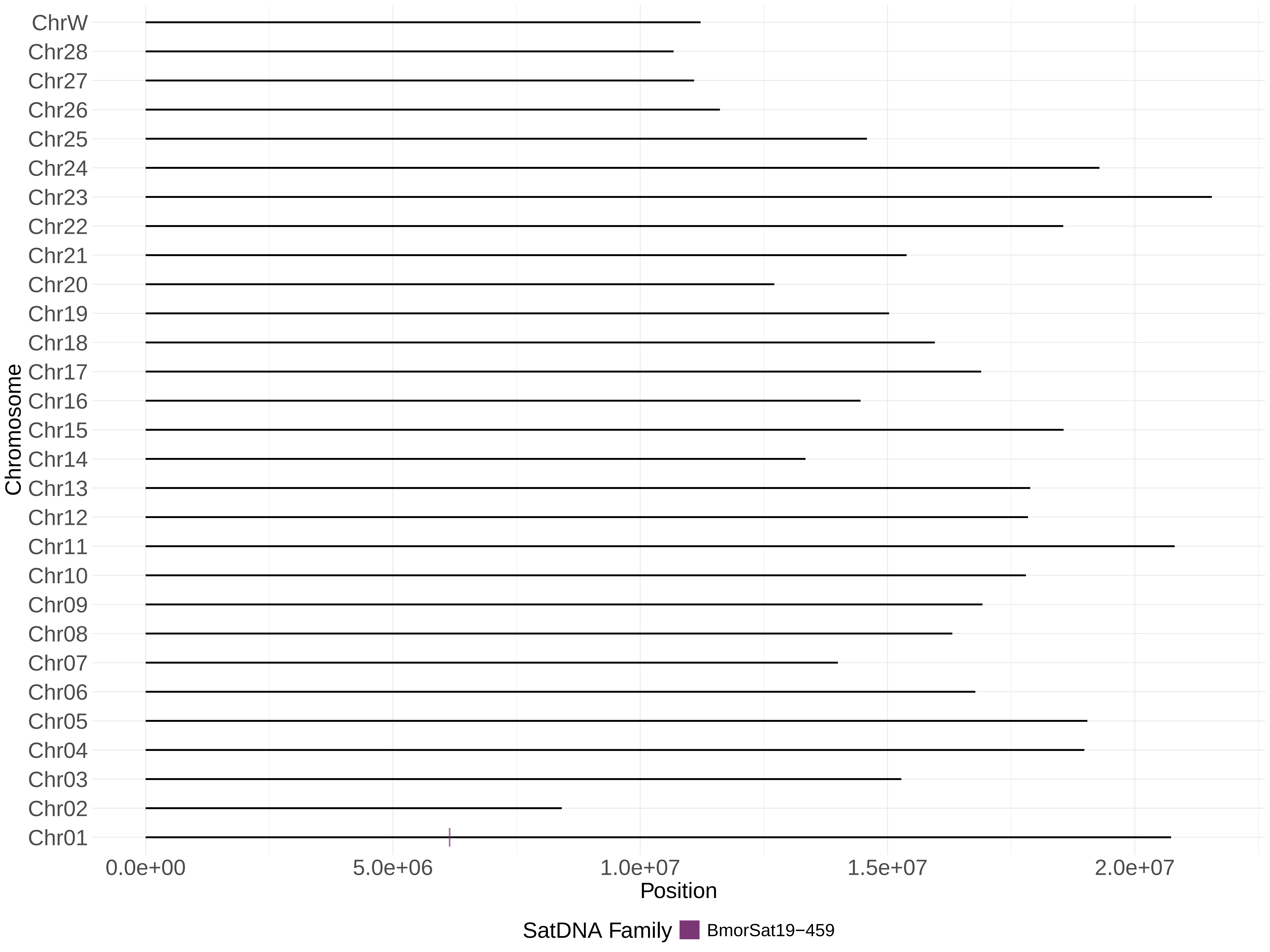

ChrW
Chr28
Chr27
Chr26
Chr25
Chr24
Chr23
Chr22
Chr21
Chr20
Chr19
Chr18
Chr17
Chr16
Chr15
Chr14
Chr13
Chr12
Chr11
Chr10
Chr09
Chr08
Chr07
Chr06
Chr05
Chr04
Chr03
Chr02
Chr01
Chromosome
0.0e+00
5.0e+06
1.0e+07
1.5e+07
2.0e+07
Position
SatDNA Family
BmorSat19−459

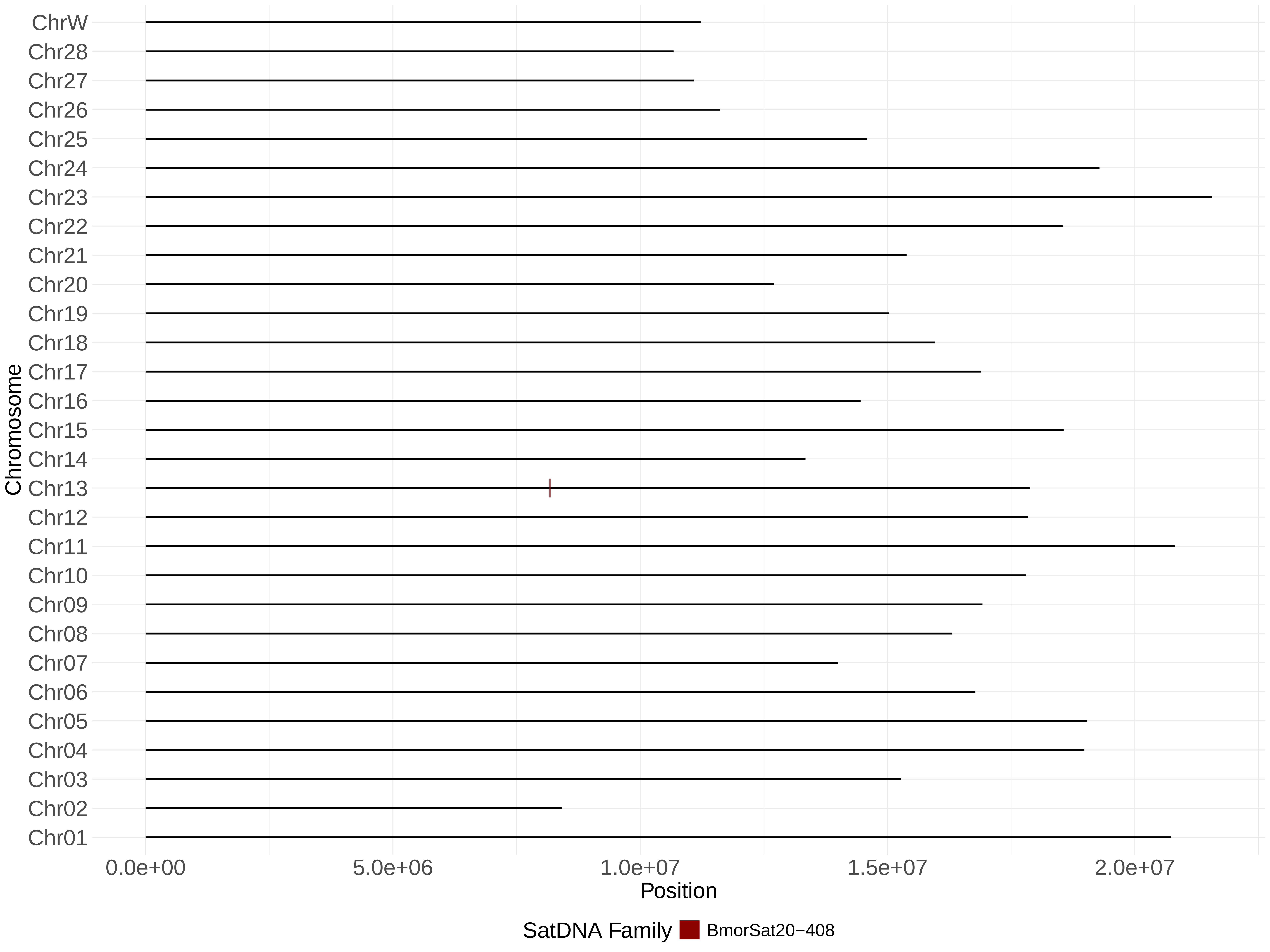

ChrW
Chr28
Chr27
Chr26
Chr25
Chr24
Chr23
Chr22
Chr21
Chr20
Chr19
Chr18
Chr17
Chr16
Chr15
Chr14
Chr13
Chr12
Chr11
Chr10
Chr09
Chr08
Chr07
Chr06
Chr05
Chr04
Chr03
Chr02
Chr01
Chromosome
0.0e+00
5.0e+06
1.0e+07
1.5e+07
2.0e+07
Position
SatDNA Family
BmorSat20−408

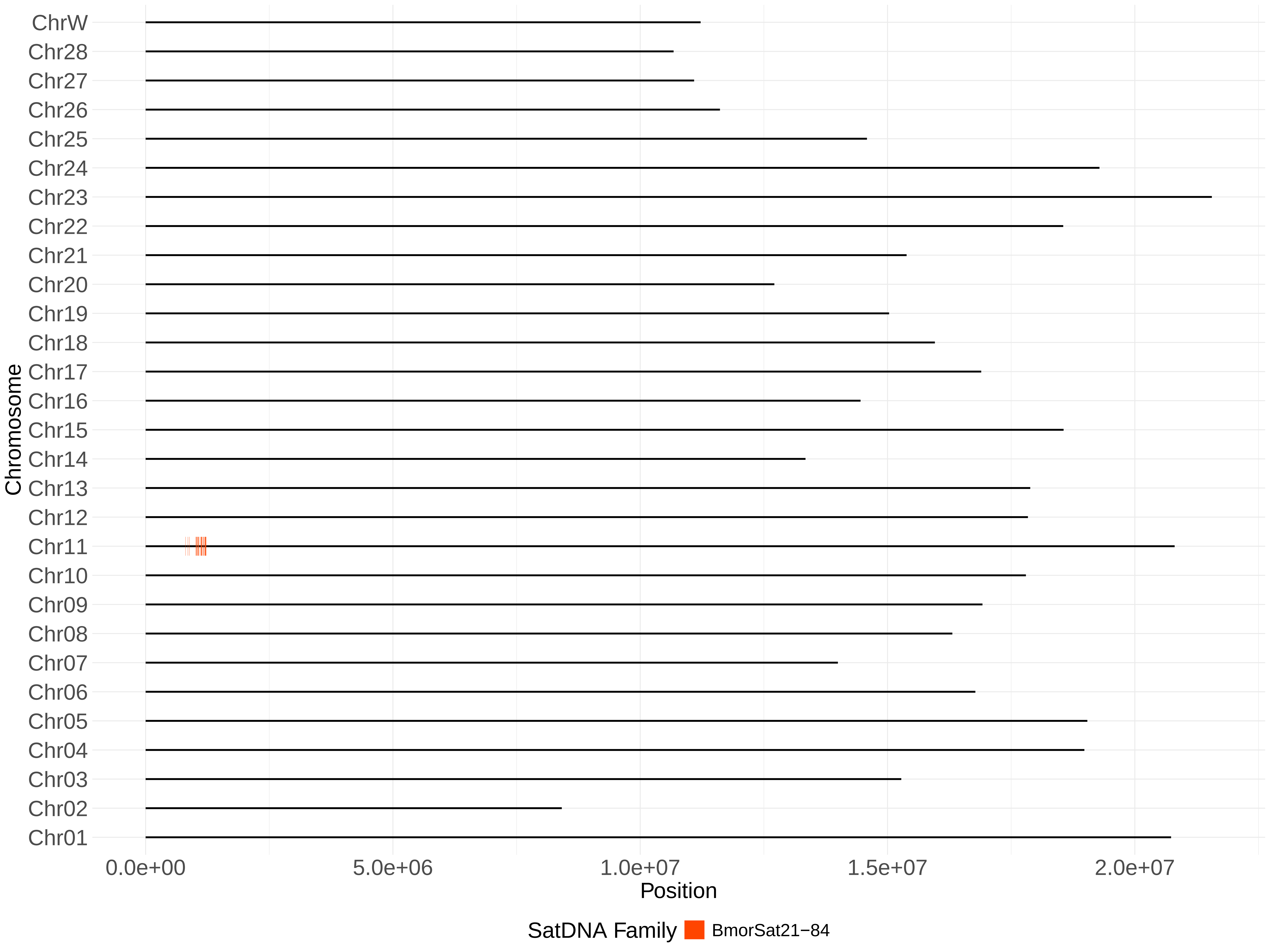

ChrW
Chr28
Chr27
Chr26
Chr25
Chr24
Chr23
Chr22
Chr21
Chr20
Chr19
Chr18
Chr17
Chr16
Chr15
Chr14
Chr13
Chr12
Chr11
Chr10
Chr09
Chr08
Chr07
Chr06
Chr05
Chr04
Chr03
Chr02
Chr01
Chromosome
0.0e+00
5.0e+06
1.0e+07
1.5e+07
2.0e+07
Position
SatDNA Family
BmorSat21−84

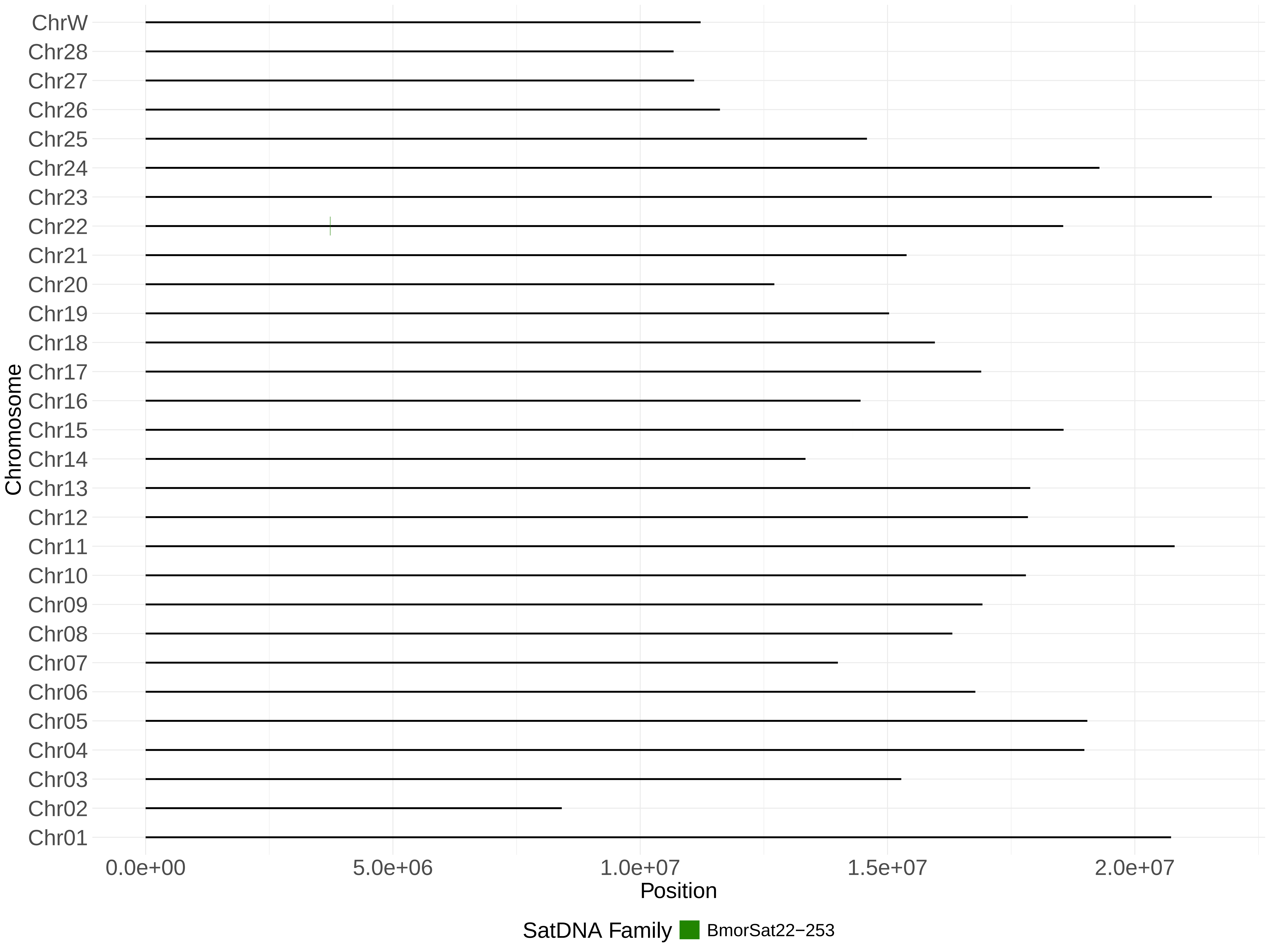

ChrW
Chr28
Chr27
Chr26
Chr25
Chr24
Chr23
Chr22
Chr21
Chr20
Chr19
Chr18
Chr17
Chr16
Chr15
Chr14
Chr13
Chr12
Chr11
Chr10
Chr09
Chr08
Chr07
Chr06
Chr05
Chr04
Chr03
Chr02
Chr01
Chromosome
0.0e+00
5.0e+06
1.0e+07
1.5e+07
2.0e+07
Position
SatDNA Family
BmorSat22−253

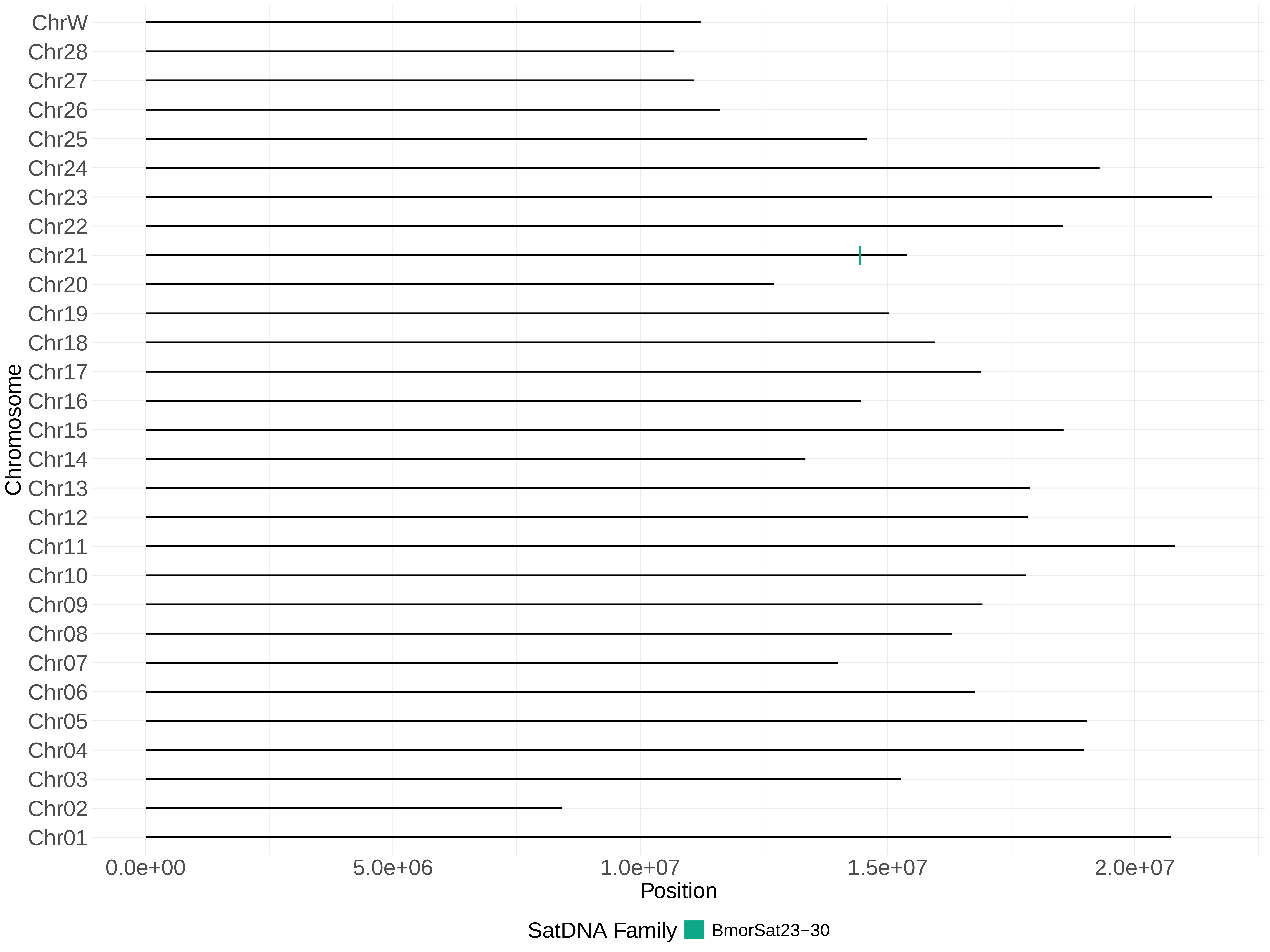

ChrW
Chr28
Chr27
Chr26
Chr25
Chr24
Chr23
Chr22
Chr21
Chr20
Chr19
Chr18
Chr17
Chr16
Chr15
Chr14
Chr13
Chr12
Chr11
Chr10
Chr09
Chr08
Chr07
Chr06
Chr05
Chr04
Chr03
Chr02
Chr01
Chromosome
0.0e+00
5.0e+06
1.0e+07
1.5e+07
2.0e+07
Position
SatDNA Family
BmorSat23−30

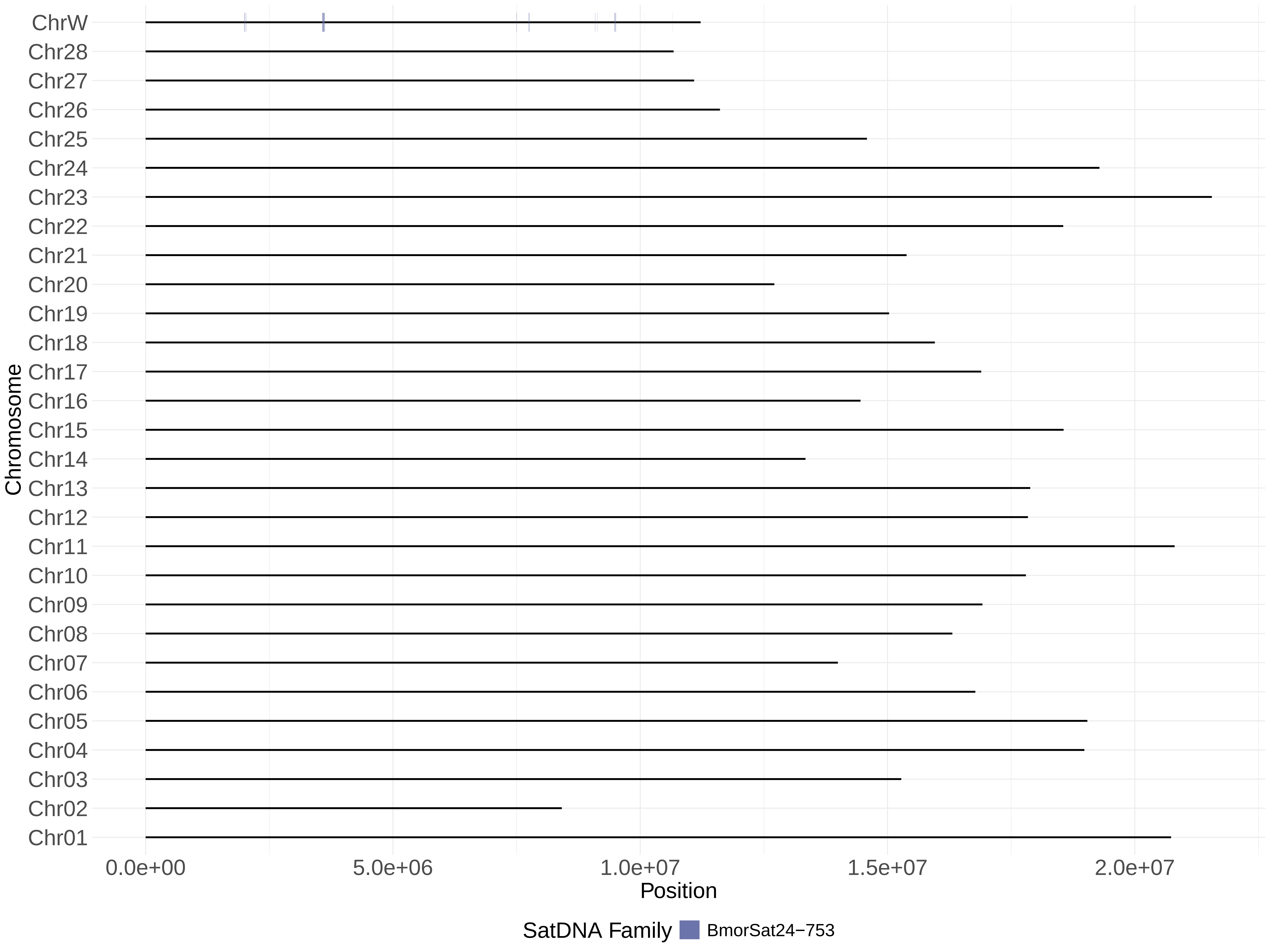
ChrW
Chr28
Chr27
Chr26
Chr25
Chr24
Chr23
Chr22
Chr21
Chr20
Chr19
Chr18
Chr17
Chr16
Chr15
Chr14
Chr13
Chr12
Chr11
Chr10
Chr09
Chr08
Chr07
Chr06
Chr05
Chr04
Chr03
Chr02
Chr01
Chromosome
0.0e+00
5.0e+06
1.0e+07
1.5e+07
2.0e+07
Position
SatDNA Family
BmorSat24−753

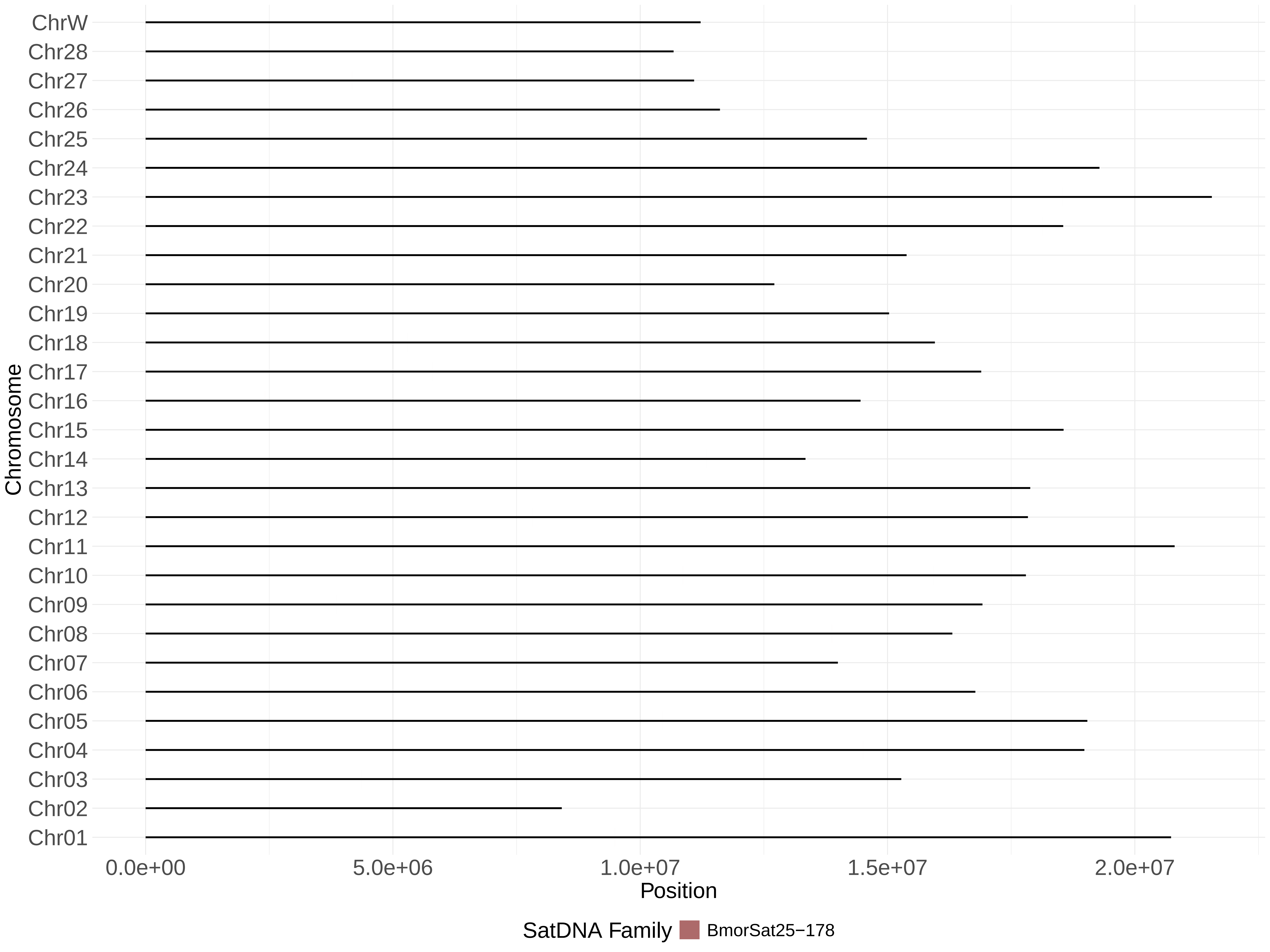
ChrW
Chr28
Chr27
Chr26
Chr25
Chr24
Chr23
Chr22
Chr21
Chr20
Chr19
Chr18
Chr17
Chr16
Chr15
Chr14
Chr13
Chr12
Chr11
Chr10
Chr09
Chr08
Chr07
Chr06
Chr05
Chr04
Chr03
Chr02
Chr01
Chromosome
0.0e+00
5.0e+06
1.0e+07
1.5e+07
2.0e+07
Position
SatDNA Family
BmorSat25−178

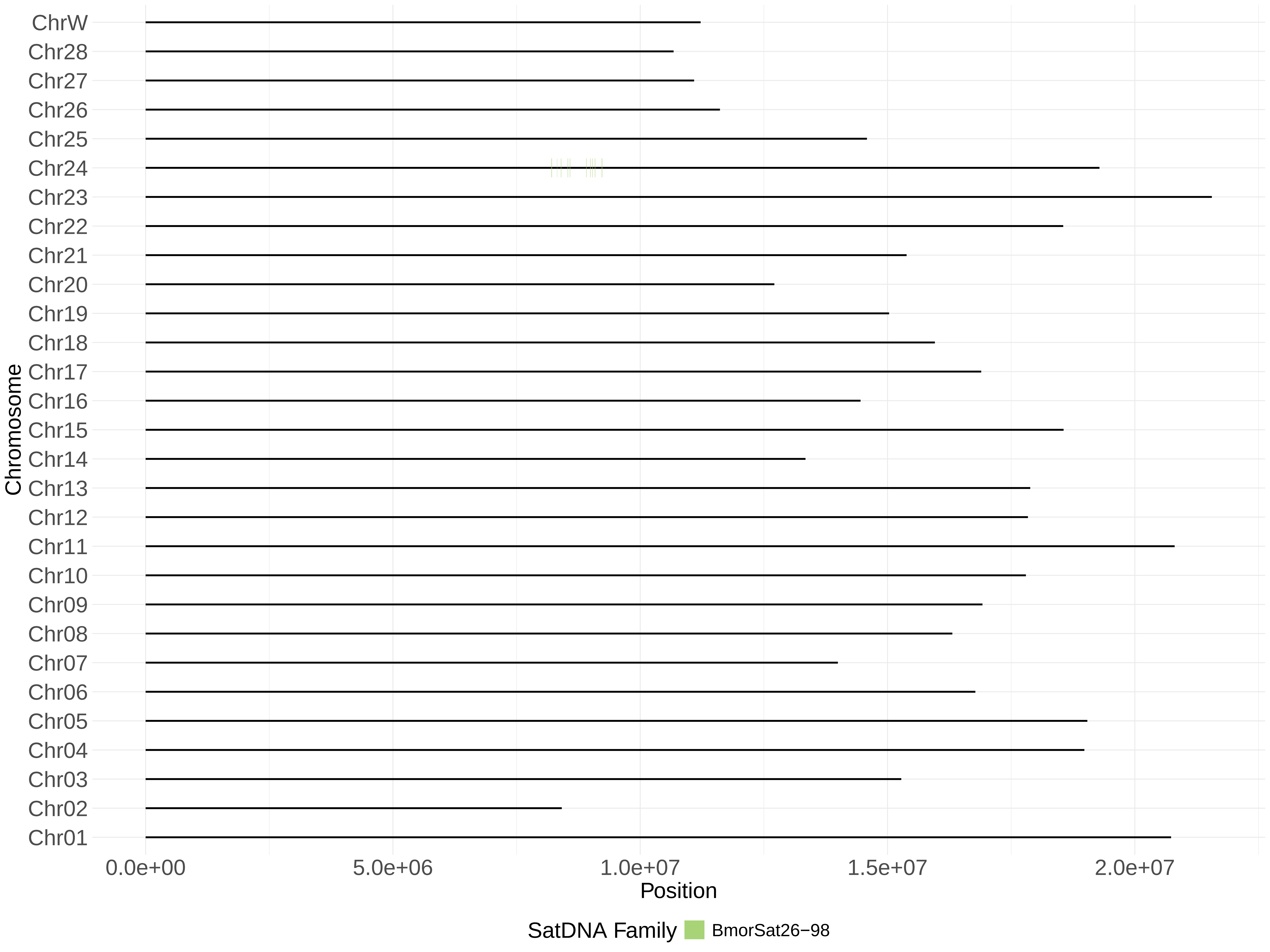
ChrW
Chr28
Chr27
Chr26
Chr25
Chr24
Chr23
Chr22
Chr21
Chr20
Chr19
Chr18
Chr17
Chr16
Chr15
Chr14
Chr13
Chr12
Chr11
Chr10
Chr09
Chr08
Chr07
Chr06
Chr05
Chr04
Chr03
Chr02
Chr01
Chromosome
0.0e+00
5.0e+06
1.0e+07
1.5e+07
2.0e+07
Position
SatDNA Family
BmorSat26−98

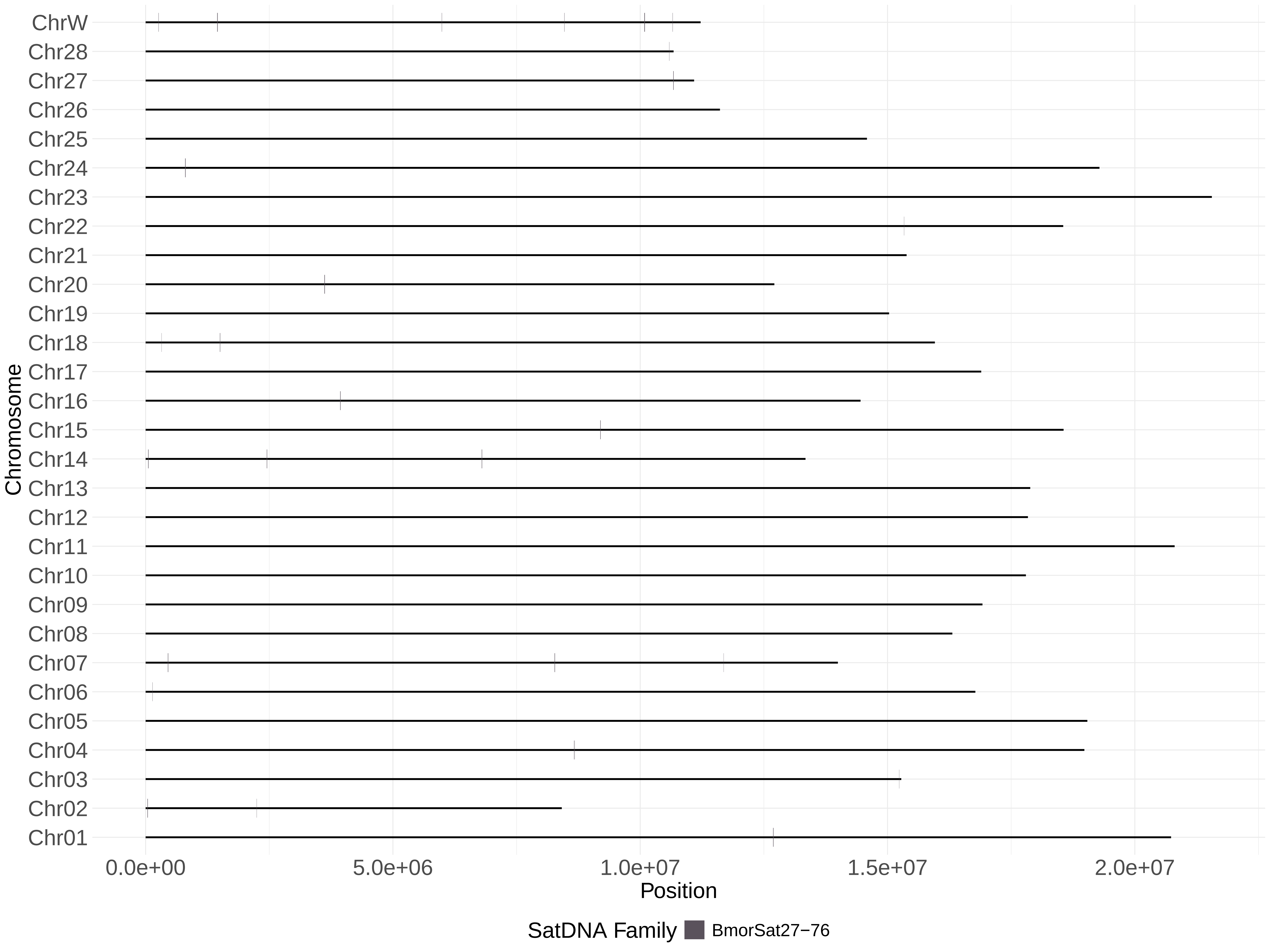

ChrW
Chr28
Chr27
Chr26
Chr25
Chr24
Chr23
Chr22
Chr21
Chr20
Chr19
Chr18
Chr17
Chr16
Chr15
Chr14
Chr13
Chr12
Chr11
Chr10
Chr09
Chr08
Chr07
Chr06
Chr05
Chr04
Chr03
Chr02
Chr01
Chromosome
0.0e+00
5.0e+06
1.0e+07
1.5e+07
2.0e+07
Position
SatDNA Family
BmorSat27−76

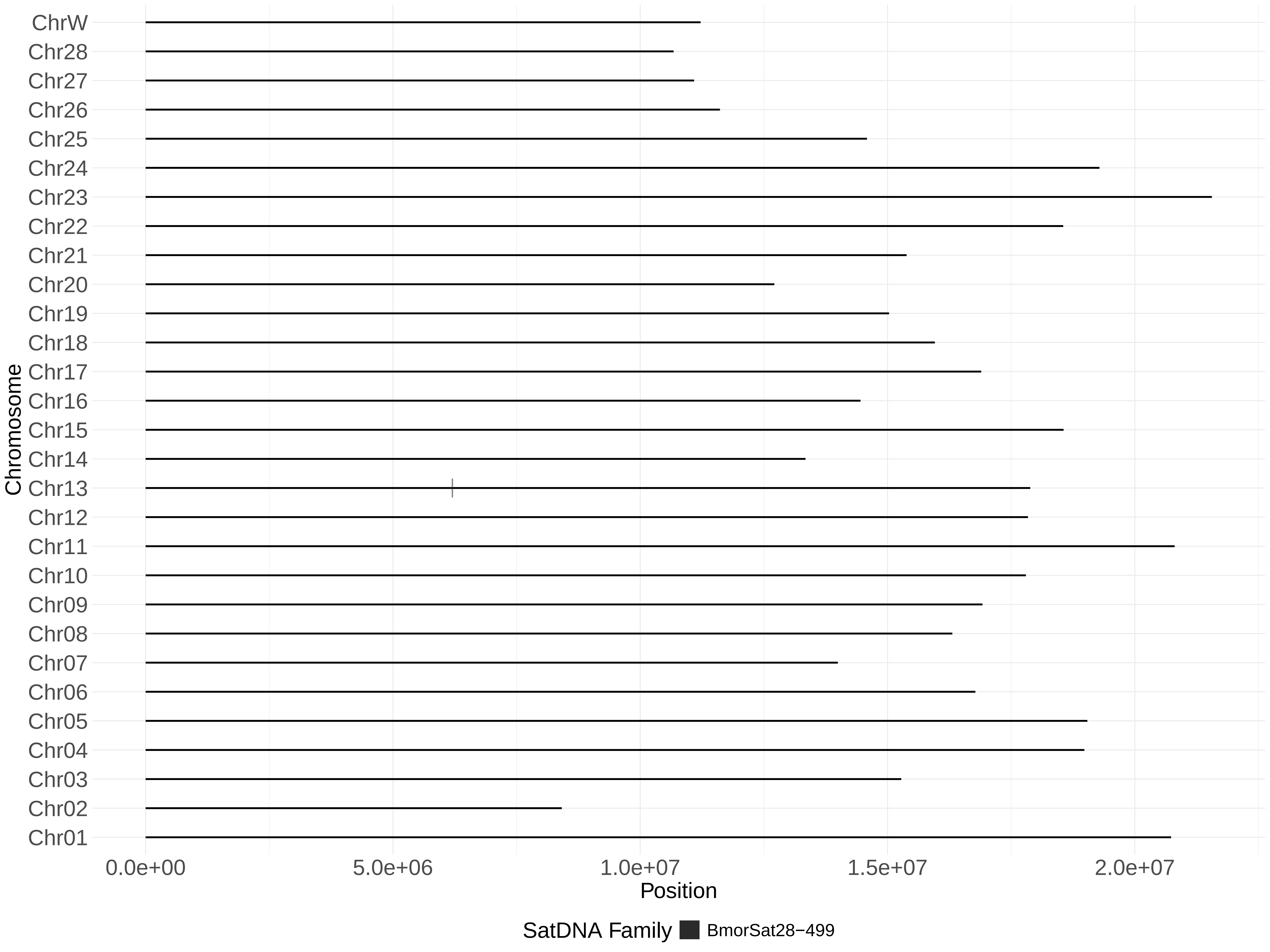
ChrW
Chr28
Chr27
Chr26
Chr25
Chr24
Chr23
Chr22
Chr21
Chr20
Chr19
Chr18
Chr17
Chr16
Chr15
Chr14
Chr13
Chr12
Chr11
Chr10
Chr09
Chr08
Chr07
Chr06
Chr05
Chr04
Chr03
Chr02
Chr01
Chromosome
0.0e+00
5.0e+06
1.0e+07
1.5e+07
2.0e+07
Position
SatDNA Family
BmorSat28−499

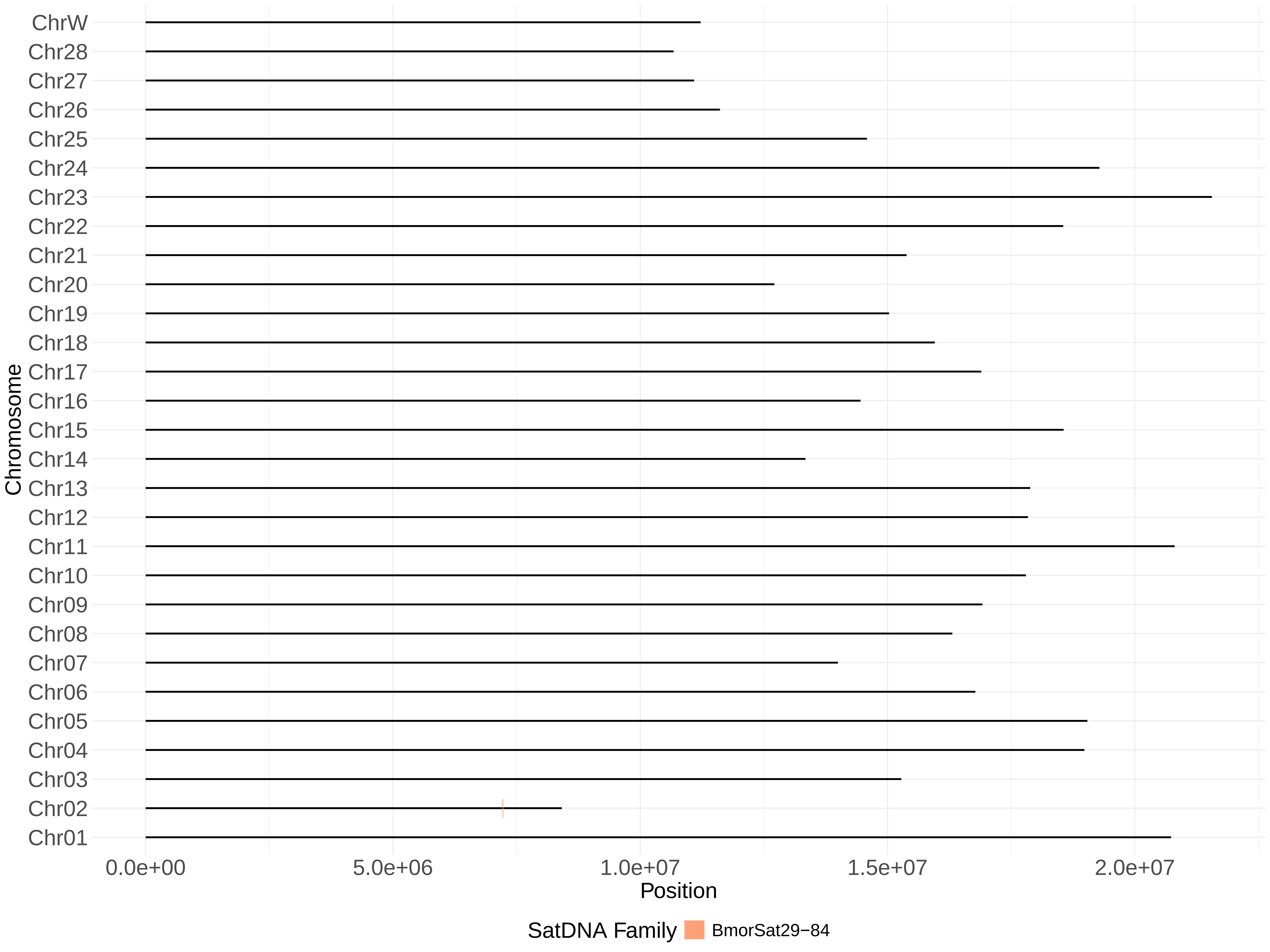
ChrW
Chr28
Chr27
Chr26
Chr25
Chr24
Chr23
Chr22
Chr21
Chr20
Chr19
Chr18
Chr17
Chr16
Chr15
Chr14
Chr13
Chr12
Chr11
Chr10
Chr09
Chr08
Chr07
Chr06
Chr05
Chr04
Chr03
Chr02
Chr01
Chromosome
0.0e+00
5.0e+06
1.0e+07
1.5e+07
2.0e+07
Position
SatDNA Family
BmorSat29−84
