## Supplementary material for "Advancing the understanding of the *Bombyx mori* genome through satellite DNA analysis: low strain differentiation and relationship with transposable elements": Supp fig 5

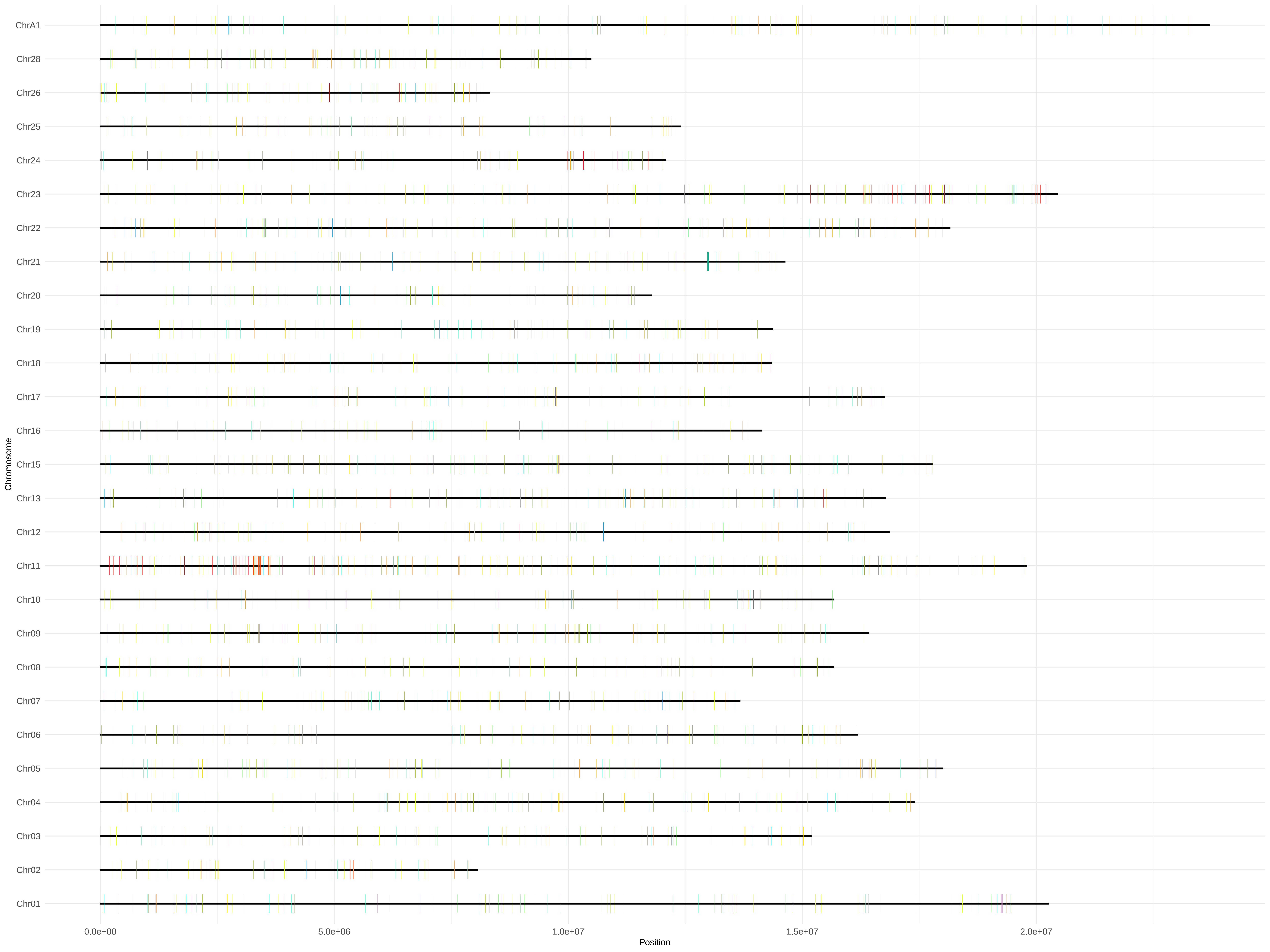
ChrA1
Chr28
Chr26
Chr25
Chr24
Chr23
Chr22
Chr21
Chr20
Chr19
Chr18
Chr17
Chr16
Chr15
Chr13
Chr12
Chr11
Chr10
Chr09
Chr08
Chr07
Chr06
Chr05
Chr04
Chr03
Chr02
Chr01
Chromosome
0.0e+00
5.0e+06
1.0e+07
1.5e+07
2.0e+07
Position

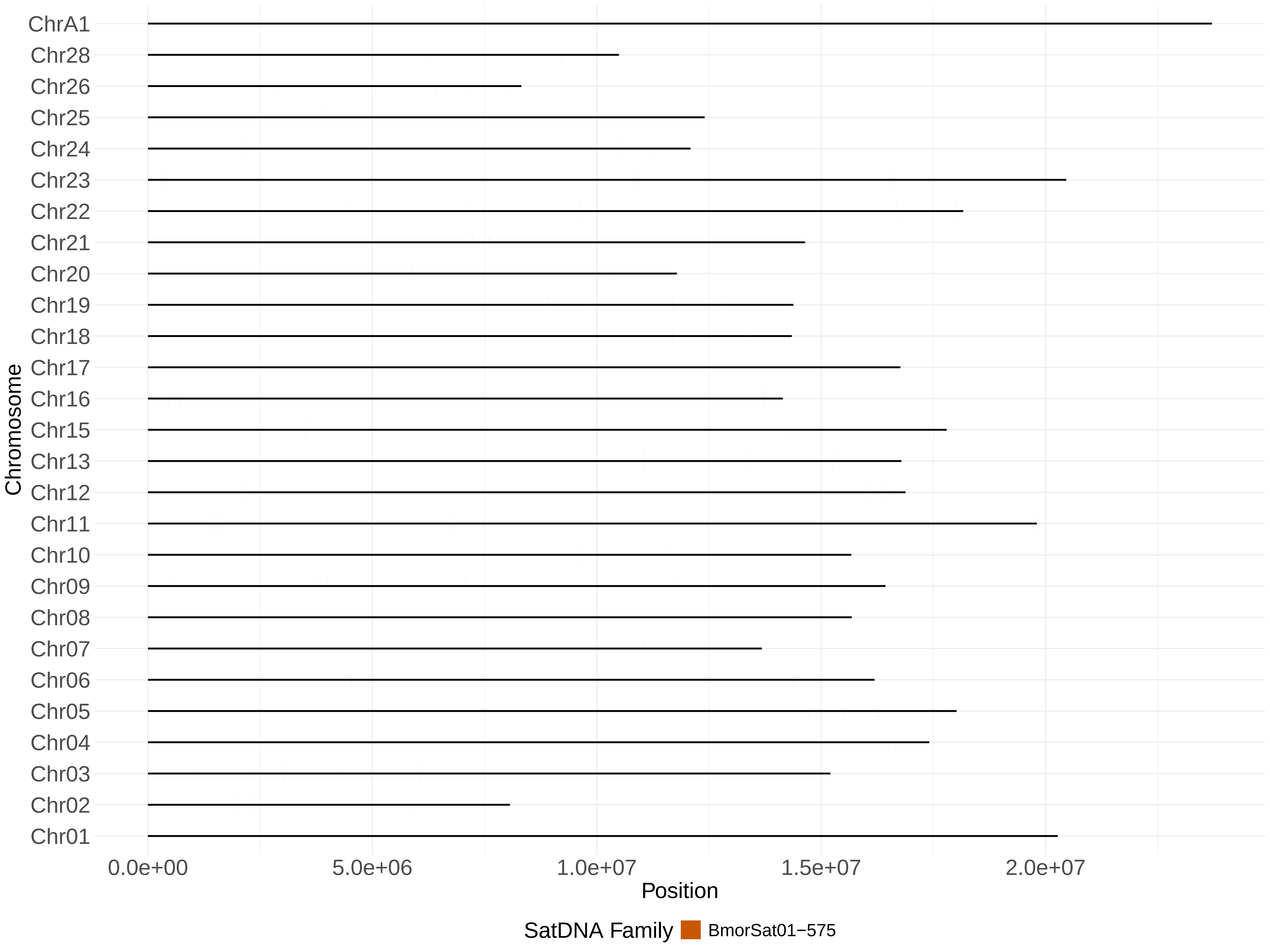
ChrA1
Chr28
Chr26
Chr25
Chr24
Chr23
Chr22
Chr21
Chr20
Chr19
Chr18
Chr17
Chr16
Chr15
Chr13
Chr12
Chr11
Chr10
Chr09
Chr08
Chr07
Chr06
Chr05
Chr04
Chr03
Chr02
Chr01
Chromosome
0.0e+00
5.0e+06
1.0e+07
1.5e+07
2.0e+07
Position
SatDNA Family
BmorSat01−575

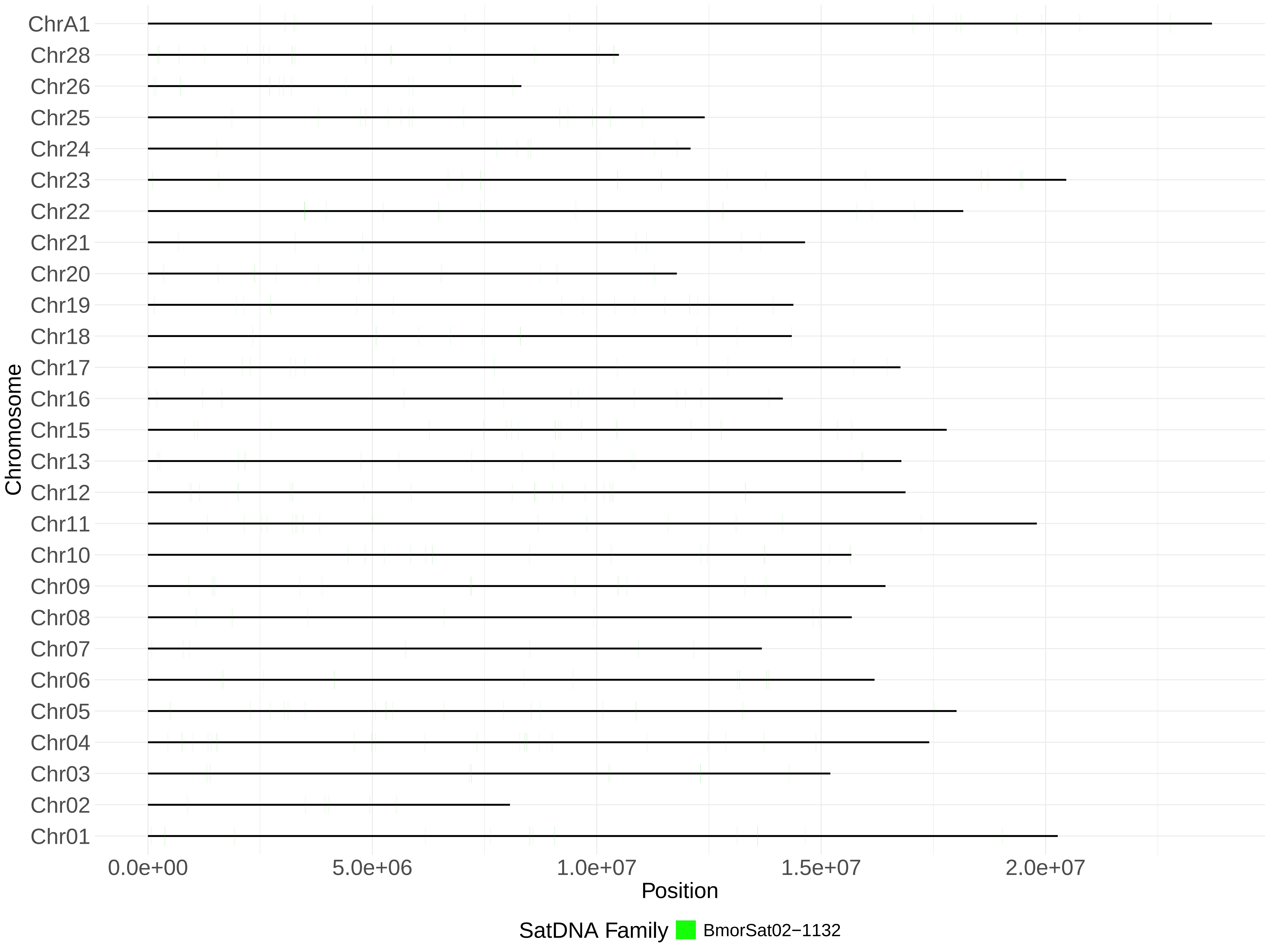

ChrA1
Chr28
Chr26
Chr25
Chr24
Chr23
Chr22
Chr21
Chr20
Chr19
Chr18
Chr17
Chr16
Chr15
Chr13
Chr12
Chr11
Chr10
Chr09
Chr08
Chr07
Chr06
Chr05
Chr04
Chr03
Chr02
Chr01
Chromosome
0.0e+00
5.0e+06
1.0e+07
1.5e+07
2.0e+07
Position
SatDNA Family
BmorSat02−1132

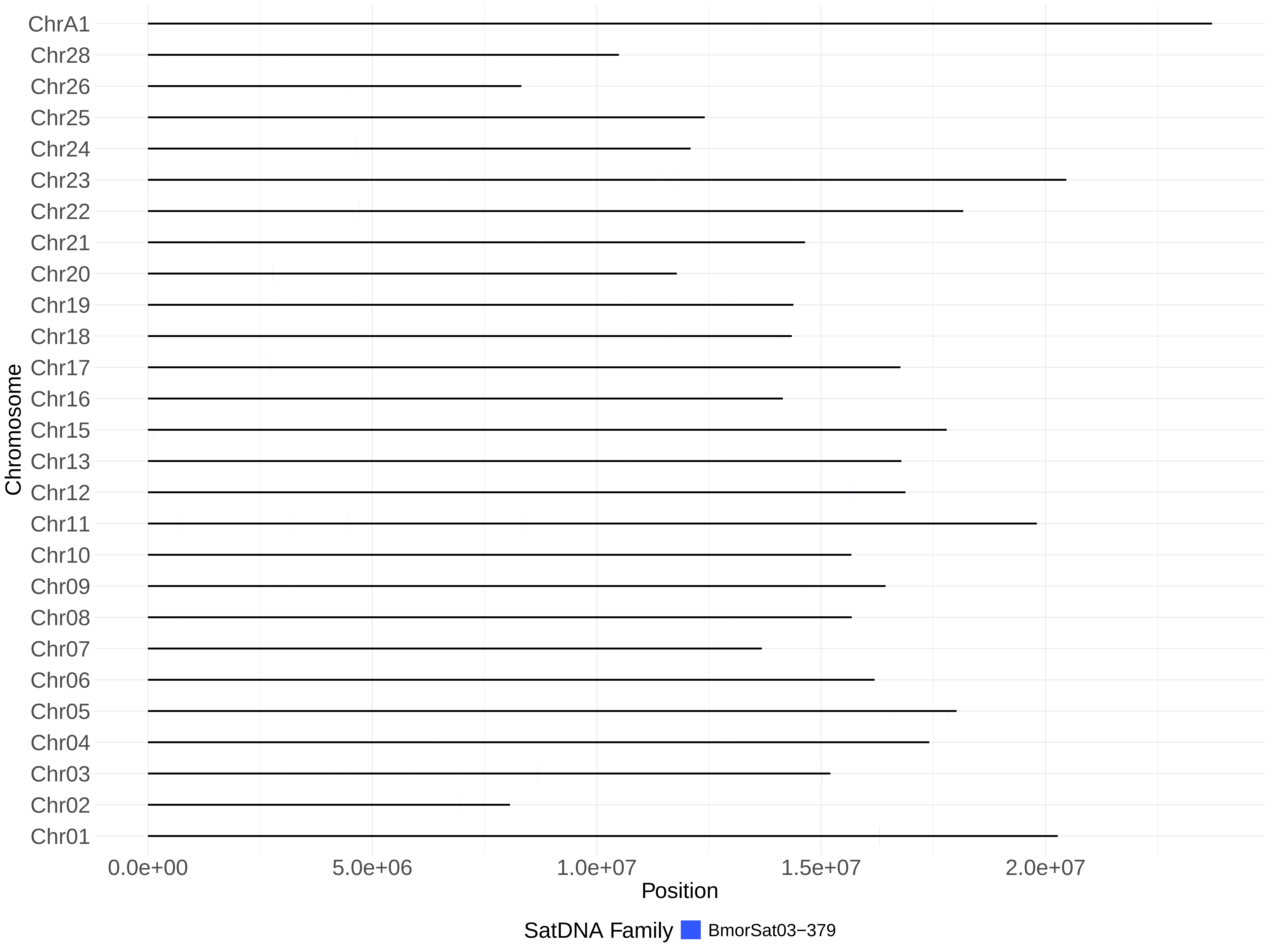
ChrA1
Chr28
Chr26
Chr25
Chr24
Chr23
Chr22
Chr21
Chr20
Chr19
Chr18
Chr17
Chr16
Chr15
Chr13
Chr12
Chr11
Chr10
Chr09
Chr08
Chr07
Chr06
Chr05
Chr04
Chr03
Chr02
Chr01
Chromosome
0.0e+00
5.0e+06
1.0e+07
1.5e+07
2.0e+07
Position
SatDNA Family
BmorSat03−379

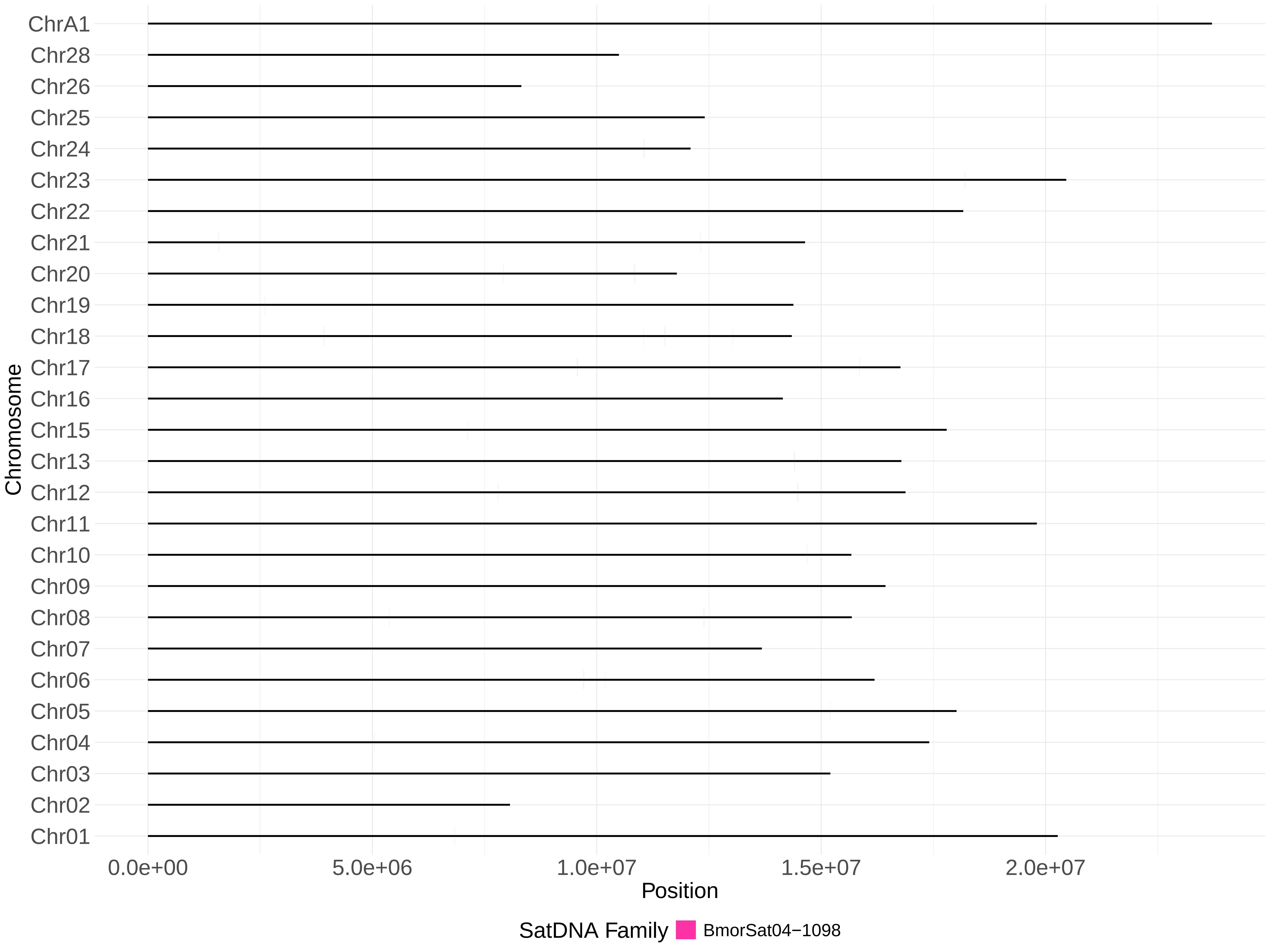

ChrA1
Chr28
Chr26
Chr25
Chr24
Chr23
Chr22
Chr21
Chr20
Chr19
Chr18
Chr17
Chr16
Chr15
Chr13
Chr12
Chr11
Chr10
Chr09
Chr08
Chr07
Chr06
Chr05
Chr04
Chr03
Chr02
Chr01
Chromosome
0.0e+00
5.0e+06
1.0e+07
1.5e+07
2.0e+07
Position
SatDNA Family
BmorSat04−1098

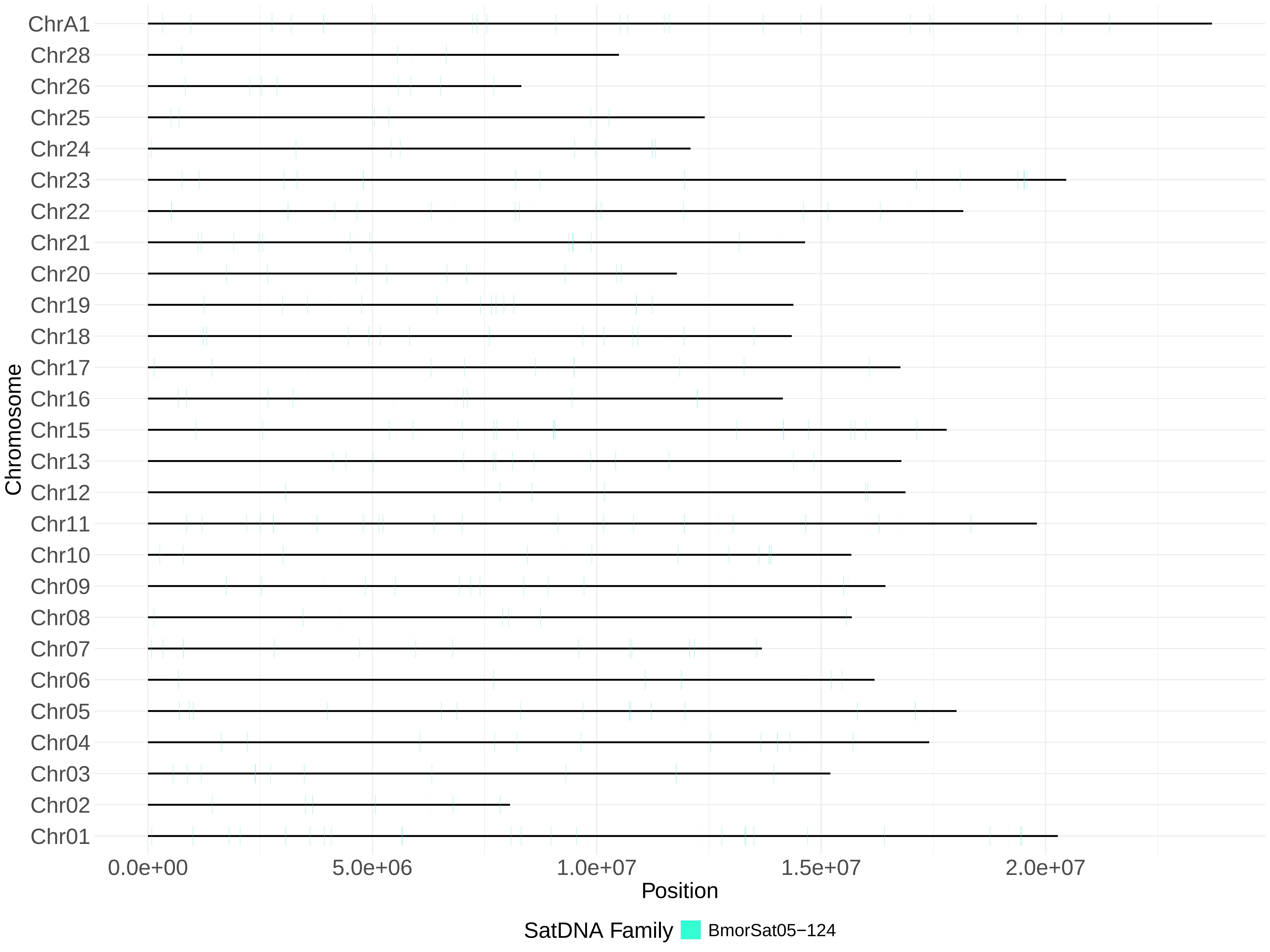
ChrA1
Chr28
Chr26
Chr25
Chr24
Chr23
Chr22
Chr21
Chr20
Chr19
Chr18
Chr17
Chr16
Chr15
Chr13
Chr12
Chr11
Chr10
Chr09
Chr08
Chr07
Chr06
Chr05
Chr04
Chr03
Chr02
Chr01
Chromosome
0.0e+00
5.0e+06
1.0e+07
1.5e+07
2.0e+07
Position
SatDNA Family
BmorSat05−124

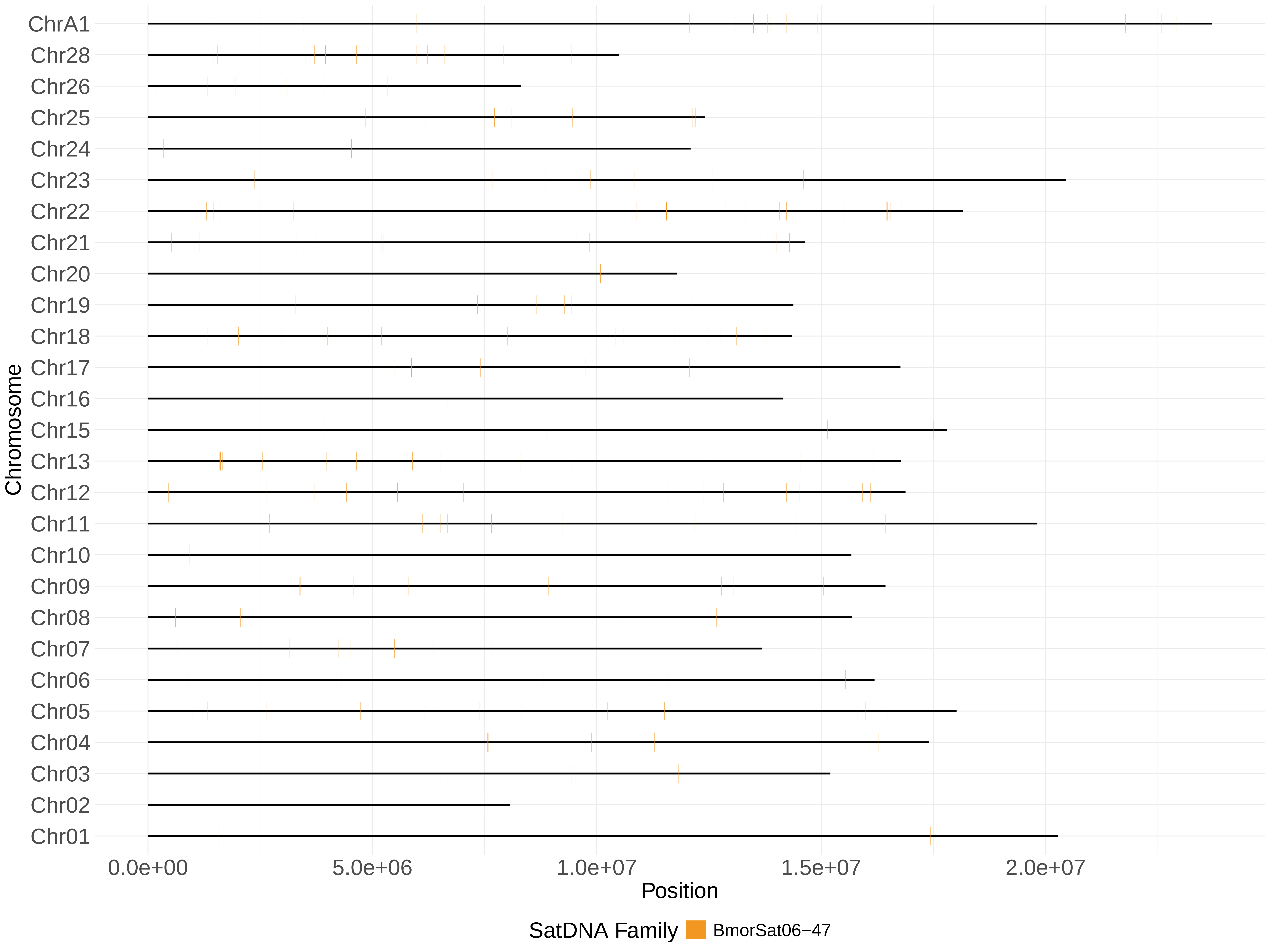

ChrA1
Chr28
Chr26
Chr25
Chr24
Chr23
Chr22
Chr21
Chr20
Chr19
Chr18
Chr17
Chr16
Chr15
Chr13
Chr12
Chr11
Chr10
Chr09
Chr08
Chr07
Chr06
Chr05
Chr04
Chr03
Chr02
Chr01
Chromosome
0.0e+00
5.0e+06
1.0e+07
1.5e+07
2.0e+07
Position
SatDNA Family
BmorSat06−47

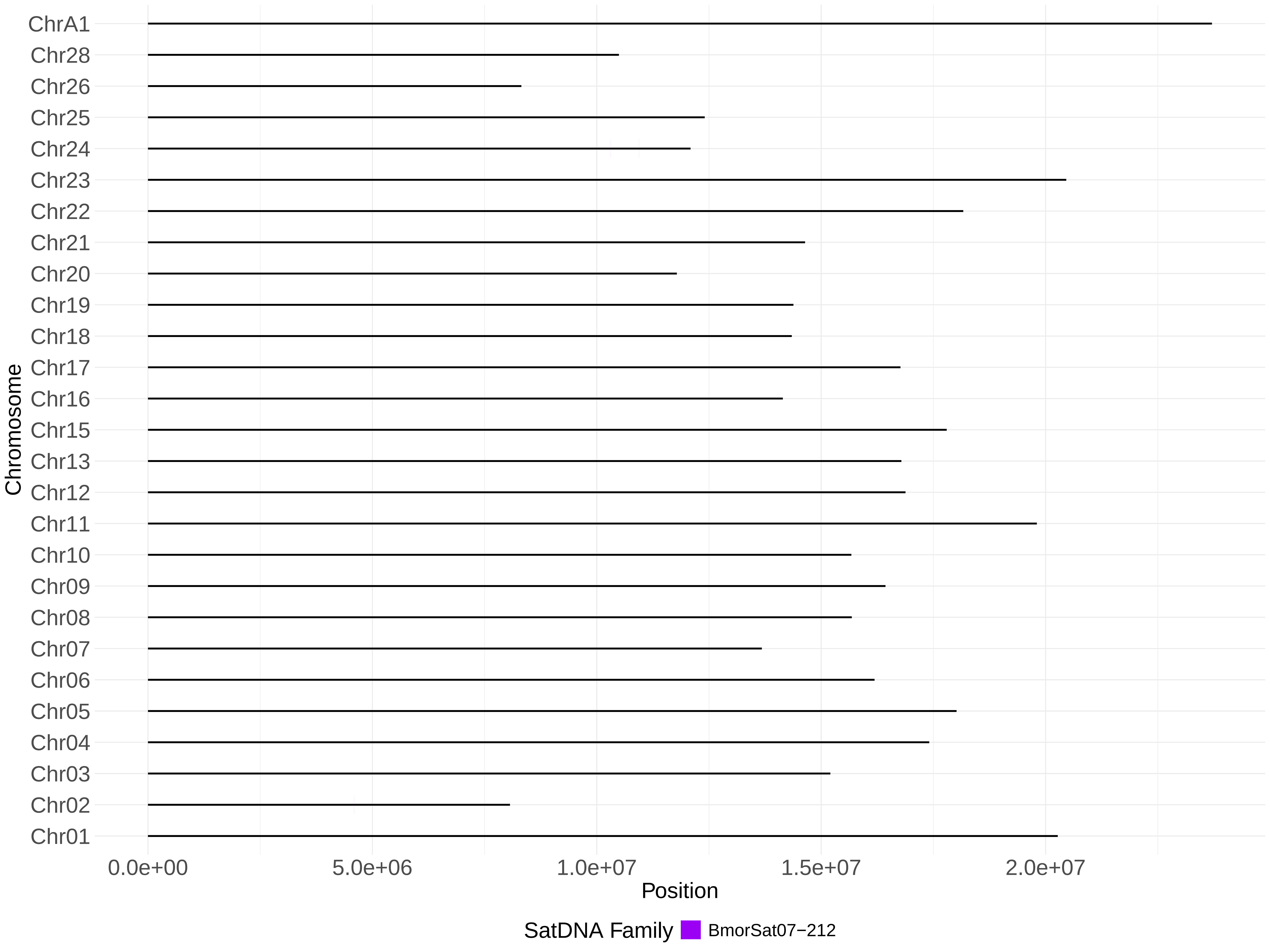

ChrA1
Chr28
Chr26
Chr25
Chr24
Chr23
Chr22
Chr21
Chr20
Chr19
Chr18
Chr17
Chr16
Chr15
Chr13
Chr12
Chr11
Chr10
Chr09
Chr08
Chr07
Chr06
Chr05
Chr04
Chr03
Chr02
Chr01
Chromosome
0.0e+00
5.0e+06
1.0e+07
1.5e+07
2.0e+07
Position
SatDNA Family
BmorSat07−212

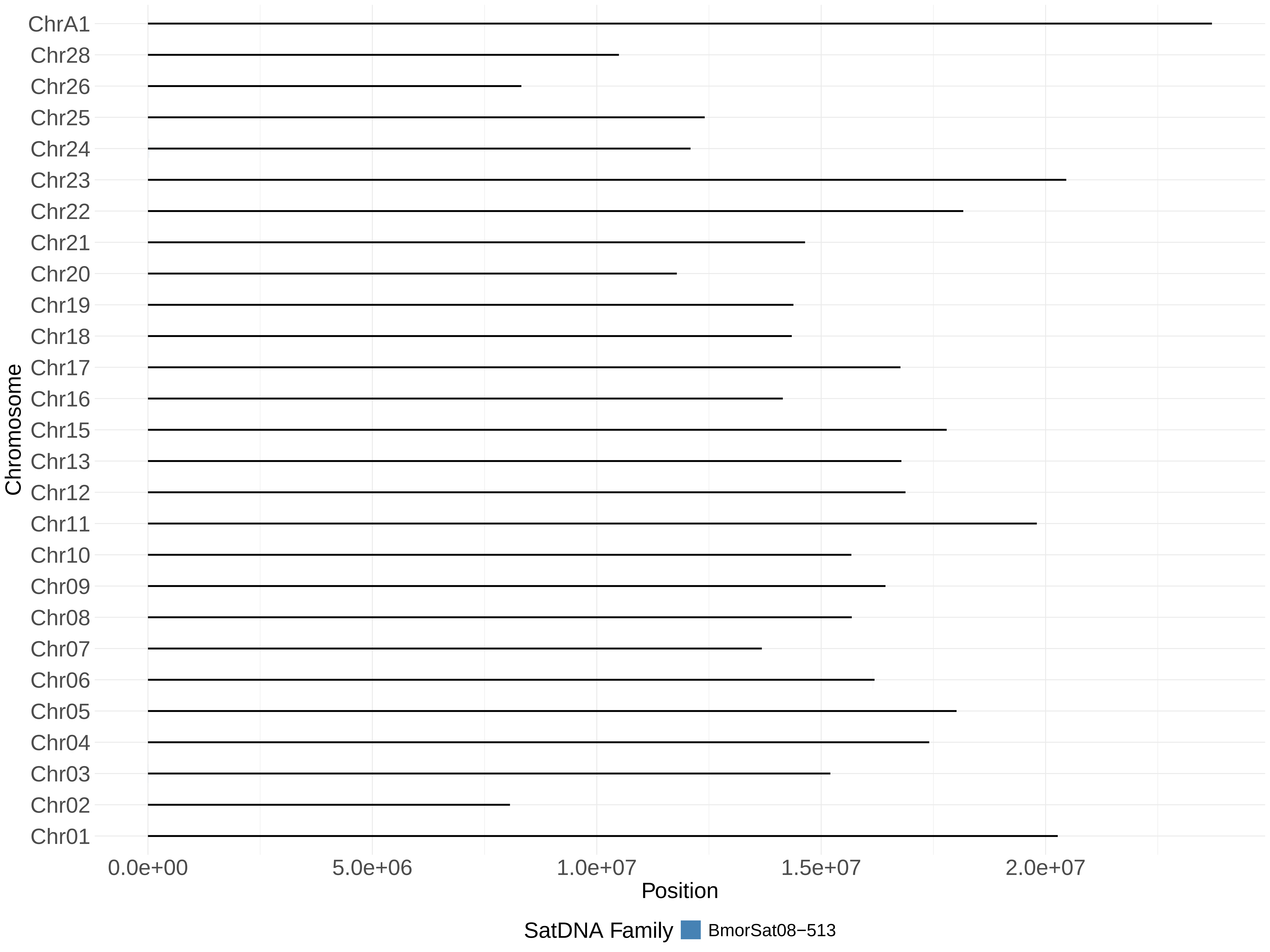
ChrA1
Chr28
Chr26
Chr25
Chr24
Chr23
Chr22
Chr21
Chr20
Chr19
Chr18
Chr17
Chr16
Chr15
Chr13
Chr12
Chr11
Chr10
Chr09
Chr08
Chr07
Chr06
Chr05
Chr04
Chr03
Chr02
Chr01
Chromosome
0.0e+00
5.0e+06
1.0e+07
1.5e+07
2.0e+07
Position
SatDNA Family
BmorSat08−513

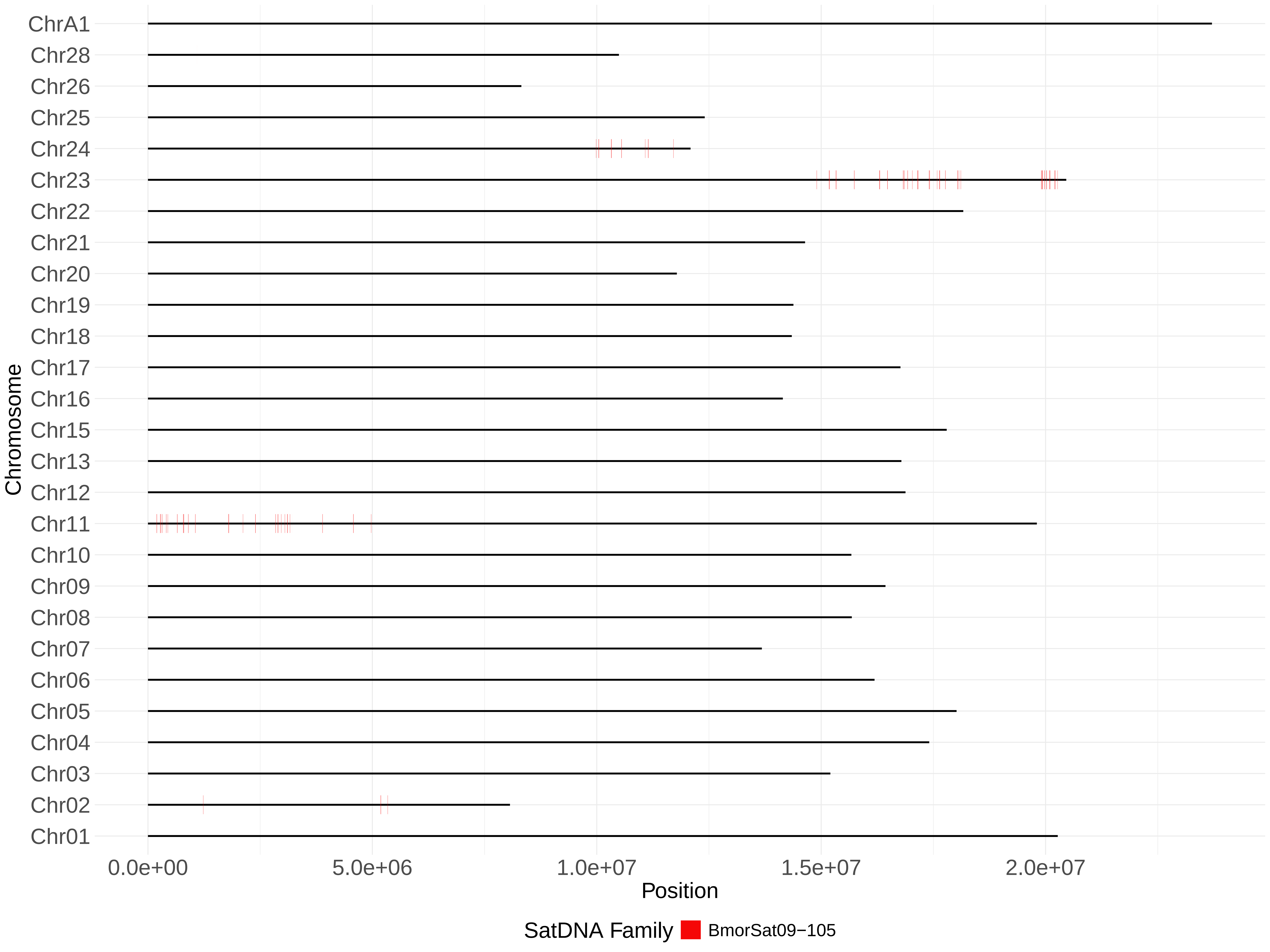

ChrA1
Chr28
Chr26
Chr25
Chr24
Chr23
Chr22
Chr21
Chr20
Chr19
Chr18
Chr17
Chr16
Chr15
Chr13
Chr12
Chr11
Chr10
Chr09
Chr08
Chr07
Chr06
Chr05
Chr04
Chr03
Chr02
Chr01
Chromosome
0.0e+00
5.0e+06
1.0e+07
1.5e+07
2.0e+07
Position
SatDNA Family
BmorSat09−105

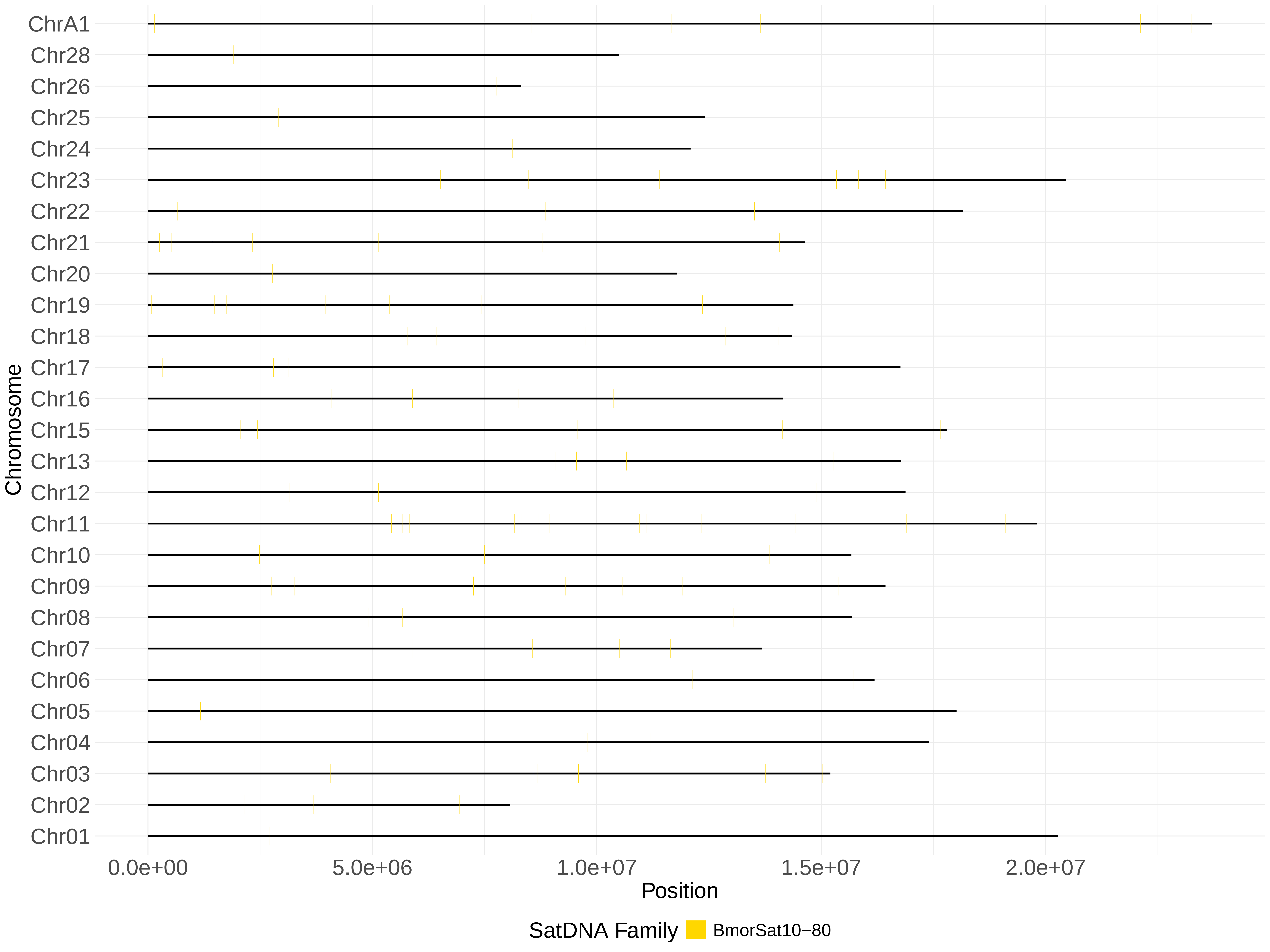
ChrA1
Chr28
Chr26
Chr25
Chr24
Chr23
Chr22
Chr21
Chr20
Chr19
Chr18
Chr17
Chr16
Chr15
Chr13
Chr12
Chr11
Chr10
Chr09
Chr08
Chr07
Chr06
Chr05
Chr04
Chr03
Chr02
Chr01
Chromosome
0.0e+00
5.0e+06
1.0e+07
1.5e+07
2.0e+07
Position
SatDNA Family
BmorSat10−80

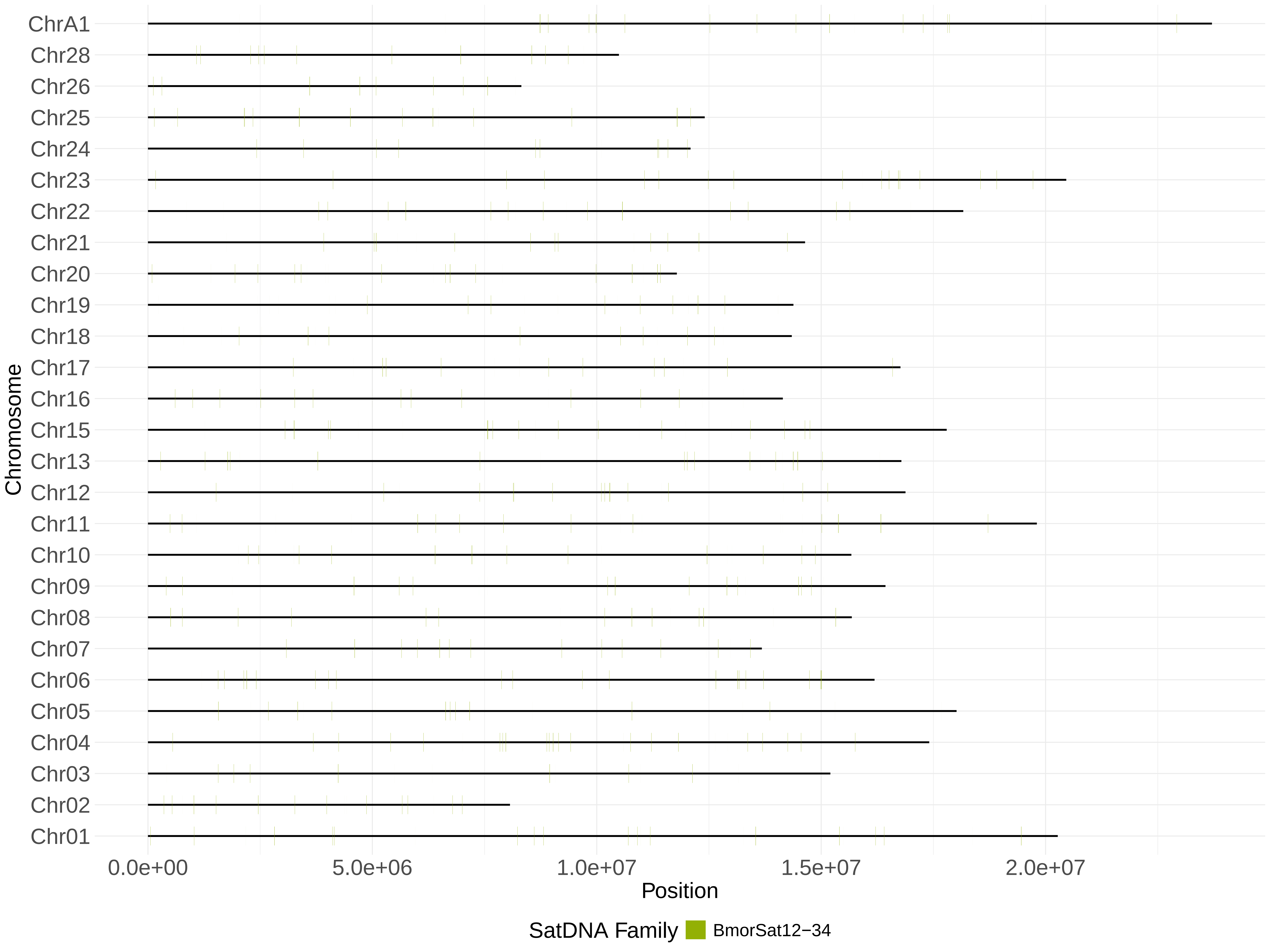
ChrA1
Chr28
Chr26
Chr25
Chr24
Chr23
Chr22
Chr21
Chr20
Chr19
Chr18
Chr17
Chr16
Chr15
Chr13
Chr12
Chr11
Chr10
Chr09
Chr08
Chr07
Chr06
Chr05
Chr04
Chr03
Chr02
Chr01
Chromosome
0.0e+00
5.0e+06
1.0e+07
1.5e+07
2.0e+07
Position
SatDNA Family
BmorSat12−34

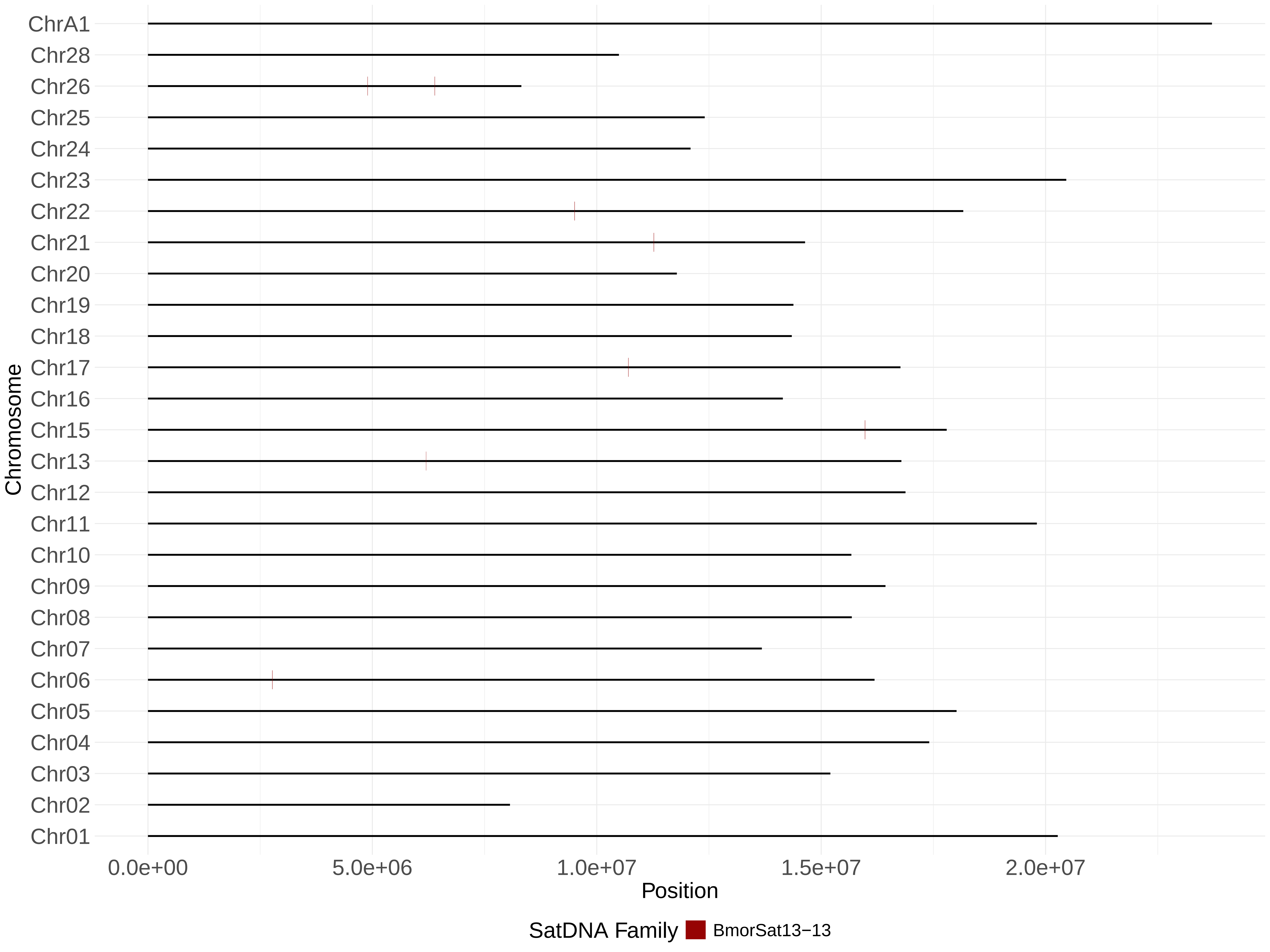
ChrA1
Chr28
Chr26
Chr25
Chr24
Chr23
Chr22
Chr21
Chr20
Chr19
Chr18
Chr17
Chr16
Chr15
Chr13
Chr12
Chr11
Chr10
Chr09
Chr08
Chr07
Chr06
Chr05
Chr04
Chr03
Chr02
Chr01
Chromosome
0.0e+00
5.0e+06
1.0e+07
1.5e+07
2.0e+07
Position
SatDNA Family
BmorSat13−13

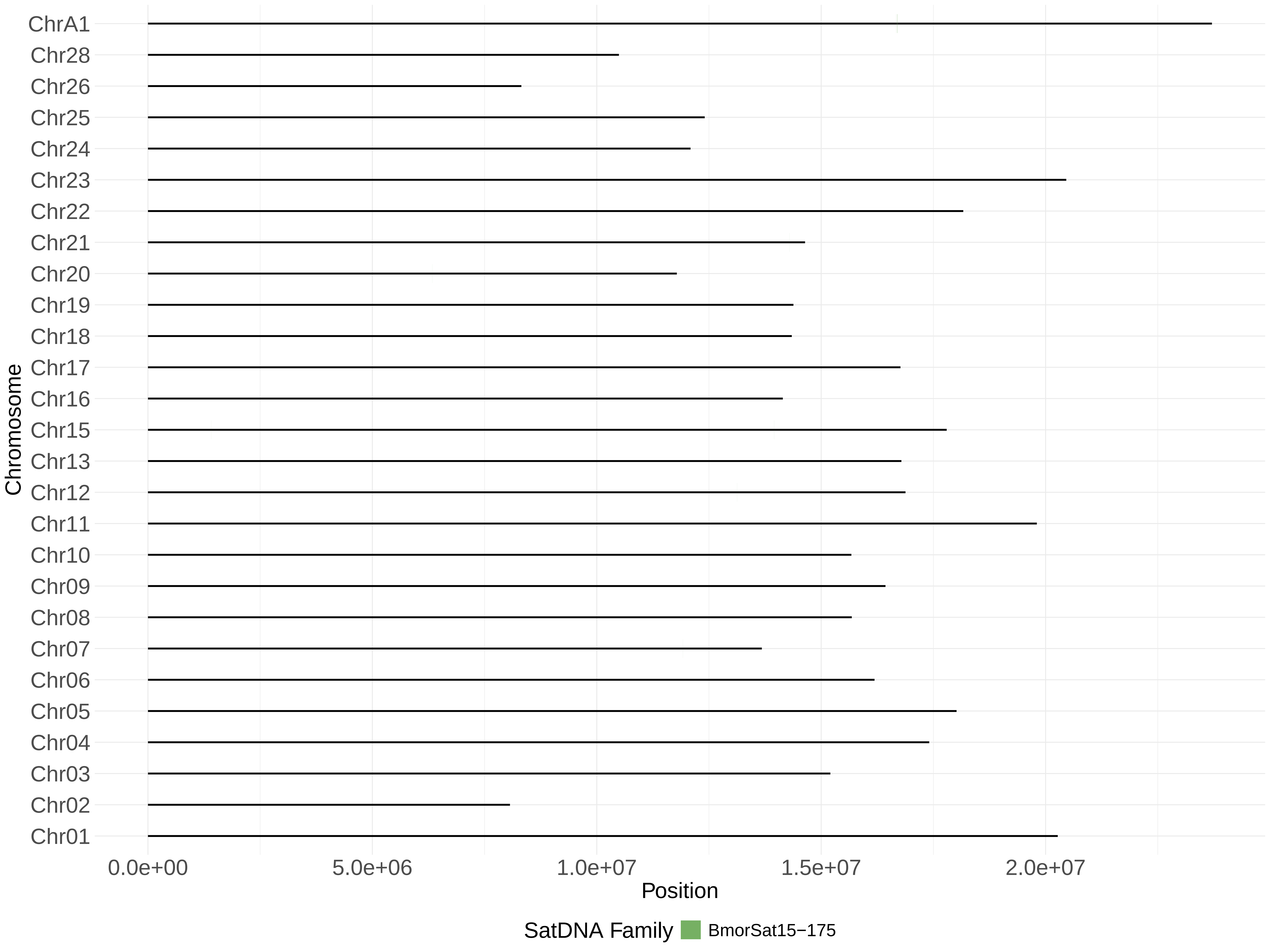

ChrA1
Chr28
Chr26
Chr25
Chr24
Chr23
Chr22
Chr21
Chr20
Chr19
Chr18
Chr17
Chr16
Chr15
Chr13
Chr12
Chr11
Chr10
Chr09
Chr08
Chr07
Chr06
Chr05
Chr04
Chr03
Chr02
Chr01
Chromosome
0.0e+00
5.0e+06
1.0e+07
1.5e+07
2.0e+07
Position
SatDNA Family
BmorSat15−175

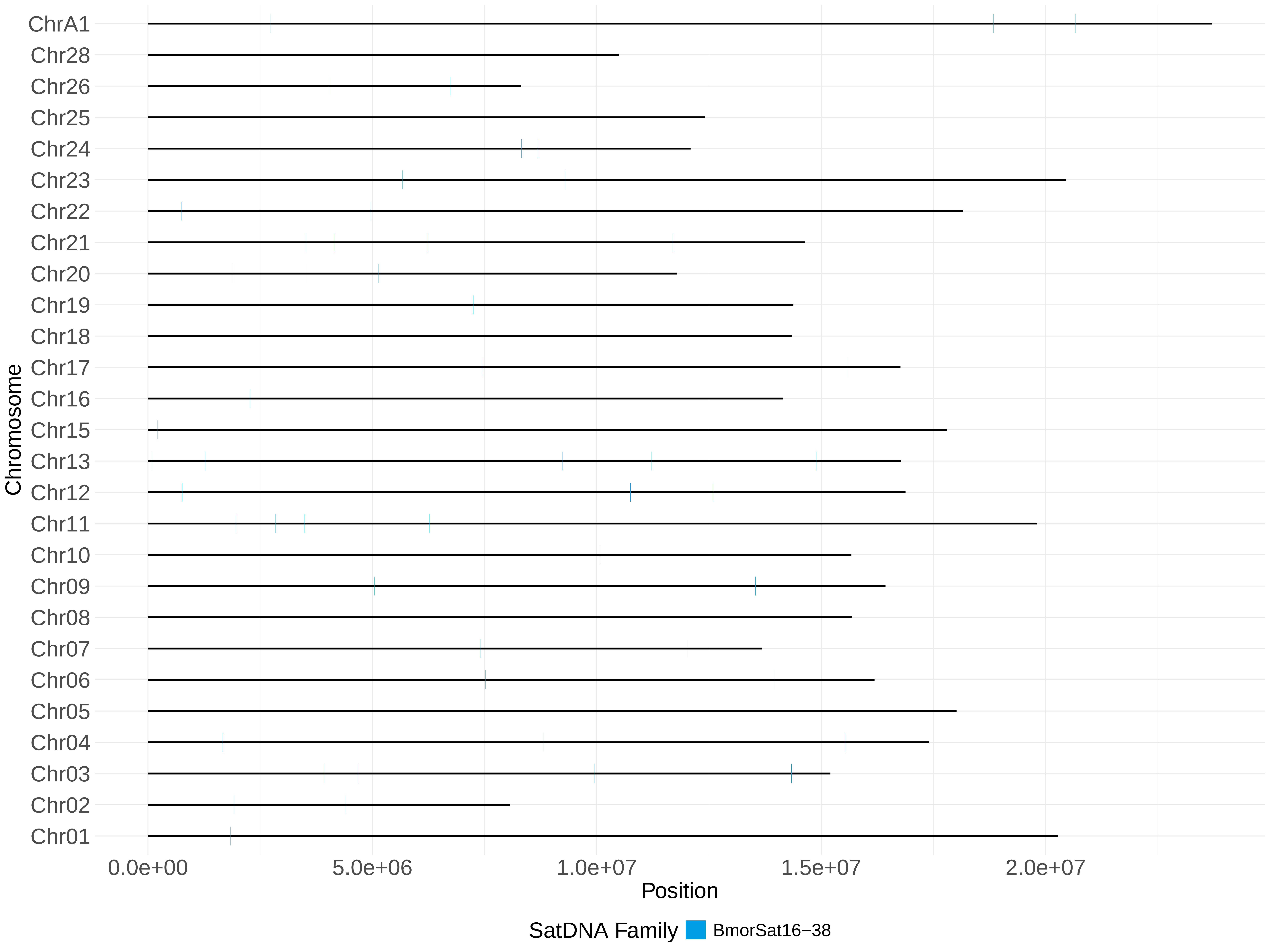
ChrA1
Chr28
Chr26
Chr25
Chr24
Chr23
Chr22
Chr21
Chr20
Chr19
Chr18
Chr17
Chr16
Chr15
Chr13
Chr12
Chr11
Chr10
Chr09
Chr08
Chr07
Chr06
Chr05
Chr04
Chr03
Chr02
Chr01
Chromosome
0.0e+00
5.0e+06
1.0e+07
1.5e+07
2.0e+07
Position
SatDNA Family
BmorSat16−38

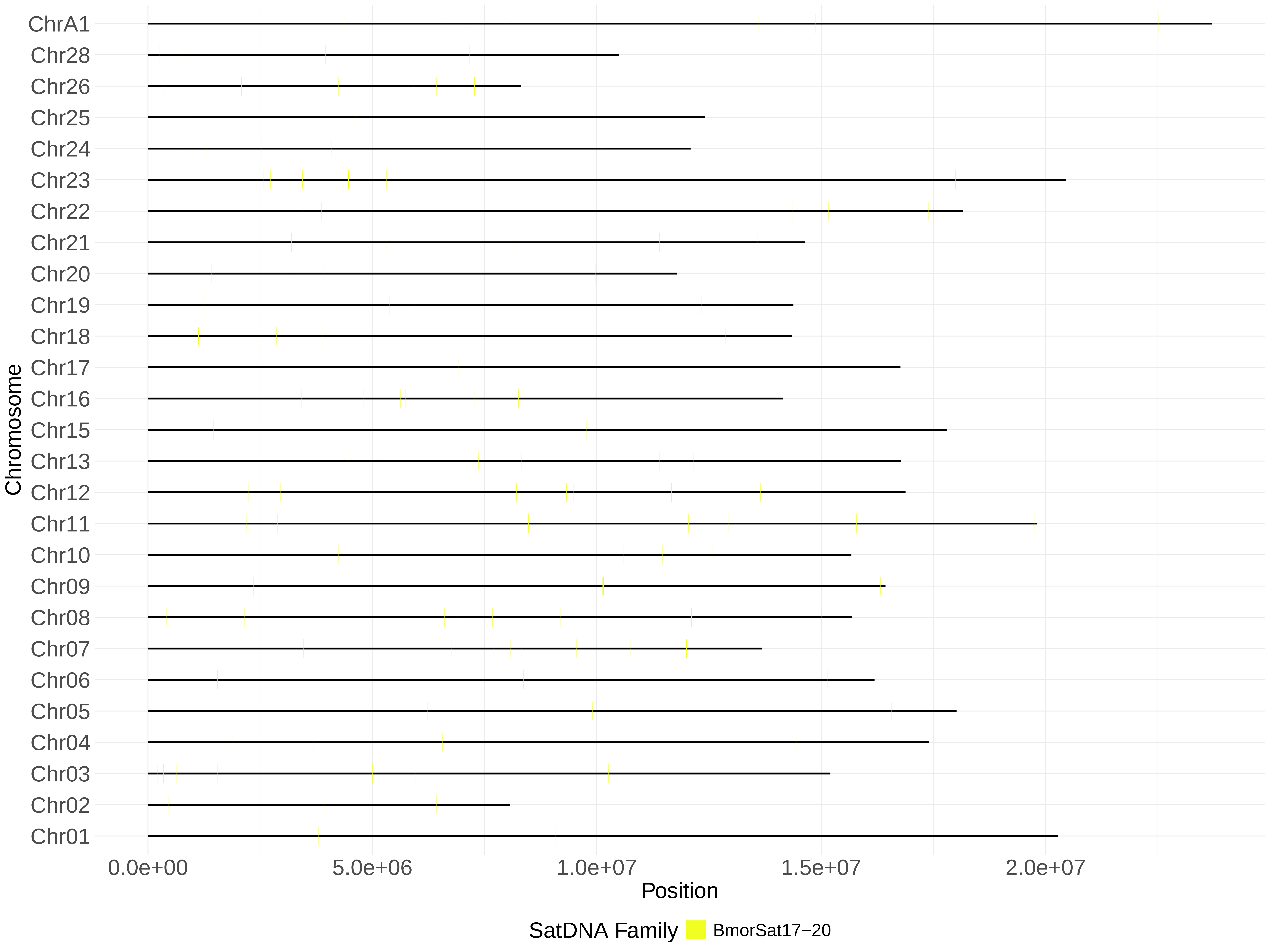
ChrA1
Chr28
Chr26
Chr25
Chr24
Chr23
Chr22
Chr21
Chr20
Chr19
Chr18
Chr17
Chr16
Chr15
Chr13
Chr12
Chr11
Chr10
Chr09
Chr08
Chr07
Chr06
Chr05
Chr04
Chr03
Chr02
Chr01
Chromosome
0.0e+00
5.0e+06
1.0e+07
1.5e+07
2.0e+07
Position
SatDNA Family
BmorSat17−20

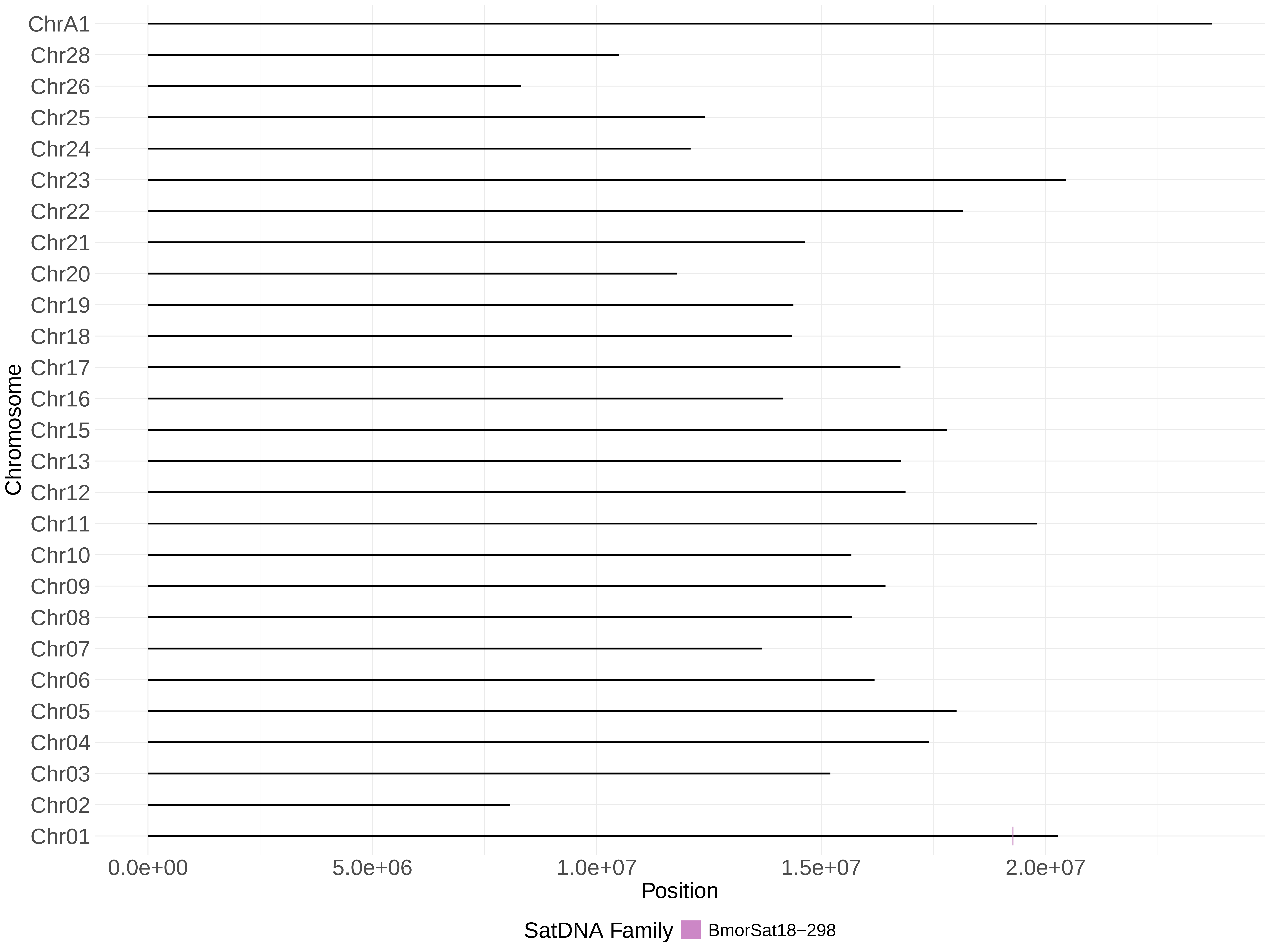

ChrA1
Chr28
Chr26
Chr25
Chr24
Chr23
Chr22
Chr21
Chr20
Chr19
Chr18
Chr17
Chr16
Chr15
Chr13
Chr12
Chr11
Chr10
Chr09
Chr08
Chr07
Chr06
Chr05
Chr04
Chr03
Chr02
Chr01
Chromosome
0.0e+00
5.0e+06
1.0e+07
1.5e+07
2.0e+07
Position
SatDNA Family
BmorSat18−298

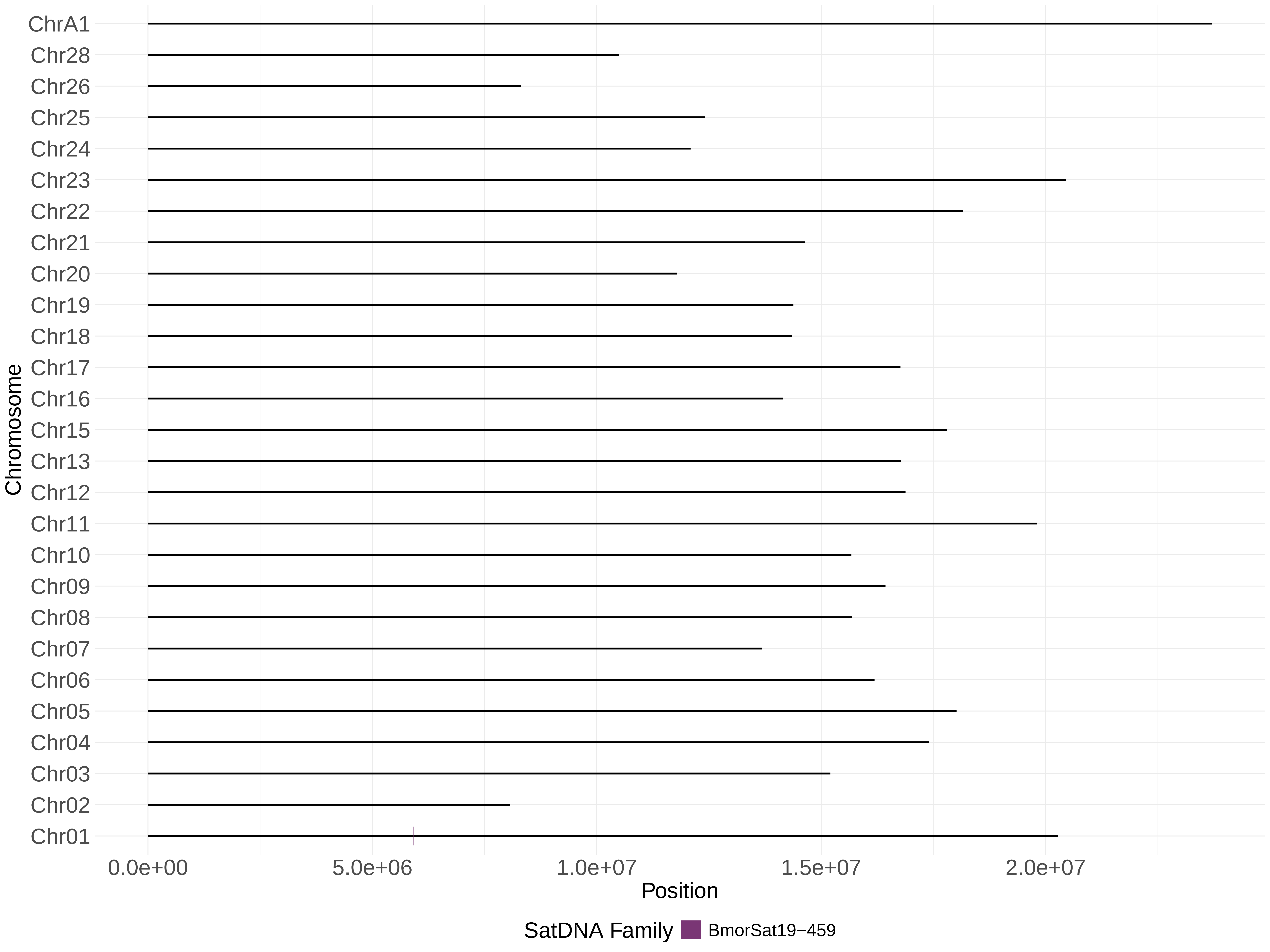
ChrA1
Chr28
Chr26
Chr25
Chr24
Chr23
Chr22
Chr21
Chr20
Chr19
Chr18
Chr17
Chr16
Chr15
Chr13
Chr12
Chr11
Chr10
Chr09
Chr08
Chr07
Chr06
Chr05
Chr04
Chr03
Chr02
Chr01
Chromosome
0.0e+00
5.0e+06
1.0e+07
1.5e+07
2.0e+07
Position
SatDNA Family
BmorSat19−459

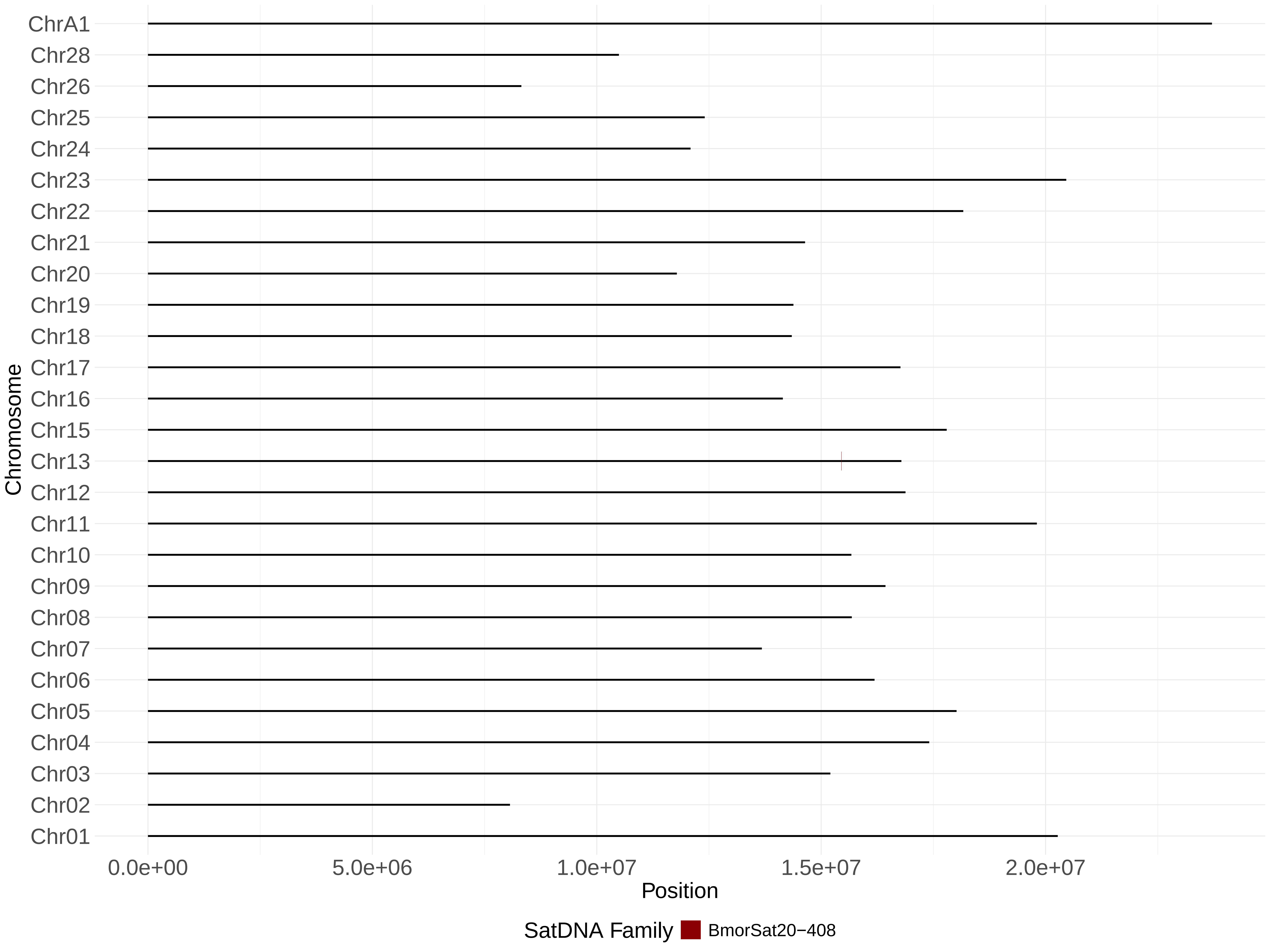
ChrA1
Chr28
Chr26
Chr25
Chr24
Chr23
Chr22
Chr21
Chr20
Chr19
Chr18
Chr17
Chr16
Chr15
Chr13
Chr12
Chr11
Chr10
Chr09
Chr08
Chr07
Chr06
Chr05
Chr04
Chr03
Chr02
Chr01
Chromosome
0.0e+00
5.0e+06
1.0e+07
1.5e+07
2.0e+07
Position
SatDNA Family
BmorSat20−408

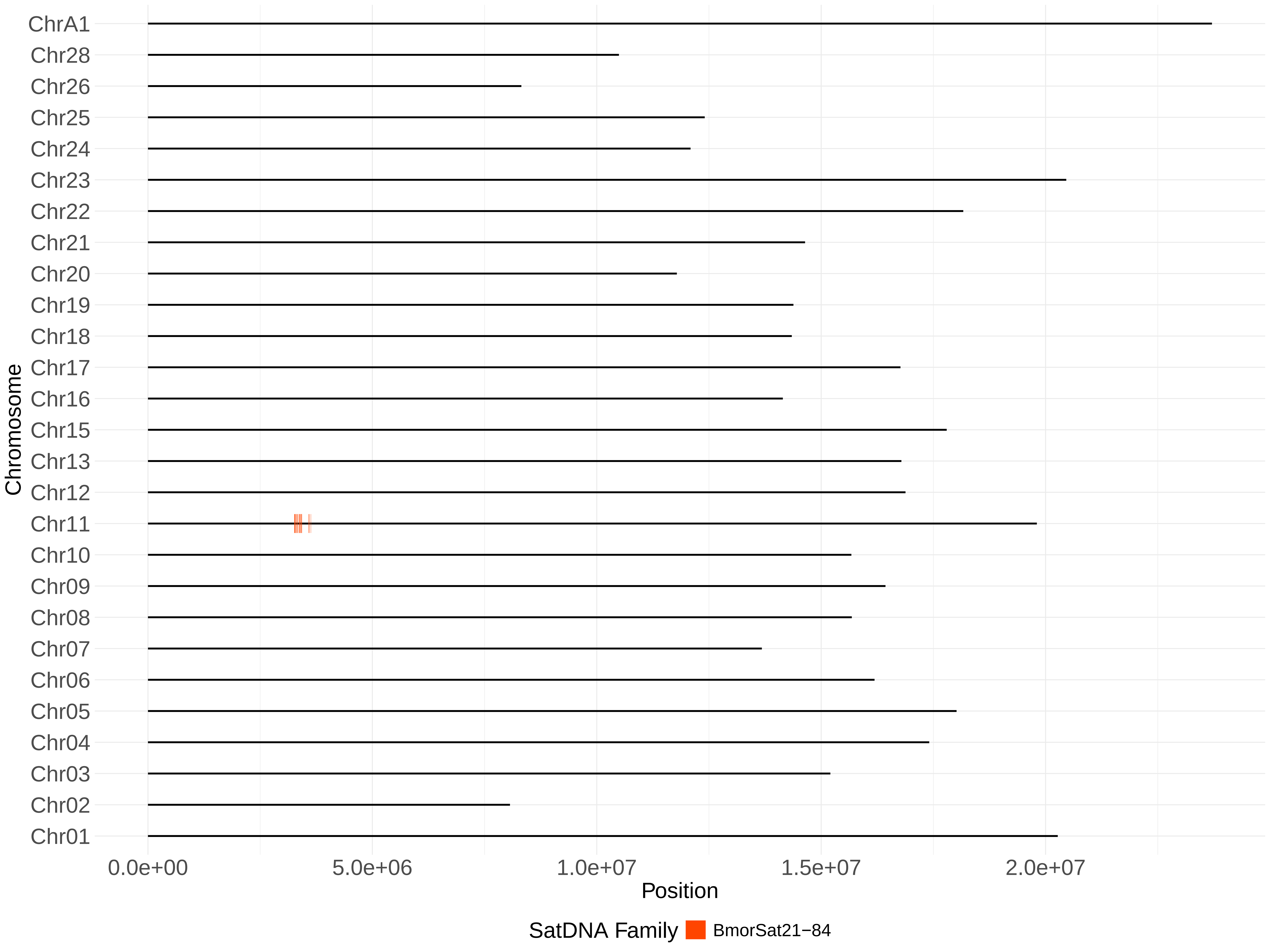
ChrA1
Chr28
Chr26
Chr25
Chr24
Chr23
Chr22
Chr21
Chr20
Chr19
Chr18
Chr17
Chr16
Chr15
Chr13
Chr12
Chr11
Chr10
Chr09
Chr08
Chr07
Chr06
Chr05
Chr04
Chr03
Chr02
Chr01
Chromosome
0.0e+00
5.0e+06
1.0e+07
1.5e+07
2.0e+07
Position
SatDNA Family
BmorSat21−84

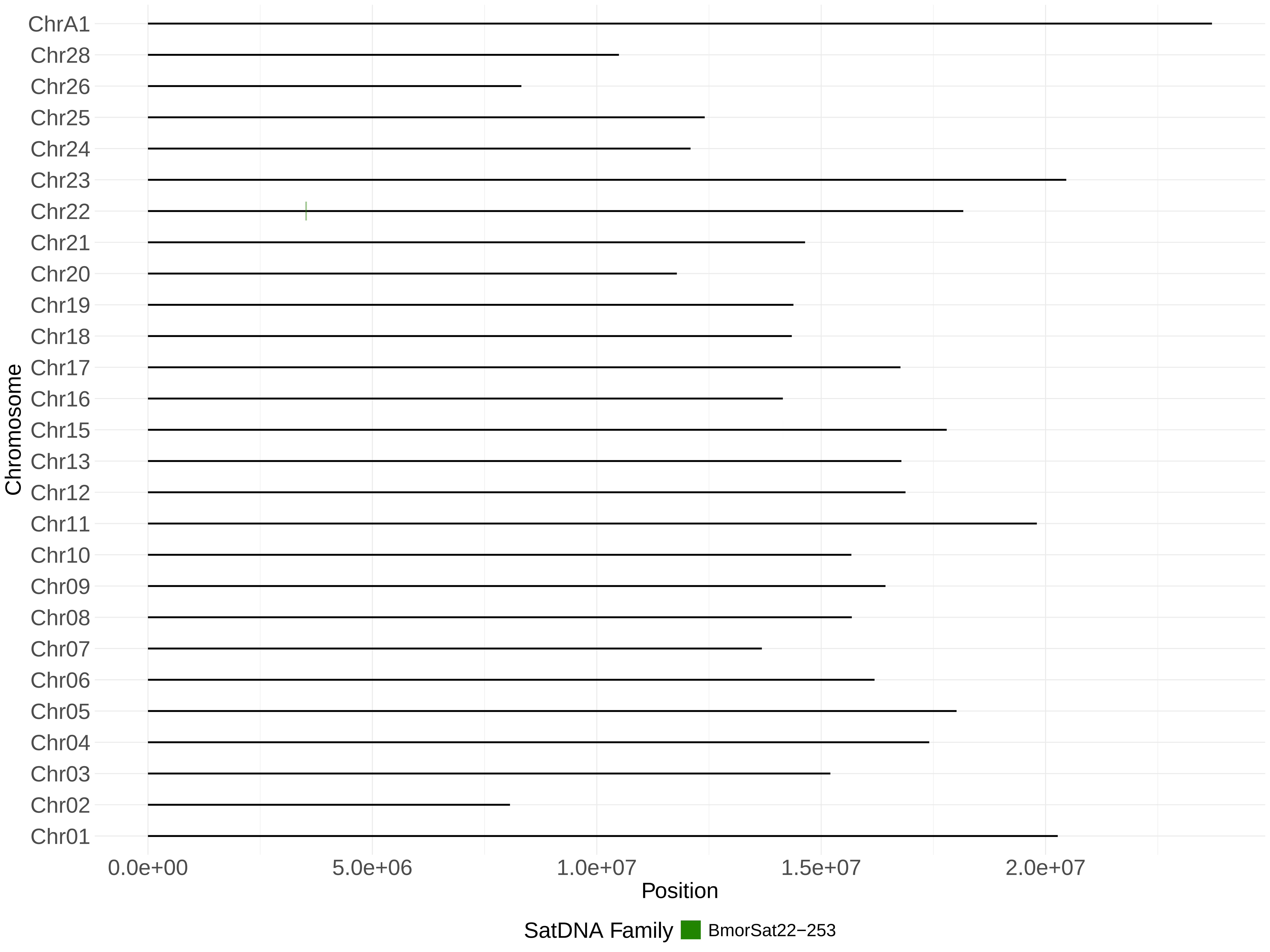
ChrA1
Chr28
Chr26
Chr25
Chr24
Chr23
Chr22
Chr21
Chr20
Chr19
Chr18
Chr17
Chr16
Chr15
Chr13
Chr12
Chr11
Chr10
Chr09
Chr08
Chr07
Chr06
Chr05
Chr04
Chr03
Chr02
Chr01
Chromosome
0.0e+00
5.0e+06
1.0e+07
1.5e+07
2.0e+07
Position
SatDNA Family
BmorSat22−253

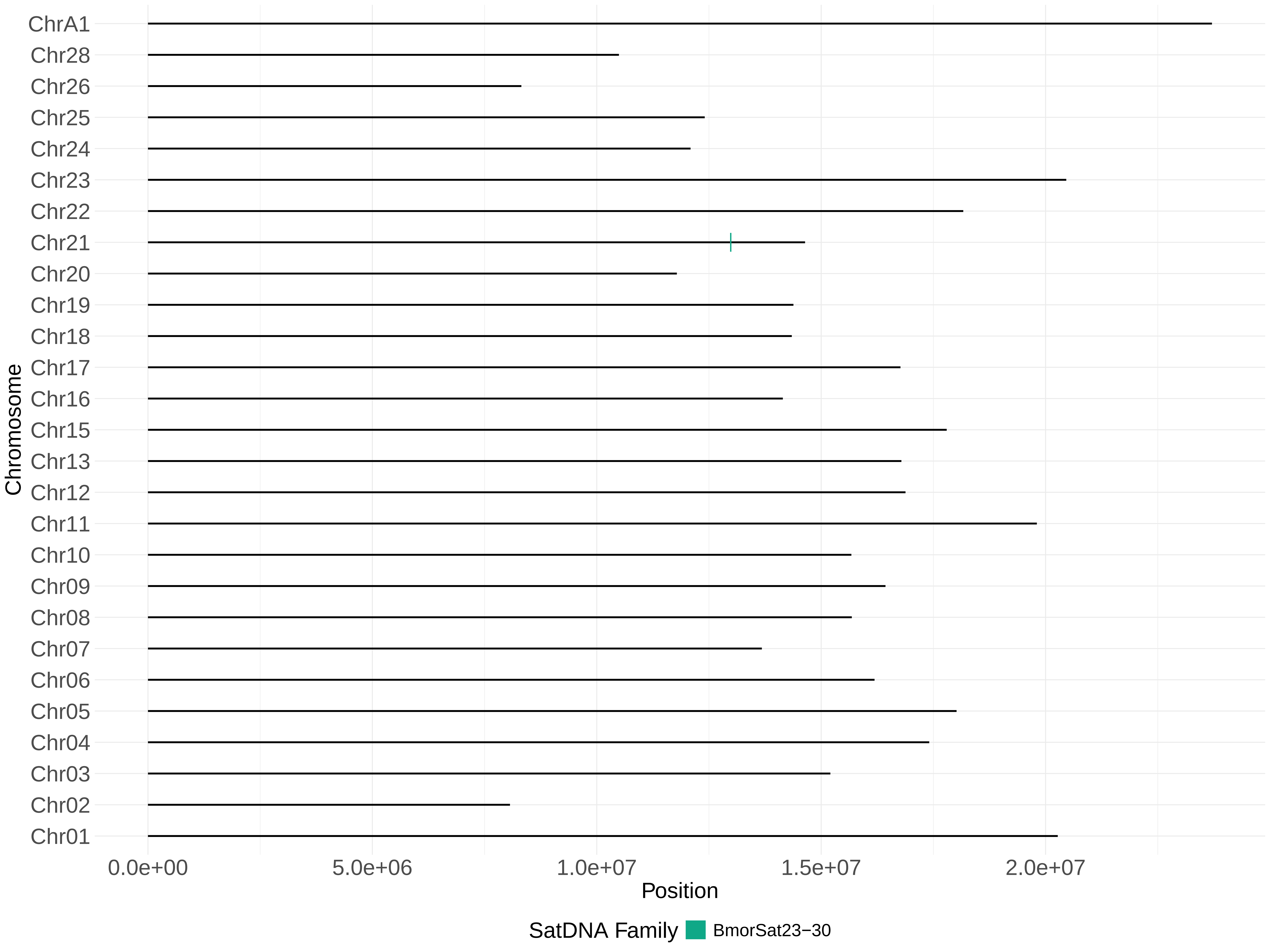
ChrA1
Chr28
Chr26
Chr25
Chr24
Chr23
Chr22
Chr21
Chr20
Chr19
Chr18
Chr17
Chr16
Chr15
Chr13
Chr12
Chr11
Chr10
Chr09
Chr08
Chr07
Chr06
Chr05
Chr04
Chr03
Chr02
Chr01
Chromosome
0.0e+00
5.0e+06
1.0e+07
1.5e+07
2.0e+07
Position
SatDNA Family
BmorSat23−30

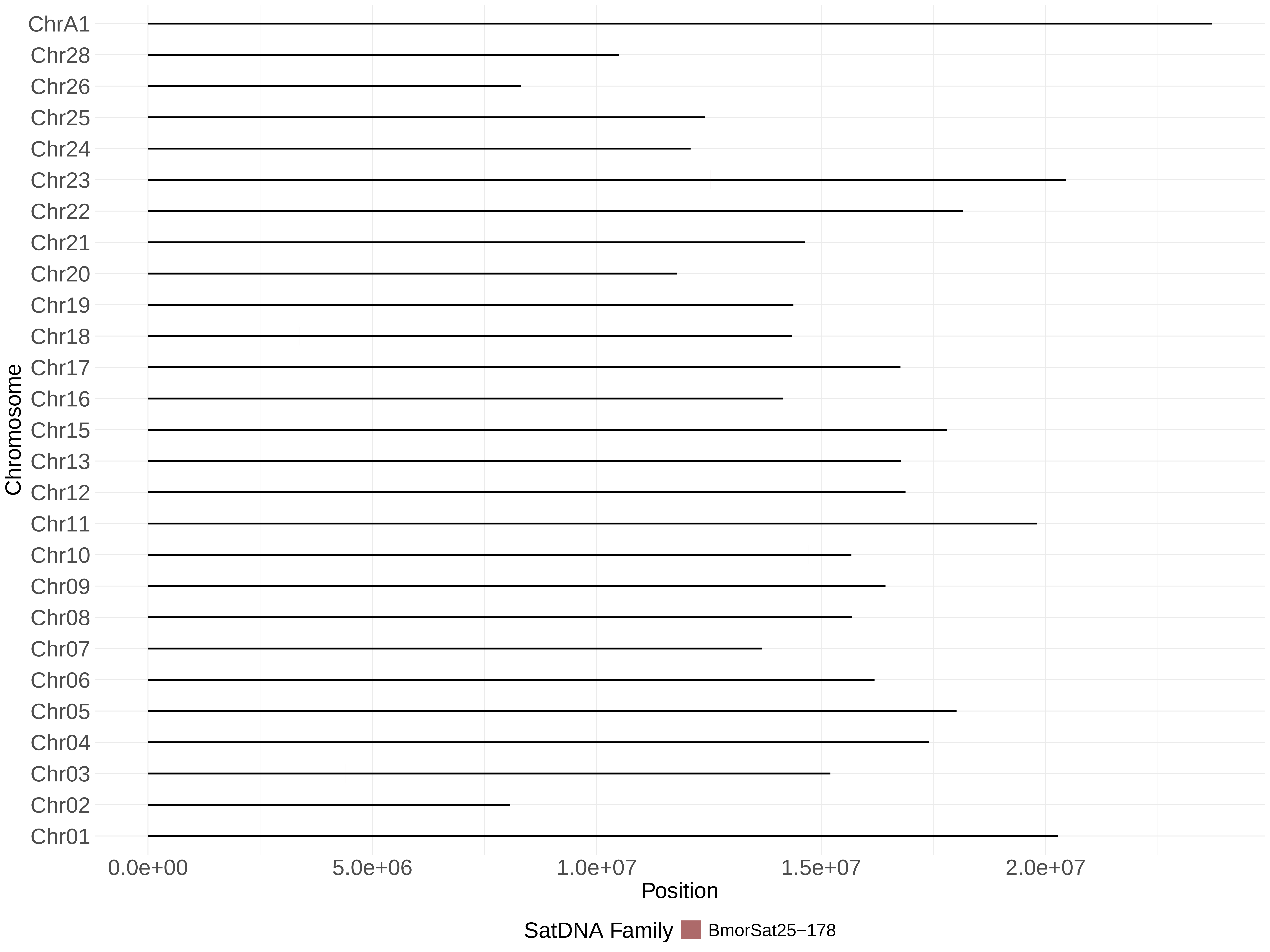
ChrA1
Chr28
Chr26
Chr25
Chr24
Chr23
Chr22
Chr21
Chr20
Chr19
Chr18
Chr17
Chr16
Chr15
Chr13
Chr12
Chr11
Chr10
Chr09
Chr08
Chr07
Chr06
Chr05
Chr04
Chr03
Chr02
Chr01
Chromosome
0.0e+00
5.0e+06
1.0e+07
1.5e+07
2.0e+07
Position
SatDNA Family
BmorSat25−178

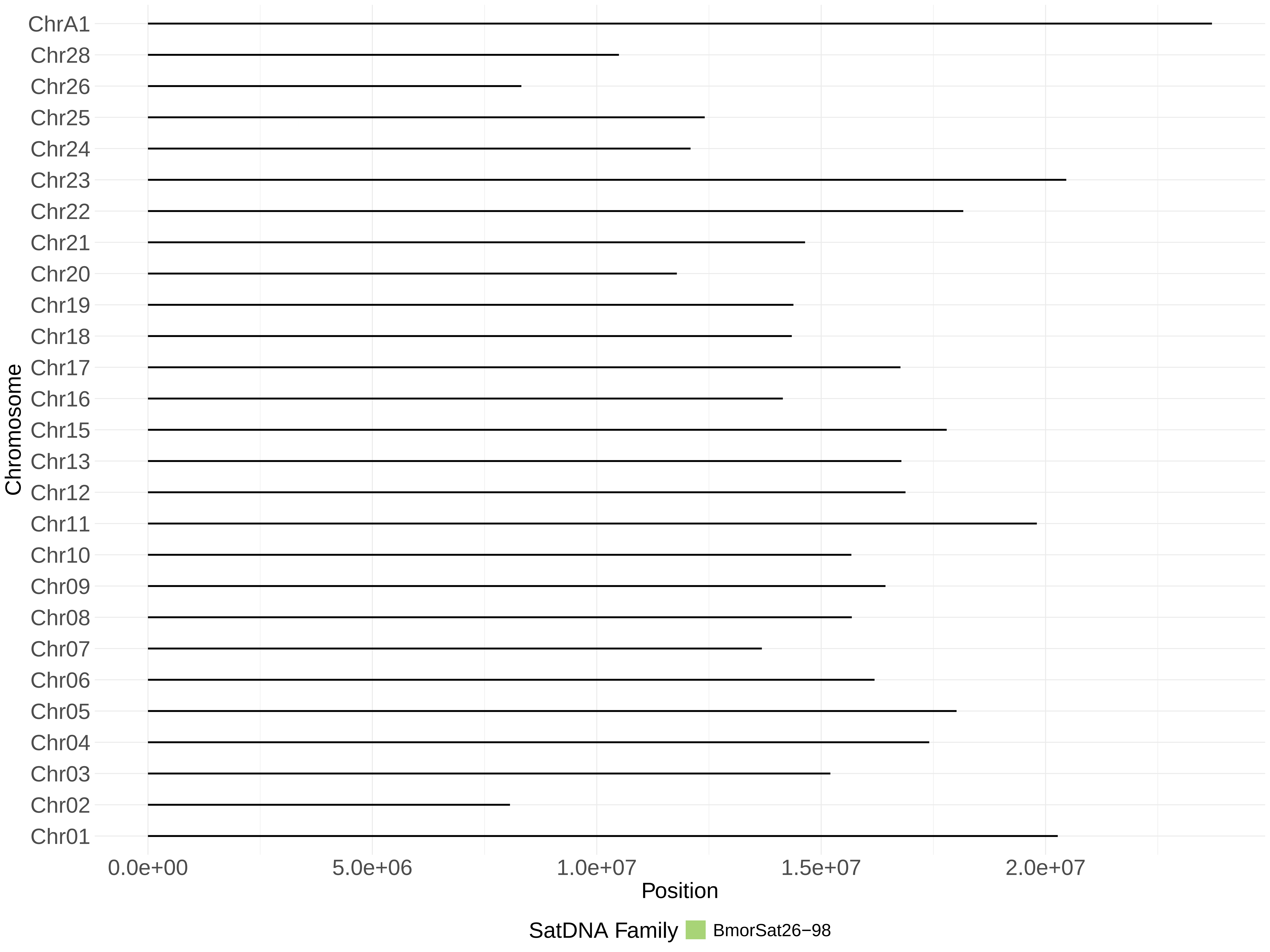

ChrA1
Chr28
Chr26
Chr25
Chr24
Chr23
Chr22
Chr21
Chr20
Chr19
Chr18
Chr17
Chr16
Chr15
Chr13
Chr12
Chr11
Chr10
Chr09
Chr08
Chr07
Chr06
Chr05
Chr04
Chr03
Chr02
Chr01
Chromosome
0.0e+00
5.0e+06
1.0e+07
1.5e+07
2.0e+07
Position
SatDNA Family
BmorSat26−98

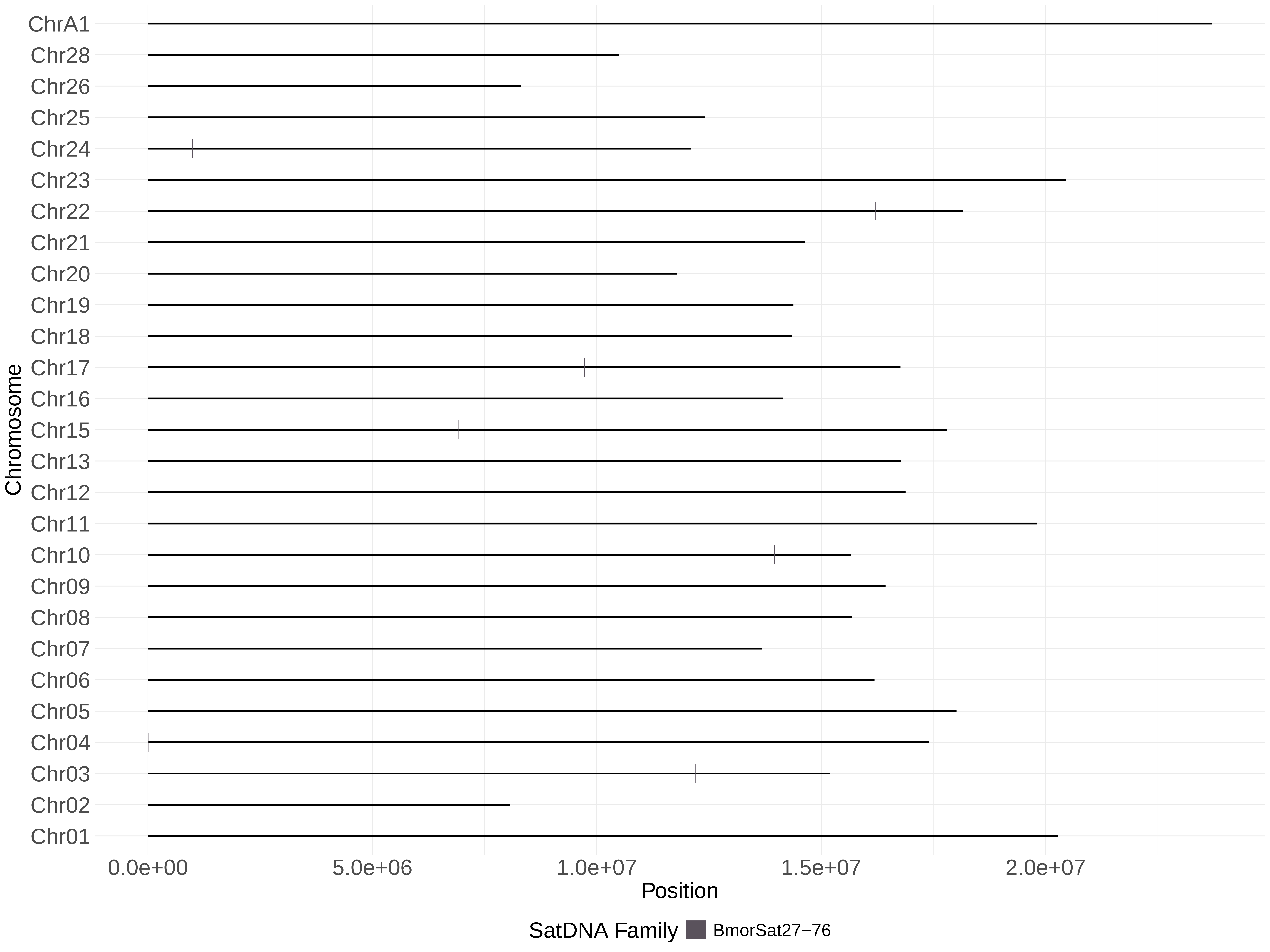

ChrA1
Chr28
Chr26
Chr25
Chr24
Chr23
Chr22
Chr21
Chr20
Chr19
Chr18
Chr17
Chr16
Chr15
Chr13
Chr12
Chr11
Chr10
Chr09
Chr08
Chr07
Chr06
Chr05
Chr04
Chr03
Chr02
Chr01
Chromosome
0.0e+00
5.0e+06
1.0e+07
1.5e+07
2.0e+07
Position
SatDNA Family
BmorSat27−76

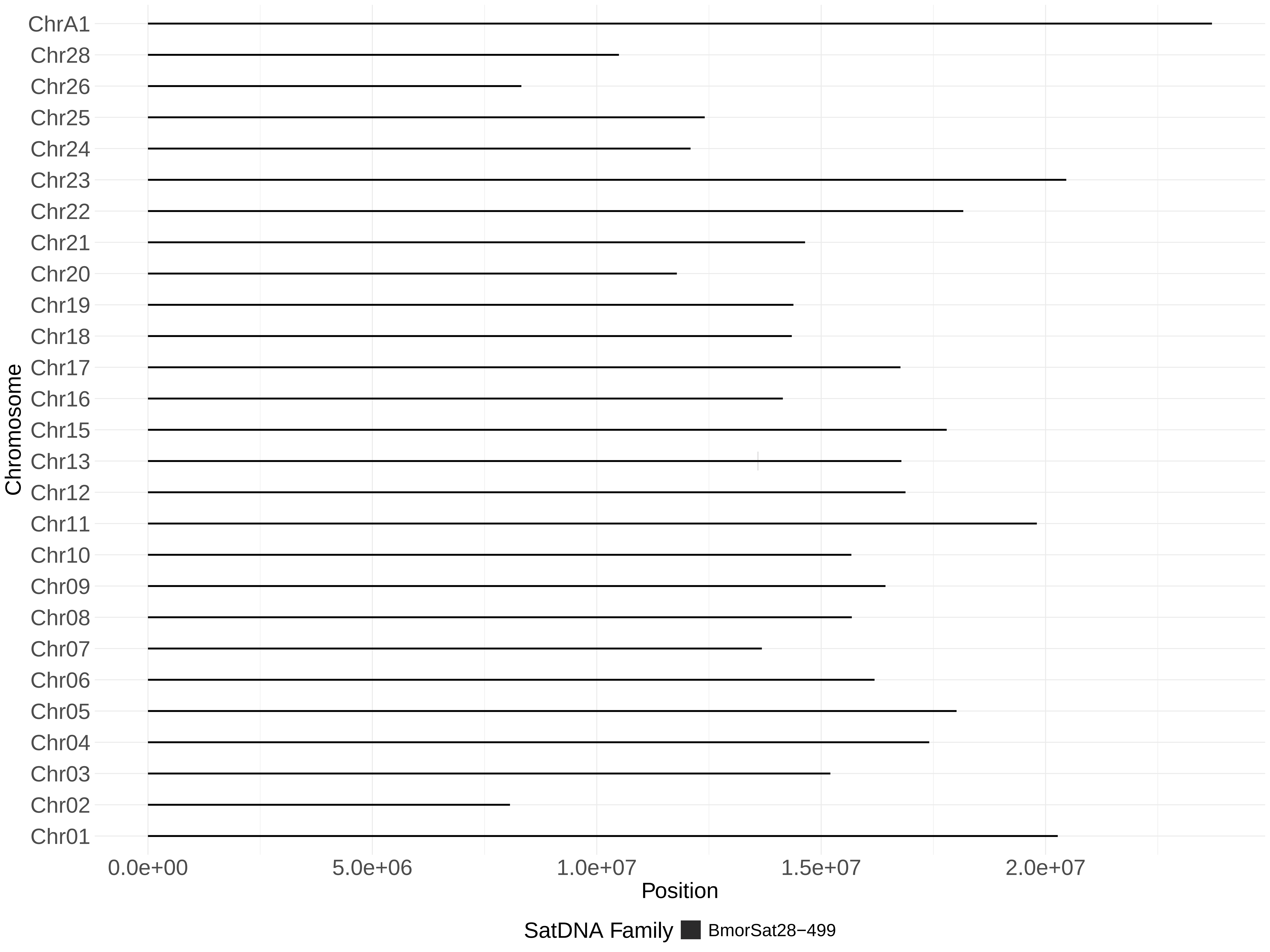
ChrA1
Chr28
Chr26
Chr25
Chr24
Chr23
Chr22
Chr21
Chr20
Chr19
Chr18
Chr17
Chr16
Chr15
Chr13
Chr12
Chr11
Chr10
Chr09
Chr08
Chr07
Chr06
Chr05
Chr04
Chr03
Chr02
Chr01
Chromosome
0.0e+00
5.0e+06
1.0e+07
1.5e+07
2.0e+07
Position
SatDNA Family
BmorSat28−499

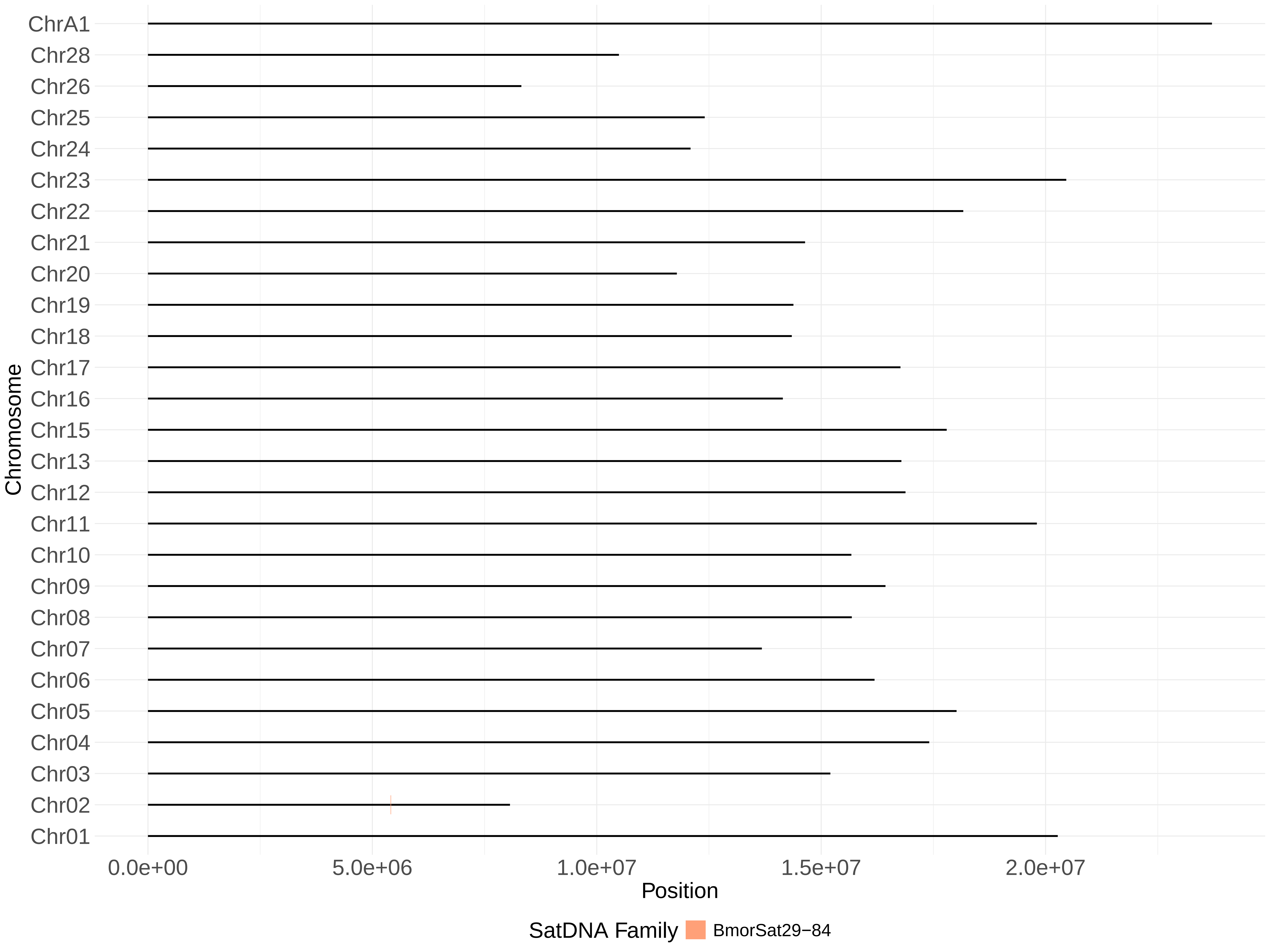

ChrA1
Chr28
Chr26
Chr25
Chr24
Chr23
Chr22
Chr21
Chr20
Chr19
Chr18
Chr17
Chr16
Chr15
Chr13
Chr12
Chr11
Chr10
Chr09
Chr08
Chr07
Chr06
Chr05
Chr04
Chr03
Chr02
Chr01
Chromosome
0.0e+00
5.0e+06
1.0e+07
1.5e+07
2.0e+07
Position
SatDNA Family
BmorSat29−84
