## Supplementary material for "Advancing the understanding of the *Bombyx mori* genome through satellite DNA analysis: low strain differentiation and relationship with transposable elements": Supp table 1

| **SatDNA family** | **Oligonucleotide 5´-3´ (probe)** |
| --- | --- |
| BmorSat01-575 | GCGATGCTGGCTTGCTGTGGGACGACTGTCAAT |
| BmorSat02-1132 | AATGAAGTAGGAAACCTATGGAAATGAAACGTC |
| BmorSat03-379 | CCCCATAATTTTCATACCCCGGGCCTATATATG |
| BmorSat04-1098 | GCTGACGTCACCGTCTTTAAGTTCTCGACGATG |
| BmorSat05-124 | TAGGGGTTGATCGTAGAGGGGTGAAAATTTGAG |
| BmorSat06-47 | GTAATGCACACAGTGTACGACTTAACTTTGTAA |
| BmorSat07-212 | TGACCCACCTTGAGACATGAGGTCGATTTCTCA |
| BmorSat08-513 | GGAGGGTGATTTTTGGTGTGCAATCACTTAGGT |
| BmorSat09-105 | GTACGAGAAGAGCTCGGGATCAGTACAGAGCTG |
| BmorSat10-80 | GCATAACGCATTTCGTGTGACATGCTAGTACGA |
| BmorSat11-138 | CGGTCGTGGATGCGGCAGATGCTGTAACAAAAG |
| BmorSat12-34 | GTTACACTTTCTTCCCTGTACAGCGGGAGGAAA |
| BmorSat13-13 | AGCGGCTGGAGCTAGCGGCTGGAGCT |
| BmorSat14-791 | ATAGGGCAAGTGACGGCTGCGATATGAAGTGCA |
| BmorSat15-175 | GATGAAATTTAGTAAGGTGATAGATGGTAACCT |
| BmorSat16-38 | GTAAGAATTTATAGCAAAAAGTGTCGTGACAAC |
| BmorSat17-20 | GAGCCAAGCCGCTGCCTACCGAGCCAAGCCGCT |
| BmorSat18-298 | ACGCGGTCCTCGTCTGATATTGTGCTCAGTACC |
| BmorSat19-459 | TACTTTCTCAGGGGTTTCTTCAACGACCTGAGC |
| BmorSat20-408 | ATGATGGCGGCTCAGATGGTGGTTCTGTAAGTG |
| BmorSat21-84 | CATAAGGTTTTTCTGCAGTATGTATTCTCATGT |
| BmorSat22-253 | GGTAAGTGAACCTATTCCCATCCTATTGCAGTT |
| BmorSat23-30 | GAGAAGGCACCAAGCCCTGTCAAGGACGAT |
| BmorSat24-753 | GTGAATCAGACTAGGAAACAGTTTTCCACAGCC |
| BmorSat25-178 | CGCGCCGTCGACGCGCGTTTGTAGCGATACAGA |
| BmorSat26-98 | GTGTGACGCGCGGTGTGCTGGAGCACAGAGCTG |
| BmorSat27-76 | GAGCGCATCGGCACCCTTTGATGGCTCGGGTCA |
| BmorSat28-499 | GAGCTTAACGCCAAACGTCACCCTAATTTGACA |
| BmorSat29-84 | GGTACAACTACAACAGAGCTACCTACAACAGTC |
