## Supplementary material for "Advancing the understanding of the *Bombyx mori* genome through satellite DNA analysis: low strain differentiation and relationship with transposable elements": Supp table 2

| **SatDNA Family** | ***Bombyx mori* - Europe local** | | | | | ***Bombyx mori* - China local** | | | | | ***Bombyx mori* - CHN-1** | | | | | ***Bombyx mori* - Japan local** | | | | | ***Bombyx mori* - Tropical local** | |  | |  |
| --- | --- | --- | --- | --- | --- | --- | --- | --- | --- | --- | --- | --- | --- | --- | --- | --- | --- | --- | --- | --- | --- | --- | --- | --- | --- |
|  | **Male** | | **Female** | | **Mean abundance sexes (%)** | **Male (BomL2)** | | **Female (BomL3)** | | **Mean abundance sexes (%)** | **Male (BomO129)** | | **Female (BomP130)** | | **Mean abundance sexes (%)** | **Male (BomP23)** | | **Female (BomP24)** | | **Mean abundance sexes (%)** | **Male (BomL122)** | | **Female (BomL201)** | | **Mean abundance sexes (%)** |
|  | **Abundance (%)** | **Divergence (%)** | **Abundance (%)** | **Divergence (%)** |  | **Abundance (%)** | **Divergence (%)** | **Abundance (%)** | **Divergence (%)** |  | **Abundance (%)** | **Divergence (%)** | **Abundance (%)** | **Divergence (%)** |  | **Abundance (%)** | **Divergence (%)** | **Abundance (%)** | **Divergence (%)** |  | **Abundance (%)** | **Divergence (%)** | **Abundance (%)** | **Divergence (%)** |  |
| BmorSat01-575 | 0.21604 | 9.97 | 0.21739 | 9.83 | 0.21671 | 0.21557 | 10.10 | 0.21962 | 10.10 | 0.21759 | 0.20631 | 10.07 | 0.22188 | 10.08 | 0.21409 | 0.21236 | 10.37 | 0.20977 | 10.13 | 0.21106 | 0.21781 | 10.40 | 0.22313 | 9.96 | 0.22046789 |
| BmorSat02-1132 | 0.12596 | 2.94 | 0.13589 | 2.88 | 0.13092 | 0.14943 | 3.24 | 0.14413 | 3.08 | 0.14678 | 0.13628 | 3.25 | 0.14617 | 3.09 | 0.14123 | 0.13495 | 3.20 | 0.13429 | 3.24 | 0.13462 | 0.14689 | 3.44 | 0.14995 | 2.88 | 0.14841958 |
| BmorSat03-379 | 0.12660 | 0.95 | 0.12677 | 0.98 | 0.12669 | 0.12259 | 1.09 | 0.11844 | 1.03 | 0.12051 | 0.13217 | 1.17 | 0.12194 | 1.05 | 0.12706 | 0.11411 | 0.99 | 0.11709 | 1.01 | 0.11560 | 0.10617 | 1.34 | 0.12234 | 0.82 | 0.11425639 |
| BmorSat04-1098 | 0.03951 | 5.59 | 0.04222 | 5.71 | 0.04087 | 0.03417 | 6.08 | 0.04271 | 5.66 | 0.03844 | 0.04540 | 5.71 | 0.04055 | 5.43 | 0.04297 | 0.03341 | 5.66 | 0.03595 | 5.40 | 0.03468 | 0.03792 | 5.93 | 0.04010 | 5.30 | 0.03900985 |
| BmorSat05-124 | 0.03910 | 11.44 | 0.03689 | 11.31 | 0.03800 | 0.05025 | 11.16 | 0.04061 | 11.30 | 0.04543 | 0.04700 | 11.45 | 0.03420 | 12.20 | 0.04060 | 0.04157 | 11.48 | 0.04255 | 11.55 | 0.04206 | 0.03906 | 11.77 | 0.04144 | 11.45 | 0.04024838 |
| BmorSat06-47 | 0.03140 | 3.45 | 0.03347 | 3.49 | 0.03244 | 0.03021 | 3.65 | 0.03269 | 3.43 | 0.03145 | 0.03408 | 3.63 | 0.03248 | 3.65 | 0.03328 | 0.03276 | 3.53 | 0.03379 | 3.45 | 0.03328 | 0.03102 | 3.75 | 0.03087 | 3.43 | 0.03094573 |
| BmorSat07-212 | 0.02838 | 15.23 | 0.02720 | 15.11 | 0.02779 | 0.02769 | 13.68 | 0.02773 | 14.71 | 0.02771 | 0.02699 | 15.04 | 0.03194 | 13.73 | 0.02946 | 0.03275 | 13.92 | 0.02959 | 14.41 | 0.03117 | 0.02719 | 14.19 | 0.03168 | 13.98 | 0.02943872 |
| BmorSat08-513 | 0.03070 | 4.57 | 0.02244 | 3.40 | 0.02657 | 0.02255 | 5.59 | 0.02057 | 5.66 | 0.02156 | 0.02465 | 4.74 | 0.01704 | 4.44 | 0.02085 | 0.01475 | 4.22 | 0.01724 | 4.25 | 0.01600 | 0.01781 | 5.61 | 0.00817 | 4.36 | 0.01298763 |
| BmorSat09-105 | 0.01256 | 7.08 | 0.01361 | 6.76 | 0.01308 | 0.01092 | 7.88 | 0.00964 | 7.27 | 0.01028 | 0.01100 | 8.05 | 0.01049 | 7.73 | 0.01074 | 0.01067 | 7.55 | 0.01045 | 8.13 | 0.01056 | 0.01105 | 8.52 | 0.01068 | 6.86 | 0.01086482 |
| BmorSat10-80 | 0.01186 | 2.57 | 0.01320 | 2.57 | 0.01253 | 0.01308 | 1.27 | 0.01212 | 1.04 | 0.01260 | 0.01240 | 1.50 | 0.01193 | 1.14 | 0.01217 | 0.01099 | 1.20 | 0.01135 | 1.25 | 0.01117 | 0.01094 | 1.50 | 0.01124 | 0.87 | 0.01108986 |
| BmorSat11-138 | 0.00946 | 6.15 | 0.01022 | 6.21 | 0.00984 | 0.00983 | 6.69 | 0.01167 | 6.79 | 0.01075 | 0.00968 | 6.74 | 0.01230 | 6.82 | 0.01099 | 0.00898 | 6.66 | 0.00811 | 6.41 | 0.00854 | 0.00389 | 6.27 | 0.01043 | 6.53 | 0.00715603 |
| BmorSat12-34 | 0.00914 | 9.89 | 0.00867 | 9.38 | 0.00890 | 0.00914 | 10.16 | 0.00886 | 9.67 | 0.00900 | 0.00849 | 9.83 | 0.00830 | 9.51 | 0.00840 | 0.00919 | 9.62 | 0.00931 | 9.22 | 0.00925 | 0.00910 | 9.29 | 0.00818 | 9.43 | 0.00863862 |
| BmorSat13-13 | 0.00758 | 14.68 | 0.00901 | 14.94 | 0.00829 | 0.00511 | 14.57 | 0.00471 | 15.01 | 0.00491 | 0.00442 | 15.95 | 0.00438 | 15.54 | 0.00440 | 0.00456 | 14.99 | 0.00485 | 15.73 | 0.00470 | 0.00711 | 15.81 | 0.00444 | 14.93 | 0.00577411 |
| BmorSat14-791 | 0.00778 | 1.02 | 0.00809 | 1.39 | 0.00794 | 0.00078 | 13.21 | 0.00075 | 13.19 | 0.00077 | 0.00723 | 2.99 | 0.00652 | 3.39 | 0.00687 | 0.01259 | 1.21 | 0.01260 | 1.22 | 0.01260 | 0.00066 | 13.83 | 0.00820 | 1.58 | 0.0044291 |
| BmorSat15-175 | 0.00700 | 15.23 | 0.00638 | 16.06 | 0.00669 | 0.00754 | 15.93 | 0.00755 | 15.63 | 0.00754 | 0.00722 | 16.07 | 0.00716 | 15.91 | 0.00719 | 0.00627 | 15.59 | 0.00621 | 16.30 | 0.00624 | 0.00628 | 15.77 | 0.00725 | 14.90 | 0.0067641 |
| BmorSat16-38 | 0.00590 | 1.68 | 0.00647 | 1.86 | 0.00619 | 0.00676 | 1.84 | 0.00615 | 1.97 | 0.00645 | 0.00627 | 2.34 | 0.00648 | 2.24 | 0.00637 | 0.00599 | 1.56 | 0.00632 | 1.79 | 0.00615 | 0.00576 | 2.09 | 0.00650 | 1.61 | 0.00612855 |
| BmorSat17-20 | 0.00538 | 6.34 | 0.00583 | 5.83 | 0.00560 | 0.00588 | 6.38 | 0.00560 | 6.26 | 0.00574 | 0.00584 | 6.19 | 0.00447 | 6.65 | 0.00515 | 0.00527 | 6.83 | 0.00568 | 5.80 | 0.00548 | 0.00560 | 5.86 | 0.00620 | 5.61 | 0.00590147 |
| BmorSat18-298 | 0.00699 | 7.49 | 0.00387 | 6.72 | 0.00543 | 0.00863 | 7.22 | 0.00263 | 7.30 | 0.00563 | 0.00697 | 6.28 | 0.00470 | 6.11 | 0.00583 | 0.00664 | 6.87 | 0.00283 | 7.82 | 0.00473 | 0.00775 | 7.66 | 0.00394 | 5.57 | 0.00584564 |
| BmorSat19-459 | 0.00707 | 5.57 | 0.00358 | 6.67 | 0.00532 | 0.00993 | 5.17 | 0.00349 | 4.59 | 0.00671 | 0.00925 | 5.46 | 0.00386 | 5.41 | 0.00656 | 0.00610 | 5.33 | 0.00439 | 4.75 | 0.00525 | 0.00736 | 5.82 | 0.00372 | 5.39 | 0.00554144 |
| BmorSat20-408 | 0.00451 | 3.21 | 0.00481 | 2.78 | 0.00466 | 0.00351 | 3.52 | 0.00397 | 3.11 | 0.00374 | 0.00618 | 3.21 | 0.00499 | 2.94 | 0.00559 | 0.00571 | 2.85 | 0.00476 | 2.90 | 0.00524 | 0.00584 | 4.79 | 0.00577 | 2.63 | 0.00580508 |
| BmorSat21-84 | 0.00465 | 27.16 | 0.00450 | 26.12 | 0.00458 | 0.00420 | 25.30 | 0.00462 | 26.09 | 0.00441 | 0.00529 | 26.52 | 0.00535 | 25.82 | 0.00532 | 0.00453 | 26.42 | 0.00438 | 26.04 | 0.00445 | 0.00430 | 26.40 | 0.00413 | 26.11 | 0.00421313 |
| BmorSat22-253 | 0.00374 | 9.79 | 0.00487 | 8.96 | 0.00430 | 0.00392 | 10.30 | 0.00529 | 9.67 | 0.00460 | 0.00446 | 9.79 | 0.00472 | 9.69 | 0.00459 | 0.00455 | 9.50 | 0.00461 | 9.47 | 0.00458 | 0.00462 | 9.67 | 0.00445 | 9.57 | 0.00453384 |
| BmorSat23-30 | 0.00433 | 23.72 | 0.00403 | 23.92 | 0.00418 | 0.00615 | 23.38 | 0.00450 | 21.73 | 0.00533 | 0.00624 | 23.08 | 0.00508 | 22.83 | 0.00566 | 0.00424 | 23.80 | 0.00495 | 23.39 | 0.00459 | 0.00388 | 23.57 | 0.00541 | 23.77 | 0.00464444 |
| BmorSat24-753 | 0.00001 | 22.53 | 0.00563 | 2.41 | 0.00282 | 0.00004 | 24.75 | 0.00860 | 2.07 | 0.00432 | 0.00000 | 0.00000 | 0.00744 | 1.91 | 0.00372 | 0.00001 | 19.90 | 0.00660 | 2.10 | 0.00331 | 0.00000 | 0.0 | 0.00622 | 1.95 | 0.00311037 |
| BmorSat25-178 | 0.00239 | 7.70 | 0.00295 | 8.29 | 0.00267 | 0.00292 | 7.95 | 0.00248 | 8.87 | 0.00270 | 0.00258 | 8.33 | 0.00322 | 9.08 | 0.00290 | 0.00245 | 8.11 | 0.00295 | 8.19 | 0.00270 | 0.00288 | 8.14 | 0.00265 | 7.54 | 0.00276629 |
| BmorSat26-98 | 0.00255 | 5.41 | 0.00247 | 5.84 | 0.00251 | 0.00358 | 3.77 | 0.00335 | 3.49 | 0.00347 | 0.00315 | 3.77 | 0.00288 | 5.46 | 0.00302 | 0.00355 | 3.81 | 0.00350 | 3.71 | 0.00352 | 0.00282 | 4.13 | 0.00217 | 3.69 | 0.00249253 |
| BmorSat27-76 | 0.00208 | 13.59 | 0.00228 | 13.75 | 0.00218 | 0.00338 | 13.57 | 0.00242 | 14.96 | 0.00290 | 0.00173 | 14.59 | 0.00254 | 13.84 | 0.00214 | 0.00221 | 13.72 | 0.00242 | 14.18 | 0.00231 | 0.00239 | 14.68 | 0.00145 | 13.89 | 0.00191843 |
| BmorSat28-499 | 0.00154 | 2.13 | 0.00251 | 1.70 | 0.00203 | 0.00172 | 3.39 | 0.00160 | 2.58 | 0.00166 | 0.00170 | 3.21 | 0.00151 | 1.48 | 0.00160 | 0.00339 | 1.30 | 0.00282 | 1.28 | 0.00311 | 0.00281 | 2.12 | 0.00169 | 2.29 | 0.00225087 |
| BmorSat29-84 | 0.00178 | 10.90 | 0.00164 | 8.36 | 0.00171 | 0.00127 | 12.40 | 0.00173 | 11.76 | 0.00150 | 0.00209 | 10.81 | 0.00114 | 12.58 | 0.00162 | 0.00165 | 10.79 | 0.00152 | 12.29 | 0.00159 | 0.00145 | 12.93 | 0.00163 | 9.58 | 0.00154056 |
| **Total Abundance** | 0.75602 |  | 0.76690 |  | 0.76146 | 0.77072 |  | 0.75825 |  | 0.76449 | 0.77505 |  | 0.76569 |  | 0.77037 | 0.73618 |  | 0.74091 |  |  | 0.73034 |  | 0.76403 |  |  |
