## Supplementary material for "Advancing the understanding of the *Bombyx mori* genome through satellite DNA analysis: low strain differentiation and relationship with transposable elements": Supp table 3

| **SatDNA family** | ***Bombyx mandarina*** | | | | |
| --- | --- | --- | --- | --- | --- |
|  | **Male** | | **Female** | | **Mean abundance sexes (%)** |
|  | **Abundance (%)** | **Divergence (%)** | **Abundance (%)** | **Divergence (%)** |  |
| BmorSat01-575 | 0.2233132 | 11.04 | 0.227573556 | 9.63 | 0.225443378 |
| BmorSat02-1132 | 0.171689333 | 3.72 | 0.145172444 | 2.78 | 0.158430889 |
| BmorSat03-379 | 0.079350667 | 1.79 | 0.135199111 | 0.98 | 0.107274889 |
| BmorSat04-1098 | 0.0937048 | 9.44 | 0.040028444 | 5.63 | 0.066866622 |
| BmorSat05-124 | 0.0352208 | 11.05 | 0.031328222 | 11.62 | 0.033274511 |
| BmorSat06-47 | 0.034705733 | 3.40 | 0.031248889 | 3.51 | 0.032977311 |
| BmorSat07-212 | 0.016369733 | 26.77 | 0.021760444 | 17.81 | 0.019065089 |
| BmorSat08-513 | 0.032715733 | 8.95 | 0.04376 | 4.30 | 0.038237867 |
| BmorSat09-105 | 0.014806 | 6.63 | 0.013954444 | 7.56 | 0.014380222 |
| BmorSat10-80 | 0.013702533 | 1.09 | 0.011046222 | 1.48 | 0.012374378 |
| BmorSat11-138 | 0.010617333 | 7.77 | 0.013148444 | 6.54 | 0.011882889 |
| BmorSat12-34 | 0.010497467 | 10.59 | 0.008214667 | 9.54 | 0.009356067 |
| BmorSat13-13 | 0.001834933 | 16.06 | 0.008918889 | 14.62 | 0.005376911 |
| BmorSat14-791 | 0.004219067 | 5.67 | 0.002804444 | 5.08 | 0.003511756 |
| BmorSat15-175 | 0.004568667 | 19.42 | 0.005801111 | 15.40 | 0.005184889 |
| BmorSat16-38 | 0.0046228 | 2.78 | 0.006843333 | 1.81 | 0.005733067 |
| BmorSat17-20 | 0.005727333 | 6.19 | 0.005895333 | 5.76 | 0.005811333 |
| BmorSat18-298 | 0.006652267 | 10.31 | 0.005717333 | 6.35 | 0.0061848 |
| BmorSat19-459 | 0.0072912 | 5.98 | 0.003858222 | 5.85 | 0.005574711 |
| BmorSat20-408 | 0.0033184 | 3.20 | 0.003416889 | 2.44 | 0.003367644 |
| BmorSat21-84 | 0.005180667 | 26.89 | 0.004791556 | 25.99 | 0.004986111 |
| BmorSat22-253 | 0.006124933 | 10.05 | 0.002723333 | 10.49 | 0.004424133 |
| BmorSat23-30 | 0.004602267 | 23.79 | 0.005533111 | 21.61 | 0.005067689 |
| BmorSat24-753 | 0.0000228 | 22.53 | 0.008856444 | 1.76 | 0.004439622 |
| BmorSat25-178 | 0.0021852 | 7.35 | 0.002512667 | 8.28 | 0.002348933 |
| BmorSat26-98 | 0.003765067 | 4.87 | 0.005284667 | 6.28 | 0.004524867 |
| BmorSat27-76 | 0.001654667 | 13.55 | 0.002770222 | 13.50 | 0.002212444 |
| BmorSat28-499 | 0.0049956 | 4.26 | 9.77778E-05 | 6.41 | 0.002546689 |
| BmorSat29-84 | 0.0017416 | 11.73 | 0.002009111 | 13.34 | 0.001875356 |
| **Total abundance** | 0.8052008 |  | 0.800269333 |  | 0.802735067 |
