## Supplementary material for "Advancing the understanding of the *Bombyx mori* genome through satellite DNA analysis: low strain differentiation and relationship with transposable elements": Supp table 4

| **SatDNA family** | ***Bombyx mori*** | | | | | | ***Bombyx mandarina*** | | | |
| --- | --- | --- | --- | --- | --- | --- | --- | --- | --- | --- |
|  | **Autosomes** | | **ChrW** | | **ChrZ** | | **Autosomes** | | **ChrZ** | |
|  | **Abundance (%)** | **Divergence (%)** | **Abundance (%)** | **Divergence (%)** | **Abundance (%)** | **Divergence (%)** | **Abundance (%)** | **Divergence (%)** | **Abundance (%)** | **Divergence (%)** |
| **BmorSat01-575** | 0.2440 | 11.58 | 0.2594 | 3.32 | 0.1601 | 12.13 | 0.2586 | 11.35 | 0.1330 | 13.17 |
| **BmorSat02-1132** | 0.1467 | 2.86 | 0.2639 | 4.82 | 0.1091 | 3.62 | 0.1548 | 3.65 | 0.1416 | 4.35 |
| **BmorSat03-379** | 0.1274 | 1.20 | 0.1348 | 1.36 | 0.0767 | 1.05 | 0.0807 | 1.39 | 0.0455 | 1.25 |
| **BmorSat04-1098** | 0.0371 | 6.55 | 0.4505 | 10.86 | 0.0083 | 12.14 | 0.1059 | 9.36 | 0.0446 | 10.10 |
| **BmorSat05-124** | 0.0455 | 11.01 | 0.0000 | 0 | 0.0518 | 12.02 | 0.0367 | 10.43 | 0.0546 | 10.16 |
| **BmorSat06-47** | 0.0364 | 3.32 | 0.0000 | 0 | 0.0183 | 3.19 | 0.0370 | 3.27 | 0.0177 | 3.48 |
| **BmorSat07-212** | 0.0312 | 13.86 | 0.0000 | 0 | 0.0188 | 12.50 | 0.0121 | 25.88 | 0.0093 | 25.78 |
| **BmorSat08-513** | 0.0186 | 3.99 | 0.2397 | 5.21 | 0.0000 | 0 | 0.0017 | 7.26 | 0.0000 | 0.0000 |
| **BmorSat09-105** | 0.0143 | 6.19 | 0.0000 | 0 | 0.0000 | 0 | 0.0147 | 6.24 | 0.0000 | 0.0000 |
| **BmorSat10-80** | 0.0100 | 0.25 | 0.0431 | 0.11 | 0.0026 | 0.26 | 0.0169 | 0.05 | 0.0020 | 0.29 |
| **BmorSat11-138** | 0.0116 | 6.14 | 0.0000 | 0 | 0.0000 | 0 | 0.0000 | 0.0000 | 0.0000 | 0.0000 |
| **BmorSat12-34** | 0.0100 | 9.83 | 0.0000 | 0 | 0.0096 | 10.26 | 0.0095 | 10.81 | 0.0096 | 13.23 |
| **BmorSat13-13** | 0.0093 | 10.51 | 0.0000 | 0 | 0.0000 | 0 | 0.0018 | 12.96 | 0.0000 | 0.0000 |
| **BmorSat14-791** | 0.0008 | 15.26 | 0.0000 | 0 | 0.0000 | 0 | 0.0000 | 0.0000 | 0.0000 | 0.0000 |
| **BmorSat15-175** | 0.0075 | 15.57 | 0.0000 | 0 | 0.0059 | 15.07 | 0.0042 | 19.19 | 0.0023 | 22.35 |
| **BmorSat16-38** | 0.0069 | 2.22 | 0.0000 | 0 | 0.0063 | 4.05 | 0.0050 | 2.56 | 0.0030 | 3.48 |
| **BmorSat17-20** | 0.0071 | 6.04 | 0.0000 | 0 | 0.0043 | 4.82 | 0.0070 | 6.45 | 0.0058 | 5.35 |
| **BmorSat18-298** | 0.0000 | 11.04 | 0.0000 | 0 | 0.1615 | 5.17 | 0.0000 | 0.0000 | 0.1356 | 8.28 |
| **BmorSat19-459** | 0.0000 | 0 | 0.0000 | 0 | 0.1599 | 4.14 | 0.0000 | 0.0000 | 0.0307 | 7.16 |
| **BmorSat20-408** | 0.0066 | 4.07 | 0.0000 | 0 | 0.0000 | 0 | 0.0023 | 3.07 | 0.0000 | 0.0000 |
| **BmorSat21-84** | 0.0060 | 25.55 | 0.0000 | 0 | 0.0000 | 0 | 0.0054 | 25.89 | 0.0000 | 0.0000 |
| **BmorSat22-253** | 0.0033 | 9.23 | 0.0000 | 0 | 0.0000 | 0 | 0.0053 | 8.95 | 0.0000 | 0.0000 |
| **BmorSat23-30** | 0.0051 | 19.45 | 0.0000 | 0 | 0.0000 | 0 | 0.0053 | 20.62 | 0.0000 | 0.0000 |
| **BmorSat24-753** | 0.0000 | 15.85 | 1.0286 | 2.86 | 0.0003 | 20.09 | 0.0000 | 0.0000 | 0.0003 | 22.53 |
| **BmorSat25-178** | 0.0028 | 8.20 | 0.0000 | 0 | 0.0037 | 5.03 | 0.0025 | 8.05 | 0.0021 | 6.51 |
| **BmorSat26-98** | 0.0037 | 2.88 | 0.0000 | 0 | 0.0002 | 5.32 | 0.0002 | 17.63 | 0.0002 | 5.32 |
| **BmorSat27-76** | 0.0025 | 11.80 | 0.0326 | 12.55 | 0.0028 | 12.58 | 0.0021 | 11.98 | 0.0000 | 0.0000 |
| **BmorSat28-499** | 0.0051 | 0.79 | 0.0000 | 0 | 0.0000 | 0 | 0.0012 | 2.04 | 0.0000 | 0.0000 |
| **BmorSat29-84** | 0.0027 | 5.18 | 0.0000 | 0 | 0.0000 | 0 | 0.0022 | 6.60 | 0.0000 | 0.0000 |
